# Concerted neuron-astrocyte gene expression declines in aging and schizophrenia

**DOI:** 10.1101/2024.01.07.574148

**Authors:** Emi Ling, James Nemesh, Melissa Goldman, Nolan Kamitaki, Nora Reed, Robert E. Handsaker, Giulio Genovese, Jonathan S. Vogelgsang, Sherif Gerges, Seva Kashin, Sulagna Ghosh, John M. Esposito, Kiely French, Daniel Meyer, Alyssa Lutservitz, Christopher D. Mullally, Alec Wysoker, Liv Spina, Anna Neumann, Marina Hogan, Kiku Ichihara, Sabina Berretta, Steven A. McCarroll

## Abstract

Human brains vary across people and over time; such variation is not yet understood in cellular terms. Here we describe a striking relationship between people’s cortical neurons and cortical astrocytes. We used single-nucleus RNA-seq to analyze the prefrontal cortex of 191 human donors ages 22-97 years, including healthy individuals and persons with schizophrenia. Latent-factor analysis of these data revealed that in persons whose cortical neurons more strongly expressed genes for synaptic components, cortical astrocytes more strongly expressed distinct genes with synaptic functions and genes for synthesizing cholesterol, an astrocyte-supplied component of synaptic membranes. We call this relationship the Synaptic Neuron-and-Astrocyte Program (SNAP). In schizophrenia and aging – two conditions that involve declines in cognitive flexibility and plasticity ^1,2^ – cells had divested from SNAP: astrocytes, glutamatergic (excitatory) neurons, and GABAergic (inhibitory) neurons all reduced SNAP expression to corresponding degrees. The distinct astrocytic and neuronal components of SNAP both involved genes in which genetic risk factors for schizophrenia were strongly concentrated. SNAP, which varies quantitatively even among healthy persons of similar age, may underlie many aspects of normal human interindividual differences and be an important point of convergence for multiple kinds of pathophysiology.

## INTRODUCTION

In natural, non-laboratory settings – in which individuals have diverse genetic inheritances, environments and life histories, as humans do – almost all aspects of biology exhibit quantitative variation across individuals ^3^. Natural variation makes it possible to observe a biological system across many contexts and potentially learn underlying principles ^4,5^.

Here we sought to recognize changes that multiple cell types in the human brain characteristically implement together. The need to be able to recognize tissue-level gene-expression programs comes from a simple but important idea in the physiology of the brain and other tissues: cells of different types collaborate to perform essential functions, working together to construct and regulate structures such as synaptic networks.

We analyzed the prefrontal cortex of 191 human brain donors by single-nucleus RNA-seq (snRNA-seq) and developed a computational approach, based on latent-factor analysis, to recognize commonly recurring multicellular gene-expression patterns in such data. Tissue-level programs whose expression varies across individuals could provide new ways to understand healthy brain function and also brain disorders, since disease processes likely act through endogenous pathways in cells and tissues. A longstanding challenge in genetically complex brain disorders is to identify the aspects of brain biology on which disparate genetic effects converge; here we applied this idea to try to better understand schizophrenia.

## RESULTS

### Dorsolateral prefrontal cortex snRNA-seq

We analyzed the dorsolateral prefrontal cortex (dlPFC, Brodmann area 46), which serves working memory, attention, executive functions, and cognitive flexibility ^6^, abilities which decline in schizophrenia and with advancing age ^1,2^. Analyses included frozen post-mortem dlPFC from 191 donors (ages 22-97, median 64), including 97 without known psychiatric conditions and 94 affected by schizophrenia **(Extended Data Fig. 1 and Supplementary Table 1)**. To generate data that were well-controlled across donors and thus amenable to integrative analysis, we processed a series of 20-donor sets of dlPFC tissue each as a single pooled sample (or “village” ^7^) **(Fig. 1a)**, then, during computational analysis, used combinations of many transcribed SNPs to identify the source donor of each nucleus **(Fig. 1a-b** and **Extended Data Fig. 2)**.

**Figure 1.**
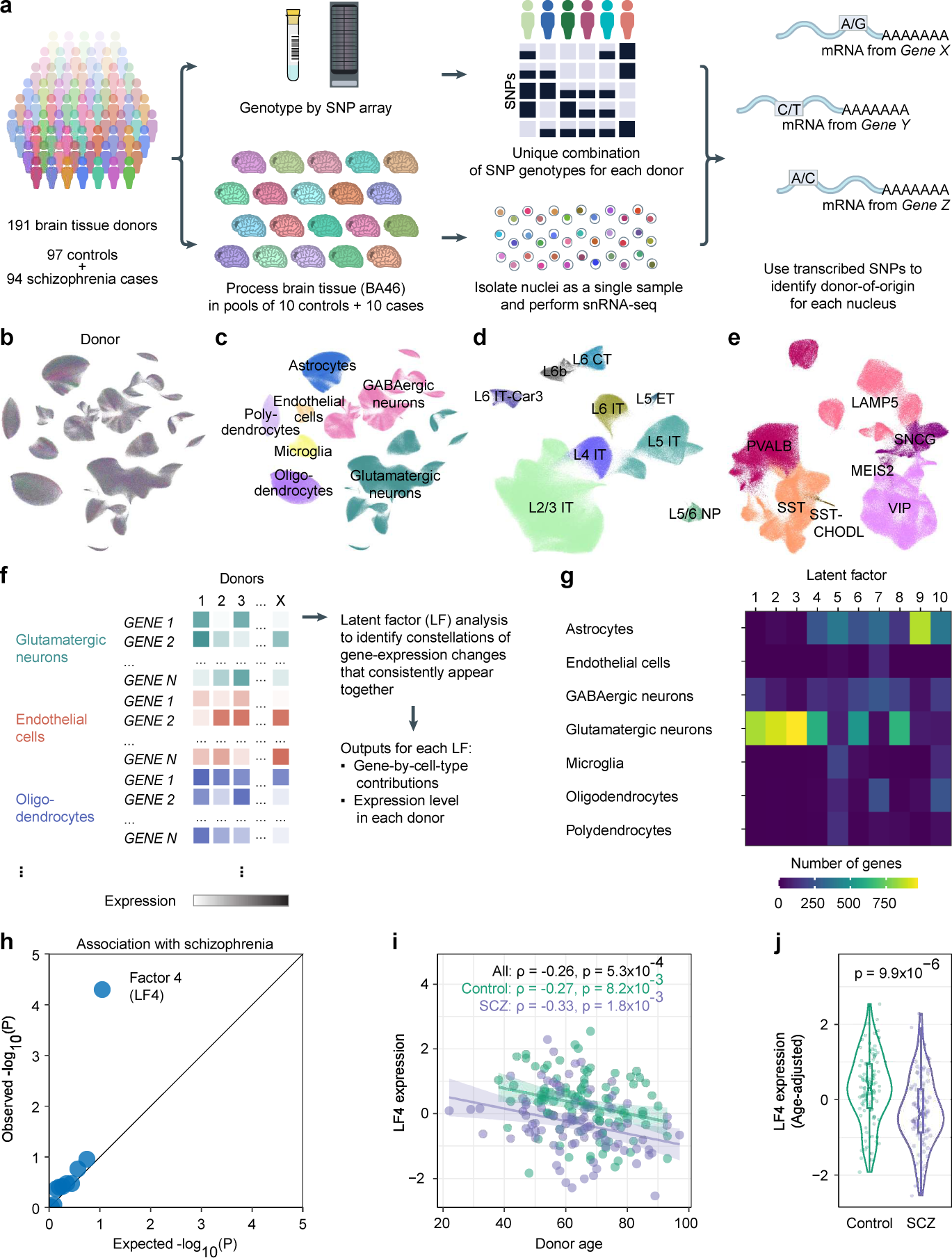
Identification of concerted multi-cellular gene-expression changes common to schizophrenia and aging. **a,** Generation of snRNA-seq data, in a series of 20-donor “villages”. **b,** Uniform manifold approximation and projection (UMAP, colored by donor) of the RNA-expression profiles of the 1,218,284 nuclei analyzed from 191 donors. **c,** Assignments of nuclei to cell types (same projection as in **b**). **d-e,** Assignments of nuclei to **(d)** glutamatergic (*n* = 524,186) and **(e)** GABAergic (*n* = 238,311) neuron subtypes. **f,** Latent factor analysis. Cell-type-resolution expression data from all donors and cell types were combined into a single analysis. Latent factor analysis identified constellations of gene-expression changes that consistently appeared together. **g,** Cell type-specificity of the latent factors inferred from 180 donors, shown as cell-type distributions of the 1,000 most strongly loading gene/cell-type combinations per factor. Factors 4-7 and 10 are strongly driven by gene-expression co-variation spanning multiple cell types. **h,** Association of schizophrenia with inter-individual variation in the expression levels of the ten latent-factors in Fig. 1g, shown as a quantile-quantile plot comparing the ten factors’ observed schizophrenia associations (-log_10_ p-values) to the distribution of association statistics expected by chance; only LF4 significantly associates with schizophrenia. See also Extended Data Fig. 10. **i,** Relationship of quantile-normalized Latent Factor 4 (LF4) donor expression levels to age (Spearman’s ⍴; *n* = 180 donors). Shaded regions represent 95% confidence intervals. **j,** Quantile-normalized LF4 donor scores (*n* = 93 controls, 87 cases), adjusted for age. P-value is from a two-sided Wilcoxon rank-sum test. In the violin plot, boxes show interquartile ranges; whiskers, 1.5x the interquartile interval; central lines, medians; notches, confidence intervals around medians.

Each of the 1,218,284 nuclei was classified into one of seven cell types – glutamatergic neurons (43% of all nuclei), GABAergic neurons (20%), astrocytes (15%), oligodendrocytes (12%), polydendrocytes (oligodendrocyte progenitor cells, 5.5%), microglia (3.6%), and endothelial cells (1.3%) **(Fig. 1c and Extended Data Fig. 3)** – as well as neuronal subtypes defined in earlier taxonomies **(Fig. 1d-e and Extended Data Figs. 4 and 5)**. Each donor contributed nuclei of all types and subtypes **(Extended Data Figs. 3, 6, and 7)**, though subsequent analyses excluded eleven atypical samples **(Extended Data Fig. 3d)**.

### Inference of multicellular gene-programs

The data revealed substantial inter-variation in cell-type-specific gene expression levels, with highly expressed genes in each cell type exhibiting a median coefficient of variation (across donors) of about 15%.

Inter-individual variation in gene expression almost certainly arises from cell-type-specific gene-expression programs, and could in principle also be shaped by concerted changes in multiple cell types. To identify such relationships, we applied latent factor analysis, a form of machine learning which infers underlying factors from the tendency of many measurements to fluctuate together ^8^; critically, we analyzed cell-type-resolution data from all cell types at once, using inter-individual variation to enable the recognition of relationships between expression patterns in different cell types **(Fig. 1f)**. Each inferred factor was defined by a set of gene-by-cell-type loadings (revealing the distinct genes it involves in each cell type) and a set of expression levels (of the factor) in each donor **(Fig. 1f)**.

Ten latent factors together explained 30% of inter-individual variation in gene expression levels; these factors appeared to be independent of one another in their gene utilization patterns (“loadings”) and their expression levels across the individual donors **(Extended Data Fig. 8a-d)**. Inter-individual variation in the factors’ inferred expression levels arose from inter-individual variation within each 20-donor experimental set **(Extended Data Fig. 8e)**. Each factor was primarily driven by gene expression in one or a few cell types **(Fig. 1g)**.

Schizophrenia associated with just one of these latent factors (LF4) **(Fig. 1h, Extended Data Fig. 9a-e, and Supplementary Table 2)** – a factor that also associated with donor age **(Fig. 1i)**. Donors with and without schizophrenia both exhibited the decline in LF4 with age **(Fig. 1i and Extended Data Fig. 1c-d)**. Joint regression analysis confirmed independent reductions of LF4 expression by age and in schizophrenia, and detected no effect of sex **(Supplementary Table 3)**.

Factors similar to LF4 emerged in all analyses testing LF4’s robustness to analysis parameters **(Extended Data Fig. 10)**. Individuals’ LF4 expression scores also did not correlate with medication use, time of day at death, post-mortem interval, or sequencing depth **(Extended Data Fig. 9f-k)**. We also found evidence that the LF4 constellation of gene-expression changes manifests at a protein level **(Extended Data Fig. 11)**.

### Neuronal and astrocyte genes driving LF4

Of the 1,000 gene/cell-type expression traits with the strongest LF4 loadings, 99% involved gene expression in glutamatergic neurons (610), GABAergic neurons (125), or astrocytes (253) **(Fig. 1g)**. LF4 involved similar genes and expression effect directions in glutamatergic and GABAergic neurons but a distinct set of genes and effect directions in astrocytes **(Fig. 2a and Extended Data Fig. 9m)**. To identify biological processes in LF4, we applied gene set enrichment analysis (GSEA, ^9^) to the LF4 gene loadings, separately for each cell type.

**Figure 2.**
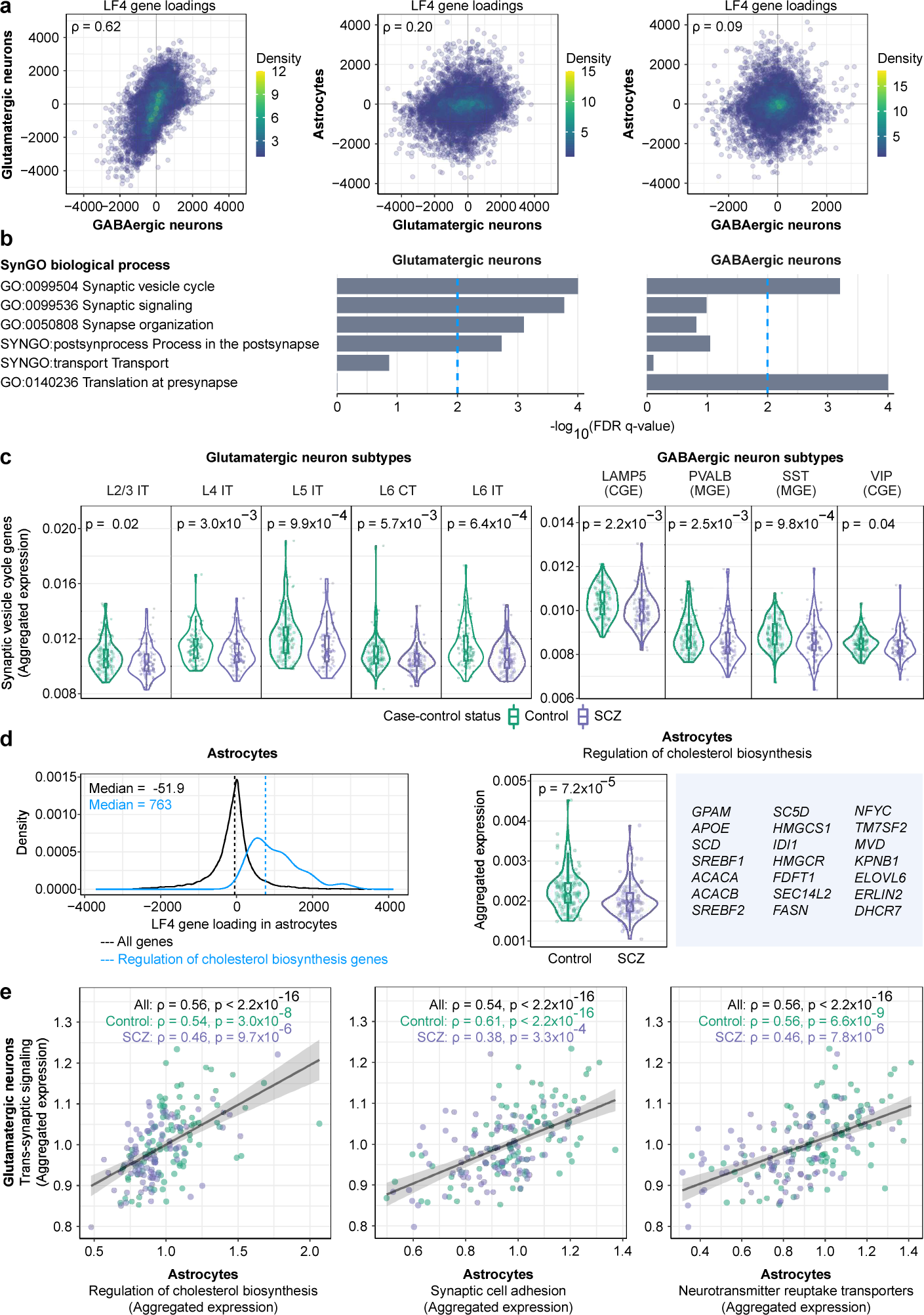
A Synaptic Neuron-Astrocyte Program (SNAP) revealed by Latent Factor 4 (LF4). **a,** Comparisons of SNAP gene recruitment between cell types. Shown, in each pairwise cell-type comparison, are the latent-factor (LF4) gene loadings of all genes expressed (≥ 1 UMI per 10^5^) in both cell types in the comparison (Spearman’s ⍴; respectively). *n* = 10,346, 11,232, 11,217 genes **b,** Concentrations of synaptic gene sets (as annotated by SynGO) in LF4’s neuronal components. **c,** Fraction of gene expression (UMIs) devoted to synaptic vesicle-cycle genes in subtypes of glutamatergic and GABAergic neurons, across 180 donors. P-values for case-control comparisons are from a two-sided Wilcoxon rank-sum test. Box plots show interquartile ranges; whiskers, 1.5x the interquartile interval; central lines, medians; notches, confidence intervals around medians. **d,** Left, distributions of astrocytes’ LF4 gene loadings for (black) all expressed genes (*n* = 18,347) and (blue) genes annotated for functions in cholesterol biosynthesis (*n* = 21) (hereafter referred to as “cholesterol biosynthesis” genes according to their GO annotation, though subsets contribute to cholesterol export and/or to synthesis of additional fatty acids). Right, proportion of astrocytic gene expression devoted to the annotated cholesterol biosynthesis genes shown, across 180 donors. P-value is from a two-sided Wilcoxon rank-sum test. Box plots show interquartile ranges; whiskers, 1.5x the interquartile interval; central lines, medians; notches, confidence intervals around medians. **e,** Concerted gene-expression variation in neurons and astrocytes. Relationships (across 180 donors) of astrocytic gene expression related to three biological activities (synapse adhesion, neurotransmitter uptake, cholesterol biosynthesis) to neuronal gene expression related to synapses (Spearman’s ⍴). Quantities plotted are the fraction of all detected nuclear mRNA transcripts (UMIs) derived from these genes in each donor’s astrocytes (x-axis) or neurons (y-axis), relative to the median expression among control donors. Shaded regions represent 95% confidence intervals for the estimated slopes.

In both glutamatergic and GABAergic neurons, LF4 involved increased expression of genes with synaptic functions **(Fig. 2b, Extended Data Fig. 9l and Supplementary Table 4)**. The strongest synaptic enrichments for both glutamatergic and GABAergic neurons involved the synaptic vesicle cycle and the presynaptic compartment; the core genes driving these enrichments encoded components of the SNARE complex and their interaction partners (*STX1A, SNAP25*, *SYP*), effectors and regulators of synaptic vesicle exocytosis (*SYT11*, *RAB3A, RPH3A*), and other synaptic vesicle components (*SV2A*, *SYN1*). In glutamatergic neurons, LF4 also appeared to involve genes encoding postsynaptic components, including signaling proteins (*PAK1*, *GSK3B, CAMK4*) and ion channels and receptors (*CACNG8, KCNN2, CHRNB2*, *GRM2, GRIA3*).

Persons with schizophrenia and persons of advanced age exhibited reduced levels of synapse-related gene expression by cortical neurons of all types **(Fig. 2c and Extended Data Fig. 12)**. In astrocytes, LF4 involved gene-expression effects distinct from those in neurons **(Fig. 2a and Extended Data Fig. 9m)**. Gene sets with roles in fatty acid and cholesterol biosynthesis and export, including genes that encode the SREBP1 and SREBP2 transcription factors and their regulators and targets, were positively correlated with LF4 and under-expressed in the cortical astrocytes of donors with schizophrenia **(Fig. 2d and Supplementary Table 4)** or advanced age **(Extended Data Fig. 13a)**. These effects appeared to be specific to astrocytes relative to other cell types **(Extended Data Fig. 14)**.

### Concerted neuron-astrocyte expression

To understand these results in terms of specific biological activities, we focused on gene sets corresponding to neuronal synaptic components and three kinds of astrocyte activities: adhesion to synapses, uptake of neurotransmitters, and cholesterol biosynthesis **(Methods: Selected gene sets)**.

The proportion of astrocyte gene expression devoted to each of these three astrocyte activities strongly correlated with the proportion of neuronal gene expression devoted to synaptic components **(Fig. 2e and Extended Data Fig. 15)**, even after adjusting for age and case-control status **(Extended Data Fig. 16)**. Donors with schizophrenia, as well as donors with advanced age, tended to have reduced expression of these genes **(Fig. 2e and Extended Data Fig. 13)**.

Because this gene expression program involves concerted effects upon the expression of (distinct) genes for synaptic components in neurons and astrocytes, we call it “SNAP” (Synaptic Neuron-Astrocyte Program), though it also involves genes with unknown functions and more-modest expression effects in additional cell types. We use donors’ LF4 expression scores to measure SNAP expression.

### Astrocyte gene-programs and SNAP

To better appreciate the astrocytic contribution to SNAP, we further analyzed the RNA-expression data from 179,764 individual astrocytes. Analysis readily recognized a known, categorical distinction among three subtypes of adult cortical astrocytes: protoplasmic astrocytes, which populate the gray matter and were the most abundant subtype; fibrous astrocytes; and interlaminar astrocytes **(Fig. 3a and Extended Data Fig. 17a-d)**. Neither schizophrenia nor age associated with variation in these subtypes’ relative abundances **(Extended Data Fig. 17e-f)**.

**Figure 3.**
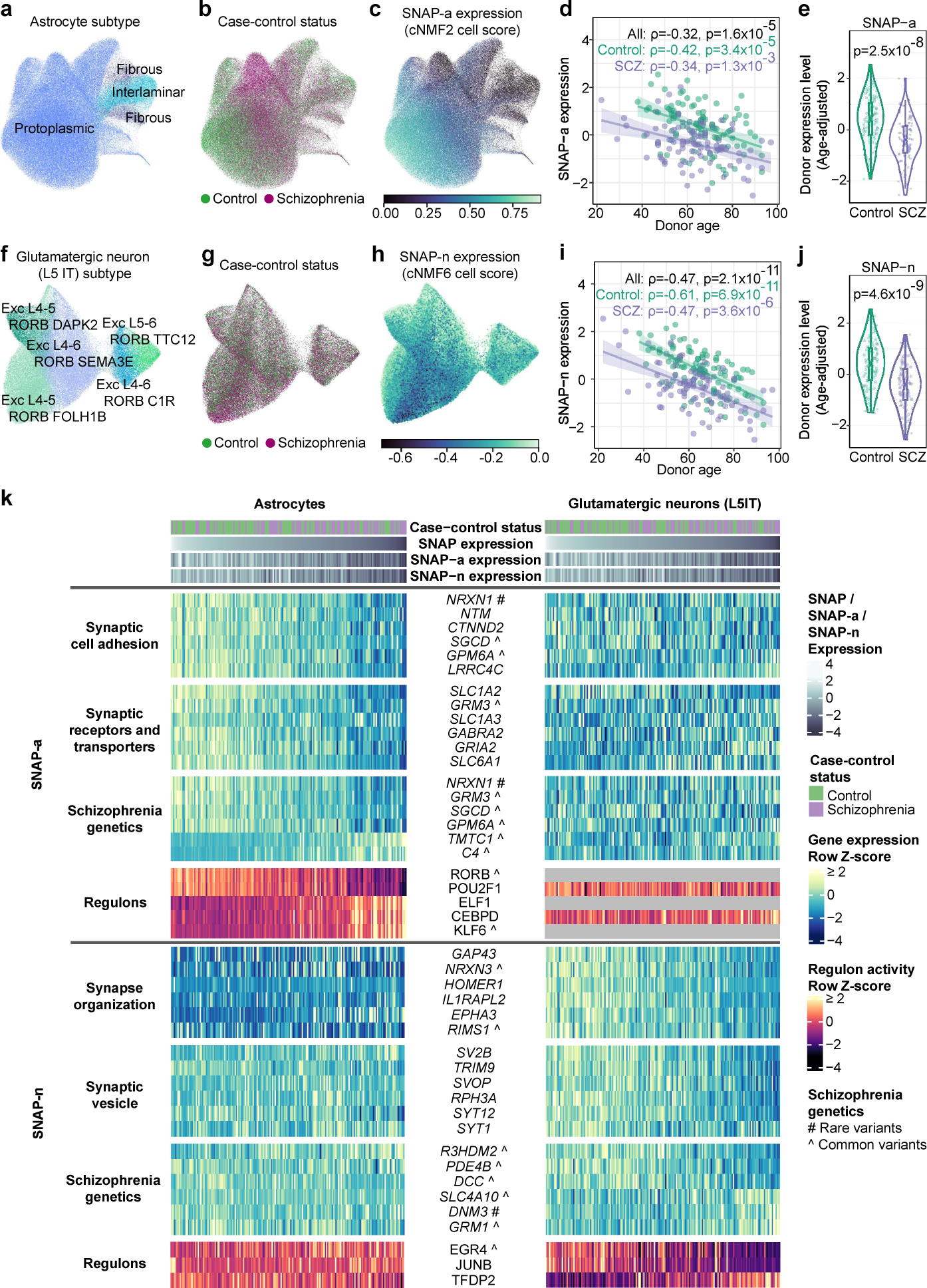
Biological states and transcriptional programs of astrocytes and L5 IT glutamatergic neurons in schizophrenia. **a-c,** UMAP of RNA-expression patterns from 179,764 astrocyte nuclei from 180 donors. Nuclei are colored by **(a)** astrocyte subtype, **(b)** schizophrenia affected/unaffected status, and **(c)** expression of the astrocyte component of SNAP (referred to as SNAP-a). **d,** Relationship of donors’ quantile-normalized SNAP-a expression scores to age (Spearman’s ⍴; *n* = 180 donors). Shaded regions represent 95% confidence intervals. **e,** Distributions of SNAP-a donor scores (age-adjusted and quantile-normalized) for persons with and without schizophrenia (*n* = 93 controls, 87 cases). P-value is from a two-sided Wilcoxon rank-sum test. Box plots show interquartile ranges; whiskers, 1.5x the interquartile interval; central lines, medians; notches, confidence intervals around medians. **f-j,** Similar plots as in **a-e** for the L5 IT glutamatergic neuron contribution to SNAP (referred to as SNAP-n; *n* = 75,929 nuclei). **k,** Heatmap showing variation in expression levels of a select set of strongly SNAP-recruited genes across the astrocytes (left) and glutamatergic neurons (right) of 180 brain donors, who are ordered from left to right by SNAP expression levels, in both the left and right panels. One set of genes (SNAP-a, above) exhibits co-regulation in astrocytes; a distinct set of genes (SNAP-n, below) exhibits co-regulation in neurons. Genes annotated by ^ are at genomic loci implicated by common genetic variation in schizophrenia ^22^. Gray bars indicate that regulon activity was not detected.

We then identified latent factors that collectively explained 25% of quantitative gene-expression variation among individual astrocytes (using cNMF, ^10^) **(Extended Data Fig. 18a-b)**. The factors appeared to capture diverse biological activities, including translation (cNMF1); zinc and cadmium ion homeostasis (cNMF7); and inflammatory responses (cNMF8) **(Supplementary Table 5)**. One factor (cNMF2) corresponded to the astrocyte component of SNAP **(Extended Data Fig. 18c-e and Supplementary Table 6)**; the strong co-expression relationships in SNAP were thus robust to the computational approach used **(Extended Data Figs. 18c-e and 19)**.

Because cNMF2 is informed by variation in the single-astrocyte expression profiles, we consider it a more precise description of the astrocyte-specific gene-expression effects in SNAP, and refer to it here as SNAP-a. Across donors, average astrocyte expression of SNAP-a associated even more strongly with schizophrenia case-control status and with age **(Fig. 3b-e and Extended Data Fig. 18f-i)**.

The strongest positive gene-set associations to SNAP-a involved adhesion to synaptic membranes and intrinsic components of synaptic membranes **(Supplementary Table 5)**. The 20 genes most strongly associated with SNAP-a **(Extended Data Fig. 20)** included eight genes with roles in adhesion of cells to synapses (*NRXN1, NTM, CTNND2, LSAMP, GPM6A, LRRC4C, LRRTM4,* and *EPHB1*) (reviewed in ^11,12^). SNAP-a also appeared to strongly recruit genes encoding synaptic neurotransmitter reuptake transporters: *SLC1A2* and *SLC1A3* (encoding glutamate transporters EAAT1 and EAAT2), and *SLC6A1* and *SLC6A11* (encoding GABA transporters GAT1 and GAT3) were all among the 1% of genes most strongly associated with SNAP-a.

We sought to relate SNAP-a to an emerging appreciation of astrocyte heterogeneity and its basis in gene expression ^13^. An earlier analysis of astrocyte molecular and morphological diversity in mice identified gene-expression modules based on their co-expression relationships^14^. SNAP-a exhibited the strongest overlap (p = 3.5 × 10^−4^, q = 0.015 by GSEA) **(Supplementary Table 5)** with the module that had correlated most closely with the size of the territory covered by astrocyte processes (the “turquoise” module in ^14^, with overlap driven by genes including *EZR* and *NTM*). A potential interpretation is that SNAP-a supports these perisynaptic astrocytic processes (PAPs ^15^).

Earlier work has identified “reactive” astrocyte states induced by strong experimental perturbations and injuries, and described as polarized cell states ^16^. We found that more than half of the human orthologs of markers for these states were expressed at levels that correlated negatively and in a continuous, graded manner with SNAP-a expression **(Extended Data Fig. 21)**. At the single-astrocyte level, SNAP-a expression exhibited continuous, quantitative variation rather than discrete state shifts **(Extended Data Fig. 18f-g)**, consistent with observations of abundant astrocyte biological variation less extreme than experimentally polarized states ^17^.

We performed an analogous cNMF analysis on the RNA-expression profiles of 75,929 glutamatergic neurons, focusing on a single, abundant subtype so that the variation among individual cells would be driven primarily by dynamic cellular programs rather than by subtype identity **(Fig. 3f)**. One factor corresponded to the neuronal gene-expression effects of SNAP; we refer to this factor as SNAP-n **(Fig. 3g-j and Supplementary Table 7)**. Like SNAP-a, average expression of SNAP-n was associated with age and with schizophrenia **(Fig. 3i-j)**. SNAP-n and SNAP-a associated with each other still more strongly, even in a controls-only, age-adjusted analysis, highlighting the close coupling of neuronal and astrocyte gene expression **(Extended Data Fig. 22)**. Although SNAP-n associated with synaptic gene-sets, the specific genes driving these enrichments were distinct from those driving SNAP-a **(Fig. 3k, Extended Data Fig. 23, and Supplementary Table 8)**.

Expression of SNAP-a and SNAP-n associated with the expression of many transcription factors and their predicted targets, and engaged distinct pathways in astrocytes and neurons **(Fig. 3k and Extended Data Figs. 22c and 24b)**: for example, SREBP1 and its well-known transcriptional targets ^18^ in astrocytes, and JUNB (AP-1) and its well-known targets ^19,20^ in neurons **(Extended Data Fig. 25)**. (The latter may reflect average neuronal activity levels in the PFC, which neuroimaging has found to decline (“hypofrontality”) in schizophrenia ^21^.) SNAP-a expression in astrocytes also associated with a RORB regulon (under-expressed in SNAP-low donors) and a KLF6 regulon (over-expressed) **(Fig. 3k and Extended Data Fig. 24b)**; common genetic variation at *RORB* and *KLF6* associates with schizophrenia ^22^.

### Schizophrenia genetics and SNAP

A key question when studying disease through human post-mortem tissue is whether observations involve disease-causing/disease-exacerbating processes, or reactions to disease circumstances such as medications. We found no relationship between SNAP expression and donors’ use of antipsychotic medications **(Extended Data Fig. 9j-k)**, or between cholesterol-biosynthesis gene expression in astrocytes and donors’ statin intake **(Extended Data Fig. 14b)**, but this does not exclude the possibility that astrocytes are primarily reacting to disease-associated synaptic hypofunction in neurons, as opposed to contributing to such hypofunction. Human genetic data provide more-powerful evidence, since inherited alleles affect risk or exacerbate disease processes rather than being caused by disease. We thus sought to evaluate the extent to which SNAP-a and SNAP-n involved genes and alleles implicated by genetic studies of schizophrenia.

Earlier work ^22–24^ has found that genes expressed most strongly by neurons (relative to other cell types), but not genes expressed most strongly by glia, are enriched for the genes implicated by genetic analyses in schizophrenia ^22–24;^ we replicated these findings in our data **(Fig. 4a and Supplementary Note)**. However, such analyses treat cell types as fixed levels of gene expression (“cell identities”), rather than as collections of dynamic transcriptional activities; SNAP-a involves a great many genes that are also strongly expressed in other cell types.

**Figure 4.**
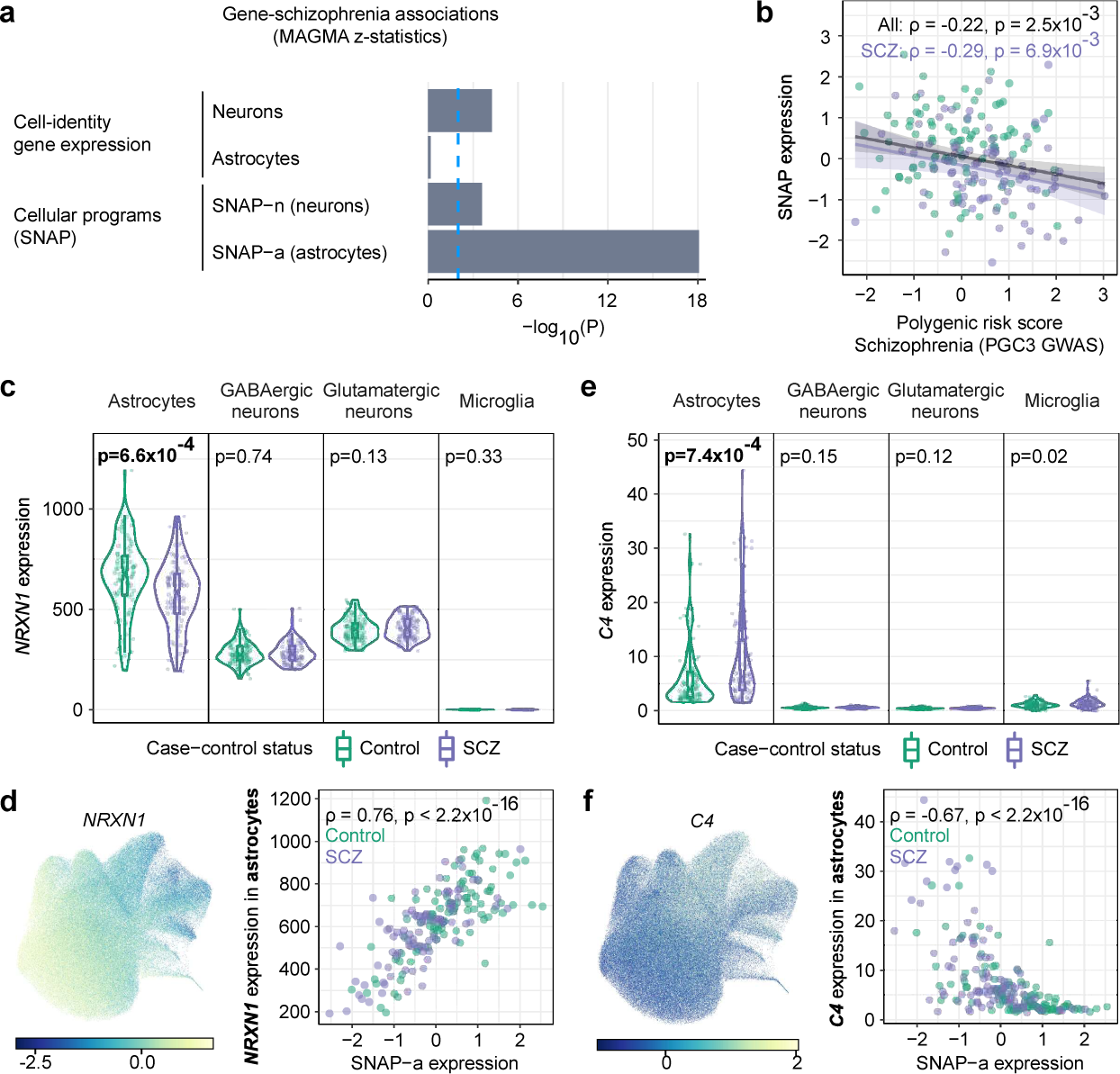
Relationship of SNAP to schizophrenia genetics. **a,** Enrichment of schizophrenia genetic association (from common variants, using MAGMA to generate a schizophrenia association Z-score for each gene) in the 2,000 genes most preferentially expressed in glutamatergic neurons and astrocytes, or the 2,000 genes whose expression is most strongly recruited by SNAP-n and SNAP-a. Values plotted are -log_10_ p-values from a joint regression analysis in which each gene set is an independent and competing predictive factor. See also **Supplementary Note**. **b,** Relationship of donors’ SNAP expression (quantile-normalized) to donors’ schizophrenia polygenic risk scores (Spearman’s ⍴; *n* = 180 donors; PGC3 GWAS from ^22^). Shaded regions represent 95% confidence intervals. **c**, *NRXN1* expression (per 10^5^ detected nuclear transcripts) in each cell type in individual donors (*n* = 93 controls, 87 cases). P-values are from a two-sided Wilcoxon rank-sum test. Box plots show interquartile ranges; whiskers, 1.5x the interquartile interval; central lines, medians; notches, confidence intervals around medians. **d,** Left, *NRXN1* expression in individual astrocytes (using the same projection as in **Fig. 3a-c)**. Values represent Pearson residuals from variance stabilizing transformation. Right, relationship of the 180 donors’ *NRXN1* expression in astrocytes to SNAP-a expression (Spearman’s ⍴). **e-f,** Similar plots as in c-d, here for complement component 4 (*C4*).

We found that the genes dynamically recruited by SNAP-a in astrocytes were 14 times more likely than other protein-coding genes to reside at genomic loci implicated by common genetic variation in schizophrenia (p = 5 × 10^−25^, 95% confidence interval (CI): 8.7-24, by logistic regression) and 7 times more likely to have strong evidence from rare variants in schizophrenia (95% CI: 2.3-21, p = 5 × 10^−4^, by logistic regression) **(Supplementary Note)**.

To evaluate whether common variation in the genes recruited by SNAP-a contributes more broadly to schizophrenia risk, beyond these strongest associations, we used gene-level association statistics from the largest schizophrenia genome-wide association study to date ^22,25^.

As expected, the strongest neuron-identity genes (as defined in the earlier work) exhibited elevated schizophrenia association, while the strongest astrocyte-identity genes did not **(Fig. 4a and Supplementary Note)**. In the same analysis, however, the genes most strongly associated with SNAP-a and SNAP-n were highly significant as additional predictive factors, particularly the genes associated with SNAP-a **(Fig. 4a)**. Analysis by linkage disequilibrium (LD) score regression ^26^ also confirmed enrichment of schizophrenia risk factors among SNAP-a genes **(Extended Data Fig. 26)**.

Polygenic risk involves thousands of common alleles across the genome, whose effects converge upon unknown biological processes. A polygenic risk score (PRS) for schizophrenia associated with reduced expression of SNAP but not with the other latent factors **(Fig. 4b and Extended Data Fig. 27)**. Higher polygenic risk also associated with deeper decline in SNAP among persons with schizophrenia **(Fig. 4b)**.

To better understand such relationships, we explored the relationship of SNAP-a to genetic risk through two specific genes: Neurexin-1 (*NRXN1*) and complement component 4 (*C4*). Exonic deletions within *NRXN1* greatly increase risk for schizophrenia ^27,28^. Our data indicated that astrocytic, but not neuronal, *NRXN1* expression was reduced in persons with schizophrenia and among persons over 70 years of age **(Fig. 4c and Extended Data Fig. 28a-b)**. Inter-individual variation in astrocytic *NRXN1* expression strongly associated with SNAP-a **(Fig. 4d)**.

Increased copy number of the complement component 4 (*C4A*) gene more-modestly increases risk for schizophrenia ^29^; far more inter-individual variation in *C4* gene expression (>80%) arises from unknown, dynamic effects on *C4* expression ^29,30^. We found that astrocytes, rather than neurons or microglia, are the main site of *C4* (including *C4A* and *C4B*) RNA expression in human prefrontal cortex **(Fig. 4e and Extended Data Fig. 28c)**. Donors with lower-than-average expression of SNAP-a tended to have greatly increased *C4* expression: such donors included 43 of the 44 donors with highest *C4* expression levels, and their astrocytes expressed 3.2-fold more *C4* than astrocytes in donors with above-average expression of SNAP-a did **(Fig. 4f)**. *C4* expression was also greatly increased among donors over 70 years of age **(Extended Data Fig. 28d-e)**.

## DISCUSSION

Here we discovered SNAP (Synaptic Neuron-Astrocyte Program), concerted gene-expression programs implemented by cortical neurons and astrocytes to corresponding degrees in the same individuals. SNAP expression varied even among neurotypical control brain donors and may be a core axis of human neurobiological variation, with potential implications for cognition and plasticity that will be important to understand.

SNAP appears to involve many genes that contribute to synapses and to astrocyte-synapse interactions **(Figs. 2 and 3k, Supplementary Table 9, and Extended Data Figs. 20 and 23)** ^31,32^. The genes associated with SNAP-a suggested a potential role in supporting perisynaptic astrocyte processes, motile astrocyte projections whose morphological plasticity and interactions with synapses can promote synaptic stability ^15^. Diverse lines of work increasingly reveal a key role for astrocytes in regulating the ability of synaptic networks to acquire and learn new information, for example by lowering thresholds for activity and synaptic plasticity ^33,34^.

An intriguing aspect of SNAP involved the astrocytic regulation of genes with roles in fatty acid and cholesterol biosynthesis and cholesterol export, which strongly correlated (across donors) with expression of synaptic-component genes by neurons **(Fig. 2d-e)**. Earlier research has defined a potential rationale for this neuron-astrocyte coordination: synapses and dendritic spines – synapse-containing morphological structures – require large amounts of cholesterol that astrocytes supply ^35^. Declines in cholesterol biosynthesis have previously been noted in mouse models of brain disorders ^36,37^ that (like schizophrenia and aging) involve cognitive losses, cortical thinning, and reduction in neuropil.

Schizophrenia and aging both brought substantial reductions in SNAP expression **(Fig. 1i-j)**. Neuropsychologic, neuroimaging, and neuronal microstructural studies have long noted similar changes in schizophrenia and aging ^1,2,38–47^. Inherited genetic risk for schizophrenia associates with decreased measures of cognition in older individuals ^48,49^, and schizophrenia greatly increases risk of dementia later in life ^50^. Our results suggest that these relationships between schizophrenia and aging arise from shared cellular and molecular changes.

Under-expression of SNAP could in principle underlie longstanding microstructural observations ^41–47^ of reduced numbers of dendritic spines – synapse-containing morphological structures – on cortical neurons in aged humans and primates and in persons with schizophrenia. These microstructural observations appear to arise from highly plastic thin spines and thus may reflect reduced rates of continuous synapse formation and stabilization (rather than pruning of mature synapses) ^42–47^. The gene-expression changes we observed in human dlPFC **(Fig. 2c)** suggest that cortical neurons of all types, including glutamatergic and GABAergic neurons, may be affected by such changes.

It is intriguing to consider whether pharmacotherapies or other interventions could be developed to promote SNAP as a way to address cognitive symptom domains in schizophrenia and aging such as cognitive flexibility, working memory, and executive function deficits, continuous and disabling features which are typically not improved by available treatments ^1^.

An important future direction will be to determine the extent to which SNAP is present in other brain areas, and the relationship of SNAP to molecular and physiological changes in dendrites, synapses, and perisynaptic astrocyte processes. Additional questions involve the molecular mechanisms that accomplish neuron-astrocyte coordination and the extent to which SNAP supports learning and/or cognitive flexibility.

SNAP was made visible by human inter-individual biological variation. Though controlled laboratory experiments usually try to eliminate genetic and environmental variation, natural variation may be able to reveal cell-cell coordination and regulatory programs in many tissues and biological contexts, offering new ways to identify pathophysiological processes within and beyond the human brain.

## METHODS

### Ethical compliance

Brain donors were recruited by the Harvard Brain Tissue Resource Center/NIH NeuroBioBank (HBTRC/NBB), in a community-based manner, across the USA. Human brain tissue was obtained from the HBTRC/NBB. The HBTRC procedures for informed consent by the donor’s legal next-of-kin and distribution of de-identified post-mortem tissue samples and demographic and clinical data for research purposes are approved by the Mass General Brigham Institutional Review Board. Post-mortem tissue collection followed the provisions of the United States Uniform Anatomical Gift Act of 2006 described in the California Health and Safety Code section 7150 and other applicable state and federal laws and regulations. Federal regulation 45 CFR 46 and associated guidance indicates that the generation of data from de-identified post-mortem specimens does not constitute human participant research that requires institutional review board review.

### Donors for single nucleus RNA-seq

Donor information with anonymized donor IDs is available in **Supplementary Table 1**. Consensus diagnosis of schizophrenia was carried out by retrospective review of medical records and extensive questionnaires concerning social and medical history provided by family members. Several regions from each brain were examined by a neuropathologist. We excluded subjects with evidence for gross and/or macroscopic brain changes, or with clinical history consistent with cerebrovascular accident or other neurological disorders. Subjects with Braak stages III or higher (modified Bielchowsky stain) were excluded. None of the subjects had significant reported history of substance dependence within 10 or more years from death, as further corroborated by negative toxicology reports. Absence of recent substance abuse is typical for samples from the HBTRC, which receives exclusively community-based tissue donations.

Exposure to psychotropic and neurotropic medications was assessed on the basis of medical records. Estimated daily milligram doses of antipsychotic drugs were converted to the approximate equivalent of chlorpromazine as a standard comparator ^51^. These values are reported as lifetime, as well as last six months’ of life, grams per patient. Exposure to other classes of psychotropic drugs was reported as present or absent.

### Single-nucleus library preparation and sequencing

We analyzed the dlPFC (Brodmann area 46 (BA46)), which exhibits functional and microstructural abnormalities in schizophrenia ^52,53^ and in aging ^46^. Frozen tissue blocks containing BA46 were obtained from the HBTRC. We used single-nucleus rather than single-cell RNA-seq to avoid effects of cell morphology upon ascertainment, and because nuclear (but not plasma) membranes (but not plasma membranes) remain intact in frozen post-mortem tissue. Nuclear suspensions from frozen tissue were generated following the protocol we have made available at dx.doi.org/10.17504/protocols.io.4r3l22e3xl1y/v1. To ensure that batch compositions were balanced, researchers were not blinded to the batch allocation or processing order of each specimen. To maximize the technical uniformity of the snRNA-seq data, we processed sets of 20 brain specimens (each consisting of affected and control donors) at once as a single pooled sample. Specimens were allocated into batches of 20 specimens per batch, ensuring that the same number of cases and age-matched controls (10 per group), and men and women (10 per group) were included in each batch. Some donors were re-sampled across multiple batches to enable quality control analyses **(Extended Data Fig. 2)**. Specimens from cases and age-matched controls were also processed in alternating order within each batch. Researchers had access to unique numerical codes assigned to the donor-of-origin of each specimen as well as basic donor metadata (e.g. case-control status, age, sex).

Some 50 mg of tissue was dissected from the dlPFC of each donor – sampling across the cortical layers and avoiding visible concentrations of white matter – and used to extract nuclei for analysis. GEM generation and library preparation followed the 10X Chromium Single Nuclei 3’ v3.1 protocol (version #CG000204_ChromiumNextGEMSingleCell3’v3.1_Rev D). We encapsulated nuclei into droplets using approximately 16,500 nuclei per reaction, understanding that about 95% of all doublets (cases in which two nuclei were encapsulated in the same droplet) would consist of nuclei from distinct donors and thus be recognized by the Dropulation analysis ^7^ as containing combinations of SNP alleles from distinct donors. cDNA amplification was performed using 13 PCR cycles.

Raw sequencing reads were aligned to the hg38 reference genome with the standard Drop-seq (v2.4.1) ^54^ workflow, modified so that reads from *C4* transcripts would not be discarded as multi-mapping (see Methods below, ***C4*: MetaGene discovery**). Reads were assigned to annotated genes if they mapped to exons or introns of those genes. Ambient / background RNA were removed from digital gene expression (DGE) matrices with CellBender (v0.1.0) ^55^ remove-background.

### Genotyping and donor assignment from snRNA-seq data

We used combinations of hundreds of transcribed SNPs to assign each nucleus to its donor-of-origin, using the Dropulation software (v2.4.1) ^7^. Previous Dropulation analyses of stem cell experiments have used whole-genome sequence (WGS) data on the individual donors for such analyses ^7^. For the current work, we developed a cost-efficient approach based on SNP array data with imputation. Genomic DNA from the individual brain donors was genotyped by SNP array (Illumina GSA).

Raw Illumina IDAT files from the GSAMD-24v1-0_20011747 array (2,085 samples) and GSAMD-24v3-0-EA_20034606 array (456 samples) were genotyped using GenCall (v3.0.0) ^56^ and genotypes were phased using SHAPEIT4 (v4.2.2) ^57^ by processing the data through the MoChA workflow (v2022-12-21) ^58,59^ (https://github.com/freeseek/mochawdl) using default settings and aligning markers against GRCh38. *APOE* genotypes for marker rs429358 were removed due to unreliable genotypes. To improve phasing, genotypes from the McLean cohort were combined with genotypes from the Genomic Psychiatry Cohort with IDAT files available also from the GSAMD-24v1-0_20011747 array (5,689 samples) ^60^. After removing 128 samples recognized as duplicates, phased genotypes were then imputed using IMPUTE5 (v1.1.5) ^61^ by processing the output data from the MoChA workflow using the MoChA imputation workflow and using the high coverage 1000 Genomes reference panel for GRCh38 ^62^ including 73,452,470 non-singleton variants across all the autosomes and chromosome X. Only SNPs with imputation quality INFO > 0.95 were used for donor assignments. Using this approach, we found that 99.6% of nuclei could be assigned confidently to a donor **(Extended Data Fig. 2b)**.

To evaluate the accuracy of this method of donor assignment, we genotyped a pilot cohort of 11 donors by both whole-genome sequencing (WGS) and by SNP array. Importantly, the two methods had 100% concordance on the assignment of individual nuclei to donors, validating both our computational donor-assignment method and the sufficiency of the SNPs-plus-imputation approach **(Extended Data Fig. 2a)**. SNP data for the individual donors are available in NeMO (accession number nemo:dat-bmx7s1t).

Following donor assignment, DGE matrices from all libraries in each batch (7 to 8 libraries per batch) were merged for downstream analyses.

### Cell-type assignments

All classification models for cell assignments were trained using scPred (v1.9.2) ^63^. DGE matrices were processed using the following R and python packages: Seurat (v3.2.2) ^64^, SeuratDisk (v0.0.0.9010) ^65^, anndata (v0.8.0) ^66^, numpy (v1.17.5) ^67^, pandas (v1.0.5) ^68,69^, and Scanpy (v1.9.1) ^70^.

#### Cell types

##### Model training

The classification model used for cell-type assignments was trained on the DGE matrix from batch 6 (BA46_2019-10-16), which was annotated as follows. Nuclei with fewer than 400 detected genes and 100 detected transcripts were removed from the DGE matrix from this batch. After normalization and variable gene selection, the DGE matrix was processed through an initial clustering analysis using independent component analysis (ICA, using fastICA (v1.2-1)^71^) as previously described ^72^. This analysis produced clustering solutions with 43 clusters of seven major cell types (astrocytes, endothelial cells, GABAergic neurons, glutamatergic neurons, microglia, oligodendrocytes, polydendrocytes) that could be identified based on expression of canonical marker genes (markers in **Extended Data Fig. 3**). (We note that ∼9% of cells within clusters annotated as endothelial cells do not express canonical endothelial cell markers, but rather those of pericytes; these ∼1,400 cells have been grouped together with endothelial cells for downstream analyses.) scPred was trained on this annotated DGE matrix, and the resulting model was subsequently used to make cell-type assignments for the remaining batches’ DGE matrices.

##### Filtering

Following an initial cell-type classification using the above model, the DGE matrices were filtered further to remove any remaining heterotypic doublets missed by scPred. First, raw DGE matrices from each of the 11 batches were subsetted to form separate DGE matrices for each of the 7 major cell types (77 subsetted DGE matrices total). Each subsetted DGE matrix was normalized using sctransform (v0.3.1) ^64^ with 7,000 variable features, scaling, and centering. For each cell type, normalized DGE matrices from the 11 batches were merged and clustered together in Scanpy (v1.9.1) ^70^ using 50 PCs, batch correction by donor using BBKNN (v1.5.1) ^73^, and Leiden clustering using a range of resolutions. The most stable clustering resolution for each cell type was selected using clustree (v0.4.4) ^74^. Clusters expressing markers of more than one cell type were determined to be heterotypic doublets; cell barcodes in these clusters were discarded from the above DGE matrices, and these filtered DGE matrices were then carried forward for integrated analyses across batches.

#### Neuronal subtypes

Classification models for neuronal subtypes were trained using DGE matrices from ^75^ that were subsetted to glutamatergic or GABAergic neuron nuclei in middle temporal gyrus (MTG). While a similar dataset exists for human brain nuclei from primary motor cortex (M1) ^76^, we only trained the model on the MTG dataset as M1 lacks a traditional layer 4 (L4), while BA46 does have a L4.

The neuronal subtypes in this dataset include glutamatergic neuron subtypes of distinct cortical layers and with predicted intratelencephalic (IT), extratelencephalic (ET), corticothalamic (CT), and near-projecting (NP) projection patterns, as well as the four cardinal GABAergic neuron subtypes arising from the caudal (CGE: *LAMP5*+, *VIP*+) and medial (MGE: *PVALB*+, *SST*+) ganglionic eminences.

We made the following adjustments to the MTG annotations prior to model training. First, as subtype-level annotations (e.g. L5 IT, as used in ^76^ for M1) were not available for the MTG dataset, we inferred these based on M1/MTG cluster correspondences (from Extended Data Fig. 10 in ^76^). Second, we reassigned the following glutamatergic neuron types in MTG from the L4 IT subtype (as inferred by integration with M1 in ^76^) to the L2/3 IT subtype: Exc L3−5 RORB FILIP1L, Exc L3−5 RORB TWIST2, and Exc L3−5 RORB COL22A1. This was on the basis of their properties described in other studies – for example, the Exc L3−5 RORB COL22A1 type has been described as a deep L3 type by Patch-seq ^77^ – and by the expression of their marker genes on a two-dimensional projection of the RNA-expression profiles of glutamatergic neuron nuclei **(Extended Data Fig. 4)**.

Feature plots for neuronal subtypes **(Extended Data Fig. 4 and Extended Data Fig. 5)** were generated using markers from the repository in https://bioportal.bioontology.org/ontologies/PCL (v1.0, 2020-04-26) ^75,76,78^, specifically those for neuronal subtypes from MTG.

#### Astrocyte subtypes

Normalized, filtered DGE matrices from the 11 batches were merged and clustered together in scanpy using 8 PCs, batch correction by donor using bbknn ^73^, and Leiden clustering using a range of resolutions. The most stable resolution that created distinct clusters for putative astrocyte subtypes (resolution 1.3) was selected using clustree ^74^. Feature plots for astrocyte subtypes previously described in both MTG and M1 ^75,76^ **(Extended Data Fig. 17)** were generated using markers from the repository in https://bioportal.bioontology.org/ontologies/PCL (v1.0, 2020-04-26) ^75,76,78^. Leiden clusters were assigned to one of three astrocyte subtypes on the basis of expression of these subtype markers.

### Donor exclusion

Donors were excluded on the basis of unusual gene-expression profiles and/or cell-type proportions (potentially related to agonal events) as outlined below.

#### Expression

Donors with fewer than 1,000 total UMIs in any cell type were first excluded. Next, for each cell type, gene-by-donor expression matrices comprising the remaining donors were scaled to 100,000 UMIs per donor and filtered to the top expressing genes (defined as having at least 10 UMIs per 100,000 for at least one donor; these were among the top 12-19% of expressed genes). These filtered expression matrices by cell type were merged into a single expression matrix that was used to calculate each donor’s pairwise similarity to the other donors (Pearson correlations of log_10_-scaled expression values across genes). The median of these pairwise correlation values was determined to be the conformity score for each donor. To identify outliers, these donor conformity scores were converted to modified Z-scores (M_i_) for each donor as described in ^79^:

M_i_ = 0.6745 * (x_i_ - x̃) / MAD

where

x_i_ : The donor’s conformity score

x̃ : The median of donor conformity scores

MAD: The median absolute deviation of donor conformity scores

Donors whose modified Z-scores had absolute values > 5 were excluded. This approach flagged a total of 5 donors (1 who had low UMI counts and 4 who were outliers on the basis of expression).

#### Cell-type proportions

Each donor’s pairwise similarity to the other donors was determined on the basis of cell-type proportions (i.e., the values plotted in **Extended Data Fig. 3c-d)**. Donor conformity scores and modified Z-scores based on these values were calculated for each donor using the same approach described above for expression values. Donors whose modified Z-scores had absolute values > 15 were excluded. This approach flagged a total of 9 donors, 2 of whom were also flagged as expression outliers.

Between the two approaches, a total of 11 unique donors were flagged as outliers (4 control, 7 schizophrenia) and excluded from downstream analyses.

### Latent factor analysis

#### snRNA-seq data

Our approach was to (i) create a gene-by-donor matrix of expression measurements for each of seven cell types; (ii) concatenate these matrices into a larger matrix in which each gene is represented multiple times (once per cell type); and (iii) perform latent factor analysis ^8,80^ on this larger matrix. We selected probabilistic estimation of expression residuals (PEER) ^81^ over other approaches (e.g. PCA) for inferring latent variables as it is more sensitive and less dependent on the number of factors modeled. A major pitfall to avoid when performing latent factor analysis is obtaining highly correlated factors due to overfitting. The latent factors we have inferred are independent from each other when we compare their gene loadings **(Extended Data Fig. 8c)**, enabling us to proceed with downstream analyses based on these factors.

Raw, filtered DGE matrices from each of the 11 batches were subsetted to form separate DGE matrices for each of the 7 major cell types (77 subsetted DGE matrices total). For each subsetted DGE matrix, cell barcodes from outlier donors were excluded, the DGE matrix was normalized using sctransform (v0.3.1) ^64^ with 3,000 variable features, and the output of Pearson residual expression values (with all input genes returned) was exported to a new DGE matrix. For each cell type, these new expression values in the 11 normalized DGE matrices were summarized across donors (taking the sum of residual expression values) to create a gene-by-donor expression matrix. Each of these expression matrices was filtered to the top 50% of expressed genes (based on feature counts scaled to 100,000 transcripts per donor), yielding expression matrices with approximately 16,000 to 18,000 genes per cell type. Within each expression matrix, each gene name was modified with a suffix to indicate the cell type of origin (e.g. *ACAP3* to *ACAP3_astrocyte*), and the 7 expression matrices were combined to produce a single expression matrix with expression values from all 7 cell types for each donor (see **Fig. 1f** for schematic). This expression matrix was used as the input to latent factor analysis with PEER (v1.0) ^81^ using default parameters and a range of requested factors *k*.

Though we looked for correlations of these factors with technical variables, these analyses were negative, with one exception: Latent Factor 2 (LF2) appeared to capture quantitative variation in the relative representation of deep and superficial cortical layers in each dissection **(Extended Data Fig. 8f)**.

Latent factor donor expression values were adjusted for age by taking the residuals from a regression of the donor expression values against age.

To improve the visualization of latent factor donor expression values while leaving the results of statistical analyses unchanged, quantile-normalized values were calculated using the formula qnorm(rank(x) / (length(x)+1)). Figure legends indicate when these quantile-normalized values are used.

#### Proteomics data

Protein intensities from the *LRRK2* Cohort Consortium (LCC) cohort in ^82^ were downloaded from the ProteomeXchange Consortium (dataset identifier PXD026491) and subset to those peptides that passed the Q-value threshold in at least 25% of all analyzed samples. These were further subset to intensities from control donors without the LRRK2 G2019S mutation and without erythrocyte contamination (*n* = 22 donors). After normalization of the protein intensities with sctransform (v0.3.1) ^64^, the output of Pearson residual expression values (with all input proteins returned) was exported to a new matrix. This matrix of normalized protein intensities was used as the input to latent factor analysis with PEER (v1.0) ^81^ using default parameters.

For comparisons of CSF protein loadings to SNAP gene loadings in **Extended Data Fig. 11**, each gene in SNAP was represented by a single composite loading representing gene loadings from all cell types. This composite loading was determined for each gene by first calculating the median expression of each gene (in each cell type), then calculating a new loading onto SNAP weighted across cell types by these median expression values.

### Rhythmicity analysis

For **Extended Data Fig. 9f**, rhythmicity analyses were performed as in ^83^ using scripts from (https://github.com/KellyCahill/Circadian-Analysis-) and donors’ time of death in zeitgeber time (ZT). Analyses also used the following packages: lme4 (v1.1-31) ^84^, minpack.lm (v1.2-4) ^85^.

### Gene set enrichment analysis

For gene set enrichment analysis (GSEA) ^9,86^ on latent factors inferred by PEER, the C5 Gene Ontology collection (v7.2) ^87,88^ from the Molecular Signatures Database ^89,90^ was merged with SynGO (release 20210225) ^91^’s biological process (BP) and cell component (CC) gene lists. Gene sets from this merged database that were enriched in each latent factor were identified with GSEAPreranked in GSEA (v4.0.3) ^9,86^ using 10,000 permutations and gene loadings as the ranking metric.

For astrocyte latent factors inferred by cNMF ^10^, GSEA was performed as described above with the addition of the following custom gene sets to the database:

- PGC3_SCZ_GWAS_GENES_1TO2_AND_SCHEMA1_GENES: A gene set comprising genes implicated in human-genetic studies of schizophrenia, including genes at 1-2 gene loci from GWAS (PGC3, ^22^ and genes with rare coding variants (FDR < 0.05 from ^23^).
- Gene sets for each of the seven astrocyte subclusters identified in ^14^.
- Gene sets for each of the 62 “color” module eigengenes identified by WGCNA in ^14^.
- Gene sets for each of the six astrocyte subcompartments analyzed in ^92^, comprising genes encoding the proteins that were unique to or enriched in these subcompartments.

For L5 IT glutamatergic neuron latent factors inferred by cNMF, GSEA was performed as described above with the addition of the following custom gene sets to the database:

- PGC3_SCZ_GWAS_GENES_1TO2_AND_SCHEMA1_GENES: A gene set comprising genes implicated in human-genetic studies of schizophrenia, including genes at 1-2 gene loci from GWAS (PGC3, ^22^ and genes with rare coding variants (FDR < 0.05 from ^23^).

### Selected gene sets

Based on the results of the gene set enrichment analyses (GSEA) described above, we selected several of the top-enriched gene sets for further analyses. These are referred to in the figures with labels modified for brevity, but are described in further detail below. Lists of genes in each gene set are in **Supplementary Table 9**.

- “Integral component of postsynaptic density membrane” **(Extended Data Fig. 13, Extended Data Fig. 15, and Extended Data Fig. 16)**: core genes contributing to the enrichment of GO:0099061 (v7.2, integral component of postsynaptic density membrane) in the glutamatergic neuron component of LF4 (SNAP).
- “Neurotransmitter reuptake transporters” **(Fig. 2e, Extended Data Fig. 13, Extended Data Fig. 15, and Extended Data Fig. 16)**: genes from among the 100 genes most strongly recruited by cNMF2 (SNAP-a) with known functions as neurotransmitter reuptake transporters. These include core genes contributing to the enrichment of GO:0140161 (v7.2, monocarboxylate: sodium symporter activity) in SNAP-a.
- “Presynapse” **(Extended Data Fig. 13, Extended Data Fig. 15, and Extended Data Fig. 16)**: core genes contributing to the enrichment of GO:0098793 (v7.2, presynapse) in the GABAergic neuron component of LF4 (SNAP).
- “Regulation of cholesterol biosynthesis” **(Fig. 2d-e, Extended Data Fig. 13, Extended Data Fig. 14, Extended Data Fig. 15, Extended Data Fig. 16, and Extended Data Fig. 24d)**: core genes contributing to the enrichment of GO:0045540 (v7.2, regulation of cholesterol biosynthetic process) in the astrocyte component of LF4 (SNAP). This enrichment is of interest as cholesterol is an astrocyte-supplied component of synaptic membranes ^35,93,94^. Products of this biosynthetic pathway also include other lipids and cholesterol metabolites with roles at synapses, including 24S-hydroxycholesterol, a positive allosteric modulator of NMDA receptors ^95^. Although we refer to this gene set by this label based on its annotation by GO, we note that subsets of these genes contribute to cholesterol export and/or to synthesis of additional fatty acids.
- “Schizophrenia genetics” **(Fig. 3k and Extended Data Fig. 24a)**: prioritized genes from ^23^ (FDR < 0.05) or ^22^.
- “Synapse organization” **(Fig. 3k)**: core genes contributing to the enrichment of GO:0050808 (v7.2, synapse organization) in cNMF6 (SNAP-n).
- “Synaptic cell adhesion” **(Fig. 2e, Fig. 3k, Extended Data Fig. 13, Extended Data Fig. 15, Extended Data Fig. 16, and Extended Data Fig. 24a)**: genes from among the 20 genes most strongly recruited by cNMF2 (SNAP-a) with known functions in synaptic cell-adhesion. This biological process was selected due to the enrichment of GO:0099560 (v7.2, synaptic membrane adhesion) in SNAP-a.
- “Synaptic receptors and transporters” **(Fig. 3k, Extended Data Fig. 24a, and Extended Data Fig. 24c)**: genes from among the 100 genes most strongly recruited by cNMF2 (SNAP-a) with known functions as synaptic receptors and transporters.
- “Synaptic vesicle” **(Fig. 3k)**: core genes contributing to the enrichment of GO:0008024 (v7.2, synaptic vesicle) in cNMF6 (SNAP-n).
- “Synaptic vesicle cycle” **(Fig. 2c and Extended Data Fig. 12)**: core genes contributing to the enrichment of GO:0099504 (v7.2, synaptic vesicle cycle) in the glutamatergic and GABAergic neuron components of LF4 (SNAP).
- “Trans-synaptic signaling” **(Fig. 2e, Extended Data Fig. 13, and Extended Data Fig. 16)**: core genes contributing to the enrichment of GO:0099537 (v7.2, trans-synaptic signaling) in the glutamatergic neuron component of LF4 (SNAP).

Gene sets displayed in **Fig. 2b** are the SynGO terms most strongly enriched in each top-level category (among biological processes: process in the presynapse, synaptic signaling, synapse organization, process in the postsynapse, transport, and metabolism respectively).

### Analysis of astrocyte and glutamatergic L5 IT neuron gene-expression programs

#### Consensus non-negative matrix factorization

Consensus non-negative matrix factorization (cNMF) (v1.2) ^10^ was performed on both astrocyte and glutamatergic L5 IT neurons. We used cNMF because of its scalability to the astrocyte and glutamatergic L5 IT neuron data sets. The cNMF protocol detailed in their github tutorial for PBMC cells (https://github.com/dylkot/cNMF/blob/master/Tutorials/analyze_pbmc_example_data.ipynb) was followed for the initial data filtering and analysis. For both data sets, data was filtered to remove cells with fewer than 200 genes or 200 UMIs. Genes expressed in fewer than 10 cells were removed. Factorization was run on raw counts data after filtering, with iterations of factorization run for each *k* (factors requested), with a *k* ranging from 3 to 30.

The astrocyte raw counts data contained 179,764 cells and 42,651 genes, of which 0 cells and 9,040 genes were excluded. Based on PCA of the gene expression matrix and the cNMF stability report, factorization with *k*=11 was selected for further analysis. The 11 cNMF factors together explained 25% of variation in gene expression levels among single astrocytes.

The L5 IT raw counts data contained 75,929 cells and 42,651 genes, of which 0 cells and 8,178 genes were excluded. Based on the PCA of the gene expression matrix and the cNMF stability report, factorization with *k*=13 was selected for further analysis. The 13 cNMF factors together explained 44% of variation in gene expression levels among single L5 IT glutamatergic neurons. To align the direction of interpretation across all 3 analyses (SNAP, SNAP-a, and SNAP-n), we took the negative of cNMF Factor 6 (SNAP-n) cell scores, gene loadings, and donor scores.

The latent factor usage matrix (cell by factor) was normalized prior to analysis to scale each cell’s total usage across all factors to 1.

#### Co-varying neighborhood analysis

To further assess the robustness of the astrocyte gene-expression changes represented by SNAP and SNAP-a, we employed a third computational approach, co-varying neighborhood analysis (CNA) (v0.1.4) ^96^. The protocol detailed in their github tutorial (https://nbviewer.org/github/yakirr/cna/blob/master/demo/demo.ipynb) was followed for data preprocessing and analysis.

Pilot association tests to find transcriptional neighborhoods associated with schizophrenia case-control status were first performed using the default value for Nnull. These pilot analyses evaluated the effects of batch correction (by batch or donor) and covariate correction (by age, sex, PMI, number of UMIs, or number of expressed genes). Nearly all analyses yielded highly similar neighborhoods associated with case-control status with the same global p-value (p=1×10^−4^), with the exception of batch correction by donor which yielded p=1. The final association test described in **Extended Data Fig. 19** was performed with an increased value for Nnull (Nnull=1000000) and without additional batch or covariate correction.

### Regulatory network inference

The goal of pySCENIC ^97,98^ is to infer transcription factors and regulatory networks from single cell gene expression data. The pySCENIC (v0.11.2) protocol detailed in the github tutorial for PBMC cells (https://github.com/aertslab/SCENICprotocol/blob/master/notebooks/PBMC10k_SCENIC-protocol-CLI.ipynb) was followed for the initial data filtering and analysis. For both astrocytes and L5 IT glutamatergic neurons, data was filtered to remove cells with fewer than 200 genes, and genes with fewer than 3 cells. Cells with high MT expression (>15% of their total transcripts) were removed.

The gene regulatory network discovery adjacency matrix was inferred by running Arboreto on the gene counts matrix and a list of all transcription factors provided by the authors (https://resources.aertslab.org/cistarget/tf_lists/allTFs_hg38.txt)] to generate an initial set of regulons. This set was further refined using ctx, which removes targets that are not enriched for a motif in the transcription factor using a provided set of human specific motifs (https://resources.aertslab.org/cistarget/motif2tf/motifs-v9-nr.hgnc-m0.001-o0.0.tbl) and cis targets (https://resources.aertslab.org/cistarget/databases/homo_sapiens/hg38/refseq_r80/mc9nr/gene_based). Finally, aucell was run to generate the per-cell enrichment scores for each discovered transcription factor.

### Super-enhancer analysis

Preparation of input BAM files: FASTQ files of bulk H3K27ac HiChIP data from middle frontal gyrus ^99^ were downloaded from GEO (accessions GSM4441830 and GSM4441833). Demultiplexed FASTQ files were trimmed with Trimmomatic (v0.33) ^100^ using the parameter SLIDINGWINDOW:5:30. Trimmed reads were aligned to the hg38 reference genome with Bowtie2 (v2.2.4) ^101^ using default parameters. Uniquely mapped reads were extracted with samtools (v1.3.1) ^102^ view using the parameters -h -b -F 3844 -q 10.

Preparation of input constituent enhancers: FitHiChIP interaction files for H3K27ac from middle frontal gyrus ^99^ were downloaded from GEO (accessions GSM4441830 and GSM4441833). These were filtered to interacting bins (at interactions with q-value < 0.01) that overlap bulk H3K27ac peaks in the one-dimensional HiChIP data in both replicates. Next, these bins were intersected with IDR-filtered scATAC-seq peaks in isocortical and unclassified astrocytes (peaks from clusters 13, 15, 17, downloaded from GEO accession GSE147672 ^99^). Unique coordinates of these filtered regions were converted to GFF files.

Super-enhancers (SEs) were called with ROSE (v1.3.1) ^103,104^ using the input files prepared above and the parameters -s 12500 -t 2500. Coordinates of promoter elements for *H. sapiens* (Dec 2013 GRCh38/hg38) were downloaded from the Eukaryotic Promoter Database (EPD) ^105^ using the “EPDnew selection tool” (https://epd.expasy.org/epd/EPDnew_select.php) ^106^. Using these sets of coordinates, FitHiChIP loops that overlap bulk H3K27ac peaks and scATAC-peaks in astrocytes were subset to those that contained a promoter in one anchor and a SE in the other anchor. Binomial smooth plots were generated as in ^107^.

### Heritability analyses

#### MAGMA

Summary statistics from ^22^ were uploaded to FUMA (v1.5.6) ^108^ web server (https://fuma.ctglab.nl). Gene-level Z-scores were calculated using SNP2GENE with the “Perform MAGMA” function (MAGMA v1.08) and default parameter settings. The reference panel population was set to “1000G Phase3 EUR”. The MHC region was excluded due to its unusual genetic architecture and linkage disequilibrium. MAGMA Z-scores were then used for downstream analyses as described in the **Supplementary Note**.

#### Stratified LD score regression

To partition SNP-heritability, we used Stratified LD score regression (S-LDSC) (v1.0.1) ^26^, which assesses the contribution of gene expression programs to disease heritability. First, for analysis of astrocyte-identity genes, we computed (within the BA46 region only), a Wilcoxon rank sum test on a per-gene basis using presto (v1.0.0) ^109^ between astrocytes and all other cell-types; for analysis of astrocyte-activity genes (SNAP-a), we sorted all genes expressed in astrocytes by their SNAP-a loadings and took the top 2,000 genes. We then converted each gene set into annotations for S-LDSC by extending the window size to 100kb (from the transcription start site and transcription end site), and ordered SNPs in the same order as the .bim file (from phase 3 of the 1000 Genomes Project ^110^) used to calculate the LD scores. We then computed LD scores for annotations using a 1 cM window and restricted the analysis to Hapmap3 SNPs. We excluded the major histocompatibility (MHC) region due to both its high LD and high gene density. We used LD weights calculated for HapMap3 SNPs for the regression weights. We then jointly model the annotations corresponding to our gene expression program, as well as all protein coding genes, and the baseline model (baseline model v1.2). We tested for enrichment of SNP heritability on the traits listed below. The LDSC script, “munge_sumstats.py” was used to prepare the summary statistics files. We used the resultant p-values, which reflect a one-sided test that the coefficient (*τ*) is greater than zero, as a determinant as to whether our cell type gene expression programs are enriched for SNP-heritability of a given trait ^111^.

We used summary statistics from the following studies in **Extended Data Fig. 26**: ADHD ^112^, ALS ^113^, Alzheimer’s disease ^114^, age of smoking initiation ^115^, autism ^116^, bipolar disorder (all, type I, and type II) ^117^, cigarettes per day ^115^, educational attainment ^118^, epilepsy (all, focal, generalized) ^119^, height ^120^, IQ ^121^, insomnia ^122^, neuroticism ^123^, OCD ^124^, schizophrenia ^22^, PTSD^125^, risk ^126^, subjective well-being ^127^, smoking cessation ^115^, smoking initiation ^115^, Tourette’s ^128^, ulcerative colitis ^129^.

### Polygenic risk scores

Clumped summary statistics for schizophrenia (from ^22^) across 99,194 autosomal markers were downloaded from the Psychiatric Genomics Consortium portal (file PGC3_SCZ_wave3_public.clumped.v2.tsv). After liftOver of markers to GRCh38 using custom tools, 99,135 markers were available for scoring. We processed the output data from the MoChA imputation workflow ^58,59^ using BCFtools (v1.16) and the MoChA score (v2022-12-21) ^58,59^ workflow (https://github.com/freeseek/score) to compute schizophrenia polygenic scores across all 2,413 imputed samples from the McLean cohort.

#### C4

##### MetaGene discovery

Genes that have high sequence homology are typically difficult to capture by standard UMI counting methods. Reads from these regions map to multiple locations in the genome with low mapping quality, and are ignored by many gene expression algorithms. MetaGene discovery leverages that high sequence similarity by looking for UMIs that consistently map to multiple genes at low mapping quality consistently across many cells.

Each UMI is associated with a single gene if at least one read from the UMI uniquely maps to a single gene model. If all reads are mapped at low quality to multiple genes, then assignment of that UMI to a specific gene model is ambiguous, and that UMI is associated with all gene models. By surveying a large number of cells, a set of gene families are discovered where UMIs are consistently associated with sets of genes. This discovery process finds expected sets of gene families with high sequence homology directly from the mapping, such as *C4A/C4B*, *CSAG2/CSAG3*, and *SERF1A/SERF1B*.

These UMIs are then extracted in the counts matrix as a joint expression of all genes in each set. We prefer to calculate expression as the joint expression of all genes in the set because the priors in the data prevent confidently distributing these ambiguous UMIs. For example, *C4A* and *C4B* have very few UMIs that map uniquely to either gene in the set (8 UMIs, < 0.5% of all UMIs captured for this set of genes), which is weak prior to proportionally assign ambiguous UMIs to the correct model.

This approach was validated for *C4* expression by generating a reference genome that contained only one copy of *C4*. This allowed each UMI to map uniquely to the single remaining copy of the gene using standard tools. The custom reference approach and joint expression of *C4A/C4B* via the metagene approach was concordant in 15,664 of 15,669 cells tested **(Extended Data Fig. 28c)**.

##### Imputation of *C4* structural variation

Phased copy number calls for structural features of the *C4* gene family were obtained by imputation using Osprey, a new method for imputing structural variation. The total copy number of *C4* genes, the number of copies of *C4A* and *C4B*, and the copy number of the polymorphic HERV element that distinguishes long from short forms of *C4* ^29^ were imputed into the McLean cohort using a reference panel based on 1000 Genomes ^62^.

An imputation reference panel was constructed for GRCh38 using 2604 unrelated individuals (out of 3202 total) from 1000 Genomes. SNPs were included in the reference panel if (a) they were within the locus chr6:24000000-34000000 but excluding the copy-number variable region chr6:31980001-32046200 and (b) they were not multi-allelic and (c) they had an allele count (AC) of at least 3 when subset to the 2604 reference individuals.

The imputation reference panel was merged with genotypes for the McLean cohort obtained from the GSA genotyping arrays. Markers not appearing in both data sets were dropped and the merged panel was phased with SHAPEIT4 (v4.2.0) ^57^ using default parameters plus “--sequencing” and the default GRCh38 genetic map supplied with SHAPEIT.

Reference copy numbers for the *C4* structural features on GRCh38 were obtained for the 3202 1000 Genomes samples using a custom pipeline based on Genome STRiP (v2.0) ^130^. Source code for this pipeline is available on Terra (http://app.terra.bio) ^131^. Briefly, the pipeline uses Genome STRiP to estimate total *C4* copy number and HERV copy number from normalized read depth-of-coverage, then estimates the number of copies of *C4A* and *C4B* using maximum-likelihood based on reads that overlap the *C4* active site (coordinates chr6:31996082-31996099 and chr6:32028820-32028837). These copy number genotypes were then subset to the 2604 unrelated individuals.

The structural features were imputed into the merged imputation panel using Osprey (v0.1-9) ^132,133^ by running ospreyIBS followed by osprey using default parameters plus “-iter 100”, the SHAPEIT4 genetic map for GRCh38 chr6, and a target genome interval of chr6:31980500-32046500.

The output from Osprey was post-processed using a custom R script (refine_C4_haplotypes.R) that enforces constraints between the copy-number features and recalibrates the likelihoods considering only “possible” haplotypes. The enforced constraints are that the *C4A*+*C4B* copies must equal total C4 and that the HERV copy number must be less than or equal to *C4* copy number.

### Source data and visualization

In addition to the software cited above, we used Color Oracle (v1.3) ^134,135^ as well as the following packages to prepare the source data and figures in this manuscript.

Python (v3.8.3): matplotlib (v3.5.2) ^136^, seaborn (v0.10.1) ^137^.

R (v4.1.3): cluster (v2.1.2) ^138^, ComplexHeatmap (v2.10.0) ^139,140^, data.table (v1.14.8) ^141^, DescTools (v0.99.48) ^142^, dplyr (v1.1.2) ^143^, gdata (v2.19.0) ^144^, ggforce (v0.4.1) ^145^, ggplot2 (v3.4.2) ^146^, ggpmisc (v0.5.3) ^147^, ggpointdensity (v0.1.0) ^148^, ggpubr (v0.5.0) ^149^, ggrastr (v1.0.2) ^150^, ggrepel (v0.9.3) ^151^, grid (v4.1.3) ^152^, gridExtra (v2.3) ^153^, gtable (v0.3.3) ^154^, matrixStats (v0.63.0) ^155^, pheatmap (v1.0.12) ^156^, plyr (v1.8.8) ^157^, purrr (v1.0.1) ^158^, RColorBrewer (v1.1-3) ^159^, readxl (v1.4.2) ^160^, reshape2 (v1.4.4) ^161^, scales (v1.2.1) ^162^, splitstackshape (v1.4.8) ^163^, stats (v4.1.3) ^152^, stringi (v1.7.12) ^164^, stringr (v1.5.0) ^165^, tidyr (v1.3.0) ^166^, viridis (v0.6.2) ^167^.

## DATA AVAILABILITY

Sequencing data generated in this study and processed sequencing files are available through the Neuroscience Multi-omic Data Archive (NeMO) (RRID:SCR_016152) at https://assets.nemoarchive.org/dat-bmx7s1t. The data are available under controlled use conditions set by human privacy regulations. To access the data, the requester must first create an account in DUOS (https://duos.broadinstitute.org) using their institutional email address. The Signing Official from the requester’s institution must also register in DUOS to issue the requester a Library Card Agreement. The requester will then need to fill out a Data Access Request through DUOS, which will be reviewed by the Broad Institute’s Data Access Committee. Once a request is approved, NeMO will be notified to authorize access to the data. Processed expression data can also be queried using an interactive public web interface that we created (https://dlpfc.mccarrolllab.org/app/dlpfc). Source data with anonymized donor IDs are provided with this paper.

The following publicly available datasets were also analyzed: ProteomeXchange Dataset PXD026491 ^82^ and Gene Expression Omnibus Series GSE147672 ^99^.

## CODE AVAILABILITY

Software and core computational analysis to align and process sequencing reads and perform donor assignment are freely available: https://github.com/broadinstitute/Drop-seq. Published or publicly available software, tools, algorithms, and packages are cited with their version numbers in the text and Reporting Summary. Other custom code is available upon request from the corresponding authors.

## ACKNOWLEDGEMENTS

This work was supported by the Broad Institute’s Stanley Center for Psychiatric Research, the Simons Collaboration on Plasticity and the Aging Brain, the National Institute of Mental Health (grants U01MH115727 to S.A.M. and P50MH115874 Project 5 to S.B.), and the National Human Genome Research Institute (grant T32 HG002295 to N.K.). Human tissue was obtained from the NIH NeuroBioBank. We thank Heather de Rivera, Rhea Kohli, and Gabriel Lind for technical assistance; Rebecca Hodge for advice on myelin removal; Trygve Bakken and Nik Jorstad for advice on glutamatergic neuron subtype classification; Frank Koopmans for SynGO analysis scripts; and members of the McCarroll lab and the Stanley Center for helpful advice and discussion. We thank Mehrtash Babadi, Kris Dickson, Marta Florio, Steven Hyman, Yong Hoon Kim, Ajay Nadig, Ralda Nehme, Chris Patil, Elise Robinson, Morgan Sheng, and Matthew Tegtmeyer for helpful comments on manuscript drafts. Finally, we would like to thank the brain tissue donors and their families, without whom this study would not be possible.

## AUTHOR CONTRIBUTIONS

E.L., S.A.M., and S.B. designed the study. E.L., M.G., N.R., and S.A.M. developed and evaluated experimental strategies for snRNA-seq from pooled human brain tissue. E.L., M.G., N.R., A.L., and C.D.M. prepared and dissected tissue, performed snRNA-seq, and prepared sequencing libraries. E.L., J.N., M.G., and S.A.M. performed sequencing, alignment, and quality-control analyses. E.L., J.N., A.W., and S.A.M. developed analysis pipelines. E.L. and S.A.M. analyzed the data with input from S.A.M., S.B., J.N., and N.K. B.H. performed analyses of *C4*. G.G. performed imputation and calculated polygenic risk scores. J.S.V. and S.B. provided tissue donor metadata. S. Gerges calculated MAGMA Z-scores and performed heritability enrichment analyses with S-LDSC. S.K. developed the scPred analysis pipeline and the RNA-expression web resource. S. Ghosh developed the pySCENIC analysis pipeline. J.M.E., K.F., and S.B. evaluated and provided tissue for snRNA-seq experiments. D.M. contributed to analysis pipelines. L.S. contributed to tissue sample management and standardization of the single-nucleus library preparation and sequencing protocol. A.N., M.H., and K.I. contributed to project management and sequencing. E.L., S.A.M., and S.B. wrote the paper with input from co-authors.

## ADDITIONAL INFORMATION

Supplementary Information is available for this paper.

Correspondence and requests for materials should be addressed to Steven A. McCarroll, Sabina Berretta, or Emi Ling.

## SUPPLEMENTARY NOTE

### Synaptic Neuron-Astrocyte Program (SNAP) in the genetics of schizophrenia

#### Cell-identity gene expression and schizophrenia genetics: replication of earlier results

Many earlier studies ^1–3^ have found that genes most strongly expressed by neurons relative to other CNS cell types, but not genes most strongly expressed by astrocytes or other glia, are enriched for the genes implicated by human-genetic studies in schizophrenia. We first replicated these findings using the data from the current experiments. Genes that were preferentially expressed in neurons (as defined by the criteria used in the earlier studies) exhibited enrichment for schizophrenia-risk genes and alleles by a variety of analysis methods, but genes that were preferentially expressed in astrocytes did not.

For example, the following is an analysis of common-variant association signals (MAGMA gene-level Z-scores) versus these sets of “cell-type preferentially expressed genes” (as defined by the methods of earlier work and applied to the current data), for neurons and astrocytes. For neurons, for example, this gene-set comprises the 2,000 genes for which neurons exhibit the highest quantitative expression levels relative to other cell types. As expected, we see strong significance for neurons but not for astrocytes:

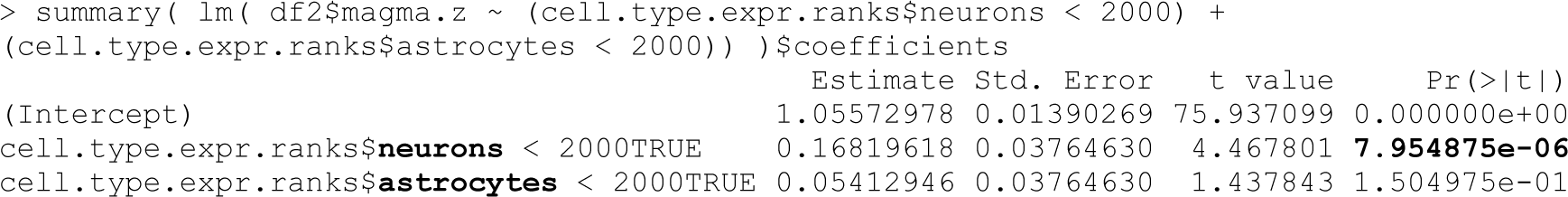

(Here we used a single, composite “neuronal” set of expression values for the analysis, but we had very similar results when we used specific types or subtypes of cortical neurons, reflecting the strong correlation among their gene-expression levels.)

Thus, these well-established and expected relationships are also visible in the current, human cell-type-specific expression data.

#### Cellular programs and schizophrenia genetics

The above analysis, like other analyses to date, treats cell types as fixed levels of cell-identity gene expression, rather than as dynamic biological entities that utilize gene expression in ways whose variation is also meaningful. Cell-identity gene expression actually tells us little about SNAP-a: of the 500 genes most strongly recruited by SNAP-a, more than 90% are also robustly expressed in neurons and/or other glia of various types; less than half (203 of 500) are most strongly expressed in astrocytes, reflecting that biological functions such as synaptic adhesion and neurotransmitter uptake are also performed by neurons. Rather, it is the close transcriptional co-regulation of these genes in astrocytes by SNAP-a that appears to strongly distinguish astrocytes from neurons **(Fig. 3k)**.

Cell types would ideally be considered, not only in terms of static cell-identity gene expression, but by their repertoires of *dynamic* transcriptional responses, such as SNAP-a, the set of astrocyte gene-expression changes that appear to be implemented in tandem with synaptic gene-expression changes in neurons (SNAP-n). To do so, we started with the 500 genes whose expression is most strongly recruited by SNAP-a (as defined by the gene loadings on this latent factor, and reflecting the fraction of their single-cell expression variance that is explained by the latent factor or cell state). We first asked whether these 500 genes are enriched for strong (genome-wide significant) associations to common and rare variants in schizophrenia. These SNAP-a-defined genes were 14 times more likely (than other protein-coding genes) to reside at genomic loci implicated by common genetic variation in schizophrenia (p = 5 × 10^−25^, 95% confidence interval: 8.7-24, by logistic regression, based on this SNAP-a-500 gene set containing 26 of the 98 protein-coding genes at 105 loci at which associated haplotypes involved SNPs in just 1-2 genes). These genes were also 7 times more likely (than other protein-coding genes) to have strong evidence from rare variants in schizophrenia (95% CI: 2.3–21, p = 5×10^−4^, by logistic regression, based on the SNAP-a-500 gene set containing 4 of the 32 genes implicated at FDR<0.05 by the SCHEMA Consortium or by rare, intragenic deletions). Note that SNAP-a was significant even in models in which “preferential expression in neurons” was a competing predictive factor:

Genes with common variation implicated in schizophrenia

(loci at which associated haplotypes involved SNPs in just 1-2 genes)

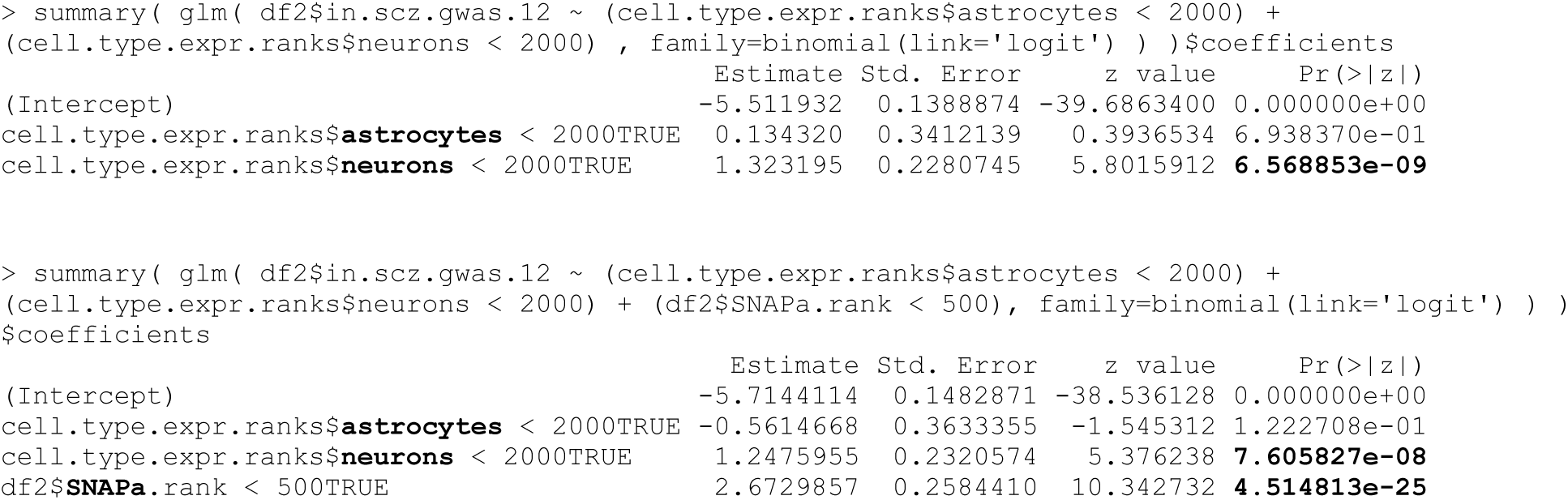

Genes with rare variation implicated in schizophrenia

(SCHEMA FDR<0.05 + *NRXN1*)

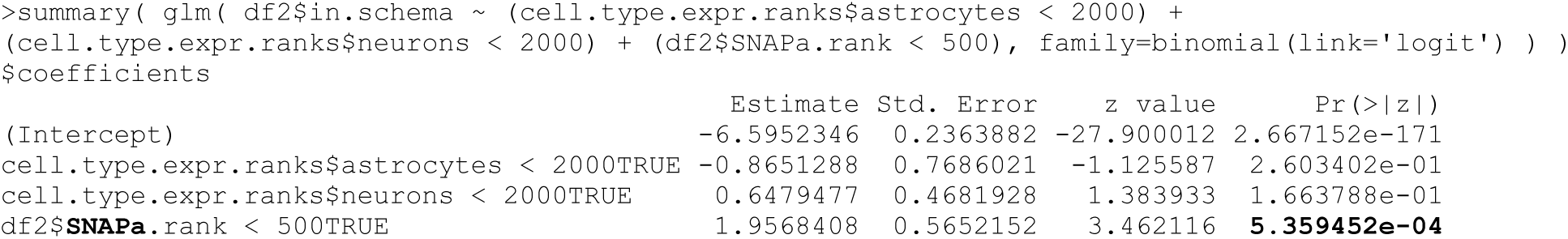

In the above analysis of genes implicated by rare variants, baseline expression in neurons was not significant. The implication of neuronally-expressed genes in the study by the SCHEMA Consortium used a different type of analysis, which used a Wilcoxon rank-sum test to evaluate whether SCHEMA genes had higher levels of neuron-preferential expression than other protein-coding genes did. In that analysis, in Figure S17 of the SCHEMA paper ^2^, about a third of the neuronal types tested yielded p-values less than 0.05, whereas no non-neuronal cell types did. We repeated this analysis with the data from the current study, with several subtypes of neurons yielding nominally significant results (p<0.05) but not astrocytes (p=0.64), in accordance with the earlier finding. When we applied an analogous analysis to the gene loadings for SNAP-a, it was highly significant (p = 8 × 10^−5^).

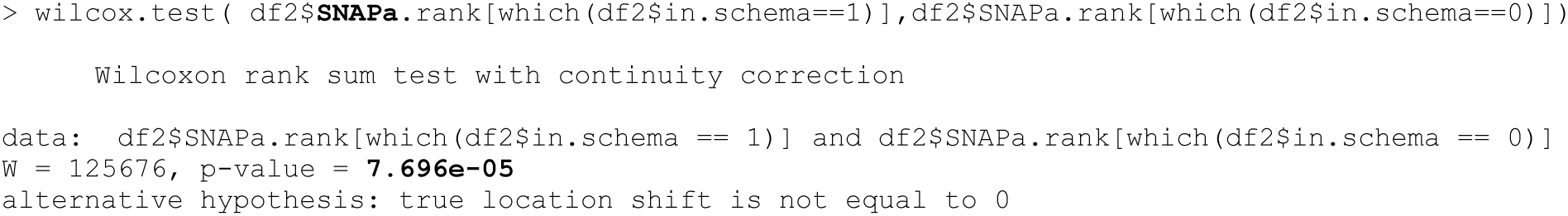

To evaluate, beyond these top genetic associations, whether common genetic variation in the genes recruited by SNAP-a contributes more broadly to schizophrenia risk, we further utilized the gene-level association statistics provided by MAGMA analysis ^1,4^, which evaluates, for every gene, the tendency of common patterns of genetic variation (as identified by principal components analysis) to have elevated levels of association. To integrate across these more-subtle genomic signals, we also used a larger number of genes prioritized by SNAP-a. The 2,000 genes whose expression is most strongly recruited by SNAP-a had elevated MAGMA z-scores for association to schizophrenia (p < 2 × 10^−20^), while astrocyte-identity gene expression did not (p = 0.53).

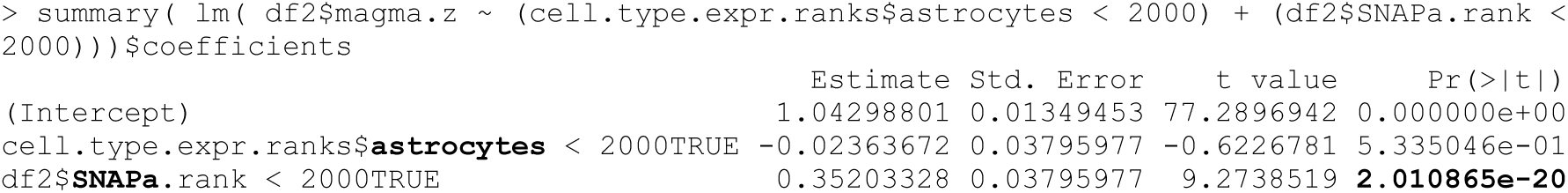

Since the number of genes in the SNAP-a gene set is a somewhat arbitrary parameter of this analysis, we explored the relationship of this enrichment to the gene depth (on the SNAP-a-ranked gene list) used in analysis. The results for eight gene depths are summarized in the table below. Genetic signals were most strongly concentrated at the top of the SNAP-a gene list (as seen by the regression coefficient estimate, first column); however, concentration was still present at greater gene depths, and the statistical significance of the enrichment (as estimated by the test statistic, third column) increased in more-inclusive analyses up to about 2,000 genes, at which point it began to drop.

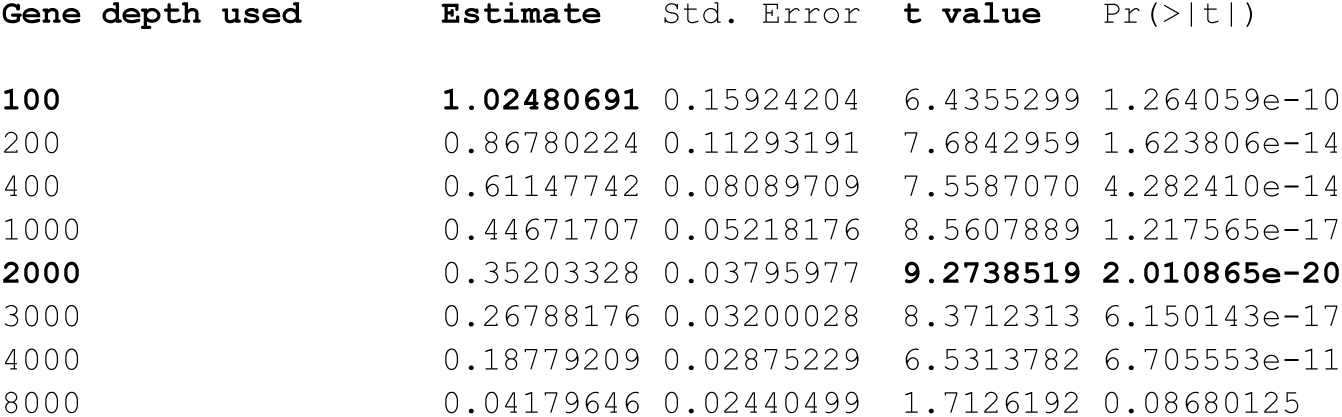

This relationship can also be recognized visually in a plot of MAGMA z-score vs. genes ordered by their SNAP-a gene loadings, which suggests that enrichment is strongest among the genes ranked most highly by SNAP-a (far left on plot) gene list but that enrichment continues, albeit more modestly, over the top 2,000 or so genes.

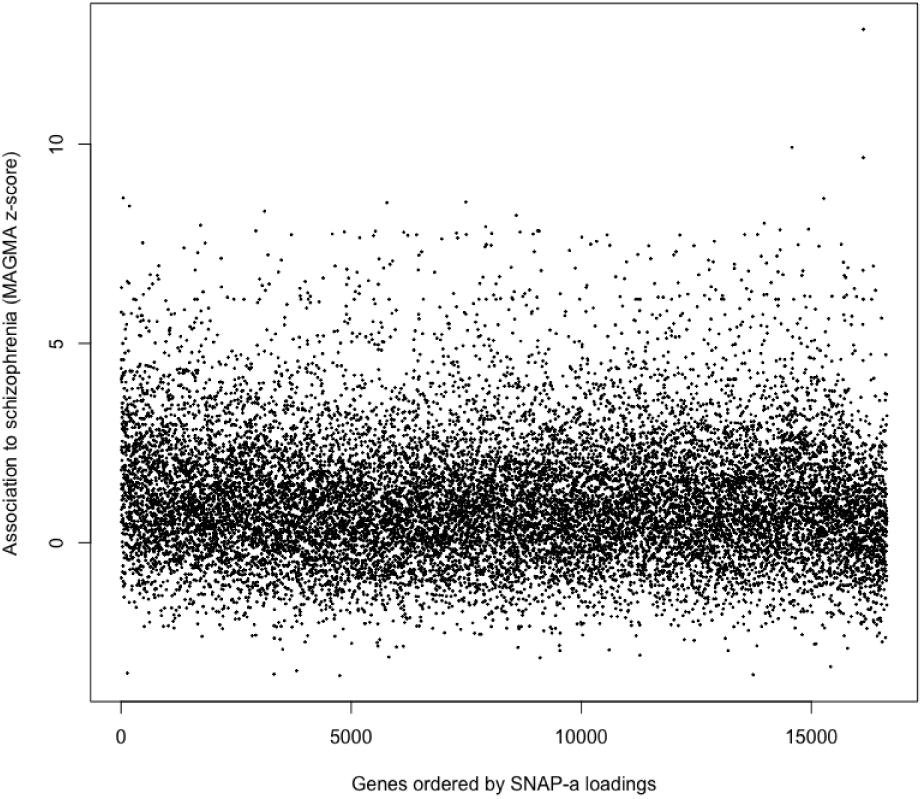

We also included neuronal-identity gene expression (as defined by the method used in the earlier studies) and SNAP-n-recruited genes in the regression analysis, as independent and competing predictive factors. All three were significant in a joint analysis, and the signal for SNAP-a genes was not attenuated by the inclusion of the two neuronal gene sets:

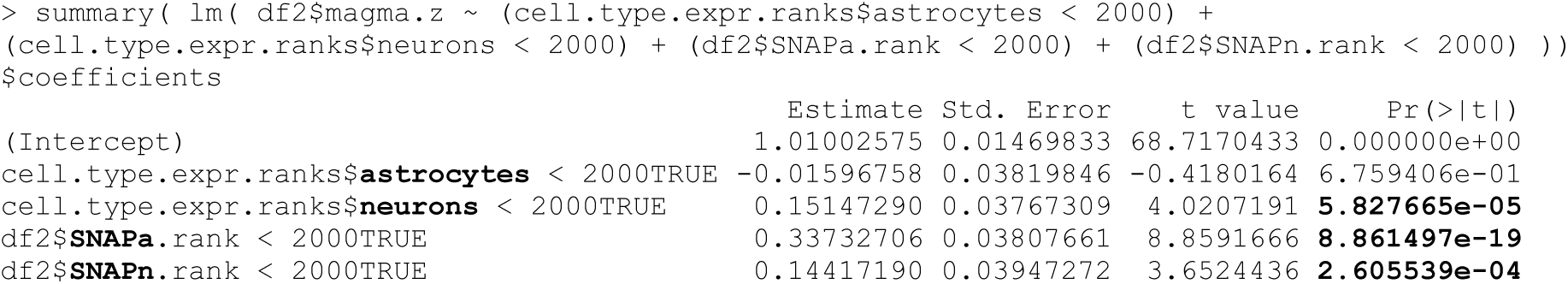

In the above result, both SNAP-n genes and neuronally-preferentially-expressed genes contributed independently to explaining gene-schizophrenia associations (MAGMA z-statistics), suggesting that – in neurons as in astrocytes – information about dynamic gene-expression programs can provide additional information beyond the information provided by cell-identity gene expression.

Finally, we used LD score regression ^5^ to evaluate per-SNP heritability enrichment across 27 brain phenotypes. Baseline astrocyte-identity gene expression (top 2,000 genes) did not exhibit heritability enrichment for any of the 27 brain phenotypes tested **(Extended Data Fig. 26a)**. SNAP-a (most strongly recruited 2,000 genes) exhibited per-SNP heritability enrichment (p = 4 × 10^−5^) for schizophrenia, nominal significance (p < 0.01) for smoking cessation and autism, and was not significant for the other 24 phenotypes tested **(Extended Data Fig. 26b)**.

## SUPPLEMENTARY TABLES

**Supplementary Table 1. Summary of human tissue donor metadata.**

Donor metadata table. Sample details include sex, age, post-mortem interval (PMI, when available), schizophrenia case-control status, and inclusion in experimental batches.

**Supplementary Table 2. Donor expression levels and gene loadings for latent factors.**

Tables of donor expression levels and genes-by-cell-types loadings for each of the 10 latent factors inferred by PEER.

**Supplementary Table 3. Regression analysis of LF4 donor expression levels.**

Joint regression analysis of LF4 donor expression levels with age, sex, and schizophrenia case-control status as independent variables.

**Supplementary Table 4. Gene set enrichment analysis (GSEA) results for LF4 by cell type.**

Tables of gene sets enriched in each cell type’s component of LF4 (at FDR < 0.05) from a preranked gene set enrichment analysis (GSEA) using LF4 gene loadings.

**Supplementary Table 5. Gene set enrichment analysis (GSEA) results for latent factors enriched in astrocytes.**

Tables of gene sets enriched in astrocyte latent factors discovered by cNMF (at FDR < 0.15) from a preranked gene set enrichment analysis (GSEA) using gene loadings for each factor.

**Supplementary Table 6. Donor expression levels and gene loadings for SNAP-a.**

Tables of donor expression levels (mean cell scores by donor) and gene loadings for SNAP-a (astrocyte latent factor 2 inferred by cNMF).

**Supplementary Table 7. Donor expression levels and gene loadings for SNAP-n.**

Tables of donor expression levels (mean cell scores by donor) and gene loadings for SNAP-n (L5 IT glutamatergic neuron latent factor 6 inferred by cNMF).

**Supplementary Table 8. Gene set enrichment analysis (GSEA) results for SNAP-n.**

Table of gene sets enriched in SNAP-n (at FDR < 0.15) from a preranked gene set enrichment analysis (GSEA) using gene loadings for SNAP-n.

**Supplementary Table 9. Genes in selected gene sets.**

Table of genes in selected gene sets used in analyses. Descriptions of selected gene sets are in Methods.

## EXTENDED DATA FIGURES

**Extended Data Figure 1.**
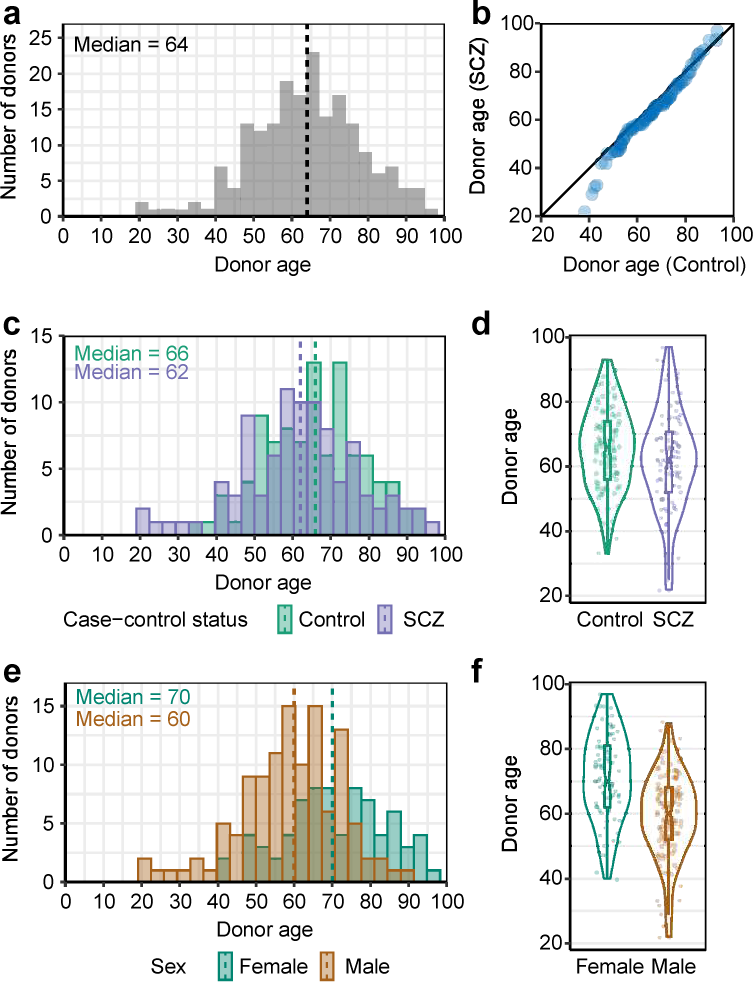
Ages of brain tissue donors. **a,** Distribution of the ages of brain donors (*n* = 191 donors). **b,** Distributions of donors’ ages by schizophrenia status, displayed as a quantile-quantile plot that compares ages of unaffected control donors (*n* = 97 donors) to ages of donors with schizophrenia (*n* = 94 donors). **c-d,** Distributions of donors’ ages separated by schizophrenia status (*n* = 97 unaffected and 94 affected), displayed as **(c)** histograms and **(d)** violin plots. **e-f,** Distributions of donors’ ages, separated by sex (*n* = 75 women and 116 men), displayed as **(e)** histograms and **(f)** violin plots. Note that while female brain donors are on average older than male donors, expression of SNAP (LF4) did not associate with sex in either a naive or age-adjusted analysis **(Extended Data Fig. 9d-e)**, nor in a simultaneous regression on age, sex, and schizophrenia affected/unaffected status **(Supplementary Table 3)**.

**Extended Data Figure 2.**
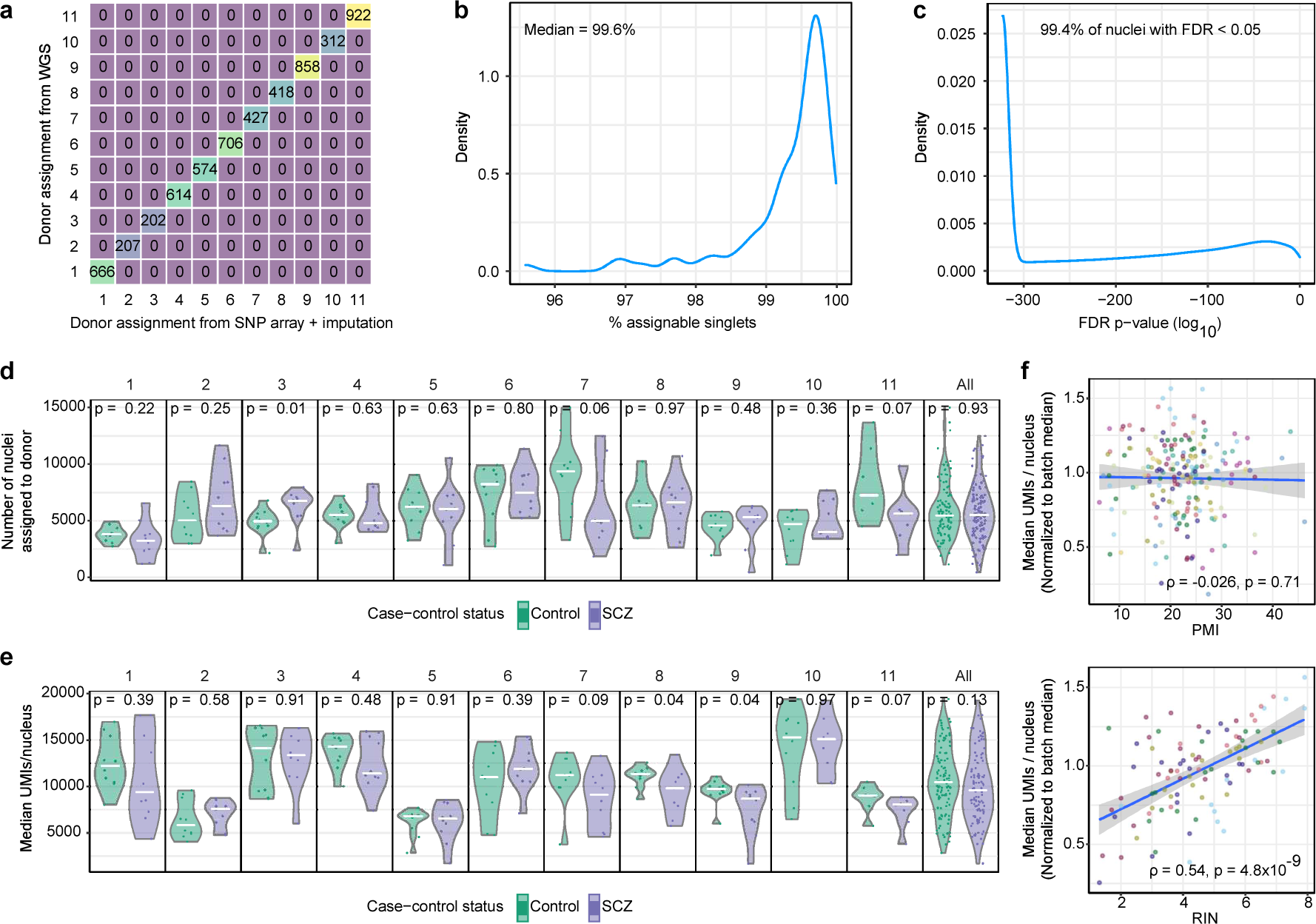
Single-donor assignment and sequencing metrics. **a,** Validation of the computational assignment of nuclei to individual brain donors whose genomes have been analyzed (individually) by SNP array-genotyping plus imputation. The matrix displays the concordance of single-donor assignment between whole-genome sequencing (WGS) (y-axis) and SNP array + imputation (x-axis) for a pilot set of 11 donors whose genomes were analyzed by both methods. (Accuracy of donor assignment when WGS data are available has been previously shown by ^7^.) Each row/column corresponds to one of the 11 donors, and each entry in the table displays the number of nuclei that were assigned to a given donor (at a false discovery rate of 0.05). **b,** Density plot showing the fraction of all nuclei that were determined to be “singlets” (containing alleles from just one donor); *n* = 1,262,765 assignable singlets out of 1,271,830). **c,** Density plot showing donor-assignment likelihoods (as false discovery rates, on a log scale) for the 1,271,830 singlet nuclei. **d,** Number of nuclei assigned to each donor in each of 11 batches or (rightmost panel) across all batches, separated by schizophrenia case-control status (*n* = 10 controls and 10 schizophrenia cases per batch). P-values from a two-sided Wilcoxon rank-sum test comparing the affected to the unaffected donors are reported at the top of each panel. Central lines represent medians. **e,** Median number of UMIs ascertained per donor in each batch or (rightmost panel) across all batches, separated by schizophrenia case-control status (*n* = 10 controls and 10 schizophrenia cases per batch). P-values from a two-sided Wilcoxon rank-sum test comparing the affected to the unaffected donors are reported at the top of each panel. Central lines represent medians. **f,** Relationship of median UMIs/nucleus (normalized to the median value of the donors in each donor’s batch) to (top) post-mortem interval (PMI) and (bottom) RIN score (Spearman’s Colors represent different batches. Shaded regions represent 95% confidence intervals.

**Extended Data Figure 3.**
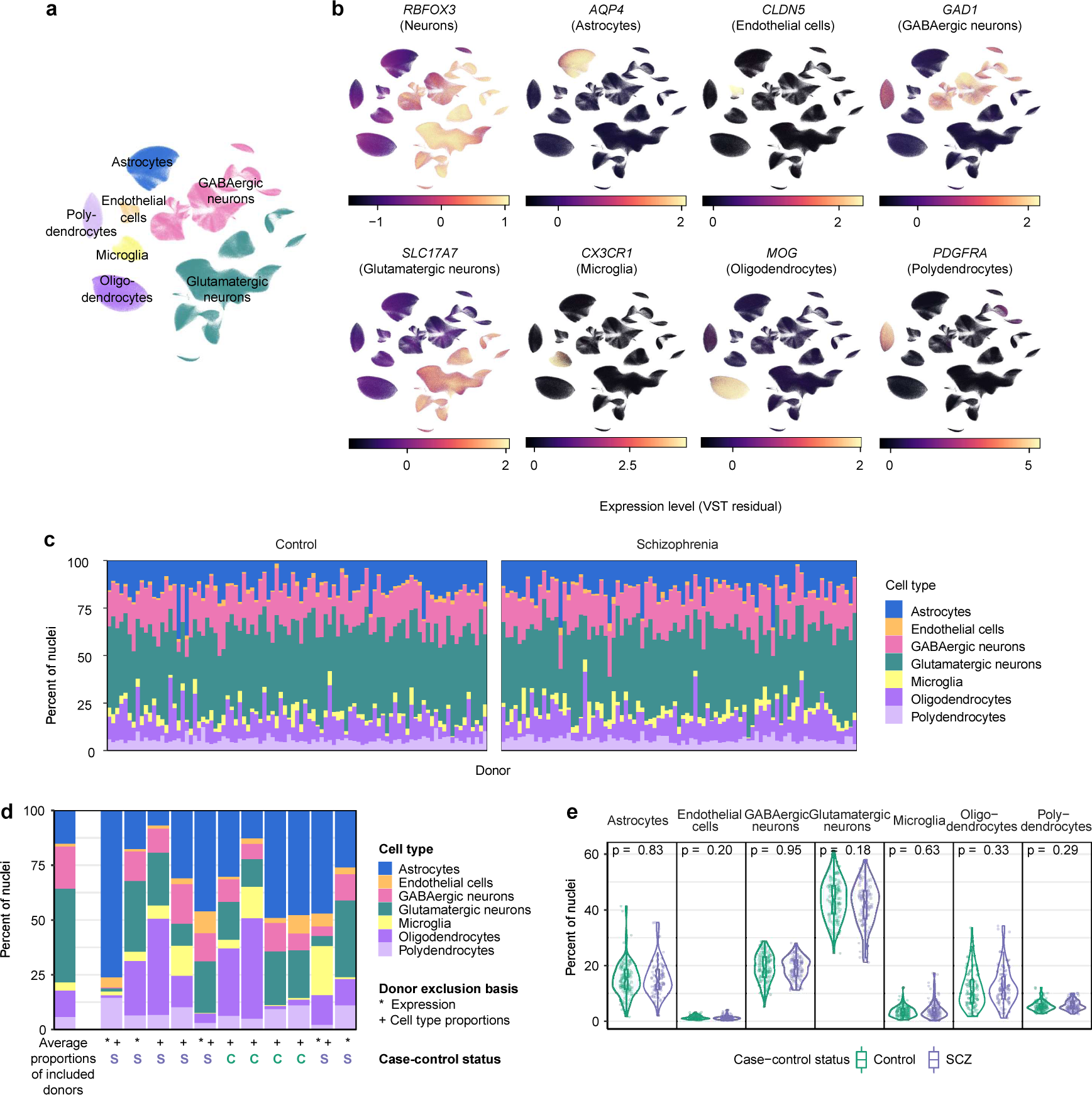
Cell-type classification and composition analysis. **a,** Two-dimensional projection of the RNA-expression profiles of the 1,218,284 nuclei analyzed from 191 donors, reproduced from **Fig. 1c**. Nuclei are colored by their assignments to the major cell types present in Brodmann area 46 (BA46). The same projection is used in panel **b**. **b,** Expression levels of canonical marker genes of cell types in BA46. Values represent Pearson residuals from variance stabilizing transformation (VST). **c,** Relative representation of each cell type among nuclei ascertained from each donor. Donors are ordered by their anonymized research IDs at the Harvard Brain Tissue Resource Center. **d,** Cell-type proportions detected in 11 donors whom we excluded from subsequent analyses, with the basis of exclusion (unusual cell-type proportions and/or expression profiles) indicated under each donor. For comparison, average cell-type proportions of the 180 donors included in subsequent analyses are displayed to the left (donors from panel **c**). **e,** Cell-type proportions ascertained in the BA46 tissue samples; data points are separated by schizophrenia status (*n* = 93 unaffected and 87 affected). P-values from a two-sided Wilcoxon rank-sum test comparing the affected to the unaffected donors are reported at the top of each panel. Box plots show interquartile ranges; whiskers, 1.5x the interquartile interval; central lines, medians; notches, confidence intervals around medians.

**Extended Data Figure 4.**
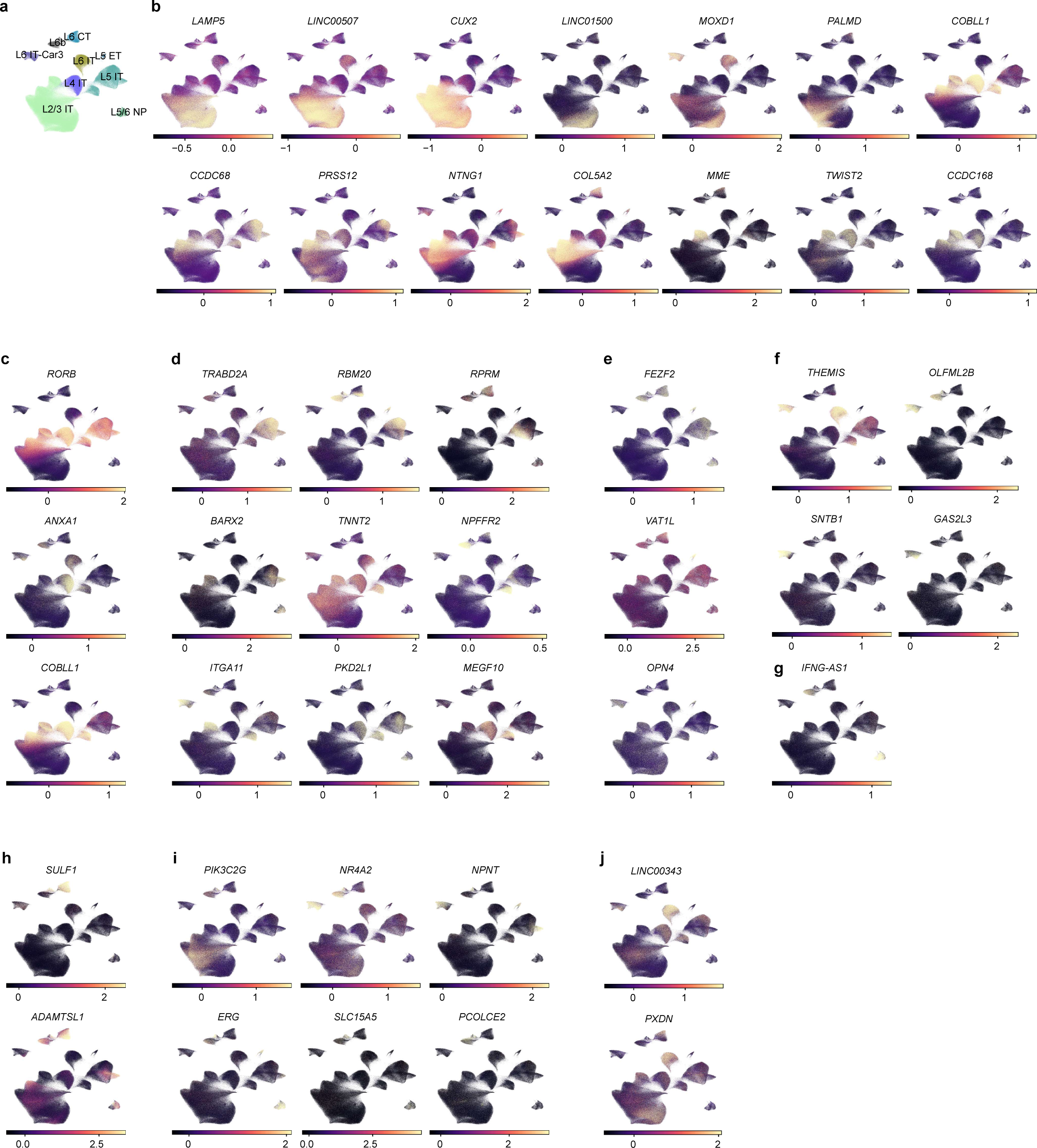
Expression of glutamatergic neuron-subtype marker genes. **a,** Two-dimensional projection of the RNA-expression profiles of 524,186 glutamatergic neuron nuclei, reproduced from **Fig. 1d**. Nuclei are colored by their assignments to subtypes of glutamatergic neurons using classifications from ^75^ and ^76^. The same projection is used in panels **b** to **j** below. **b-j** Expression levels of marker genes for subtypes of **(b)** L2/3 IT, **(c)** L4 IT, **(d)** L5 IT, **(e)** L5 ET, **(f)** L6 IT-Car3, **(g)** L5/6 NP, **(h)** L6 CT, **(i)** L6b, and **(j)** L6 IT glutamatergic neurons. Markers are from ^75^ or from transcriptomically similar subtypes in ^76^. Values represent Pearson residuals from variance stabilizing transformation (VST).

**Extended Data Figure 5.**
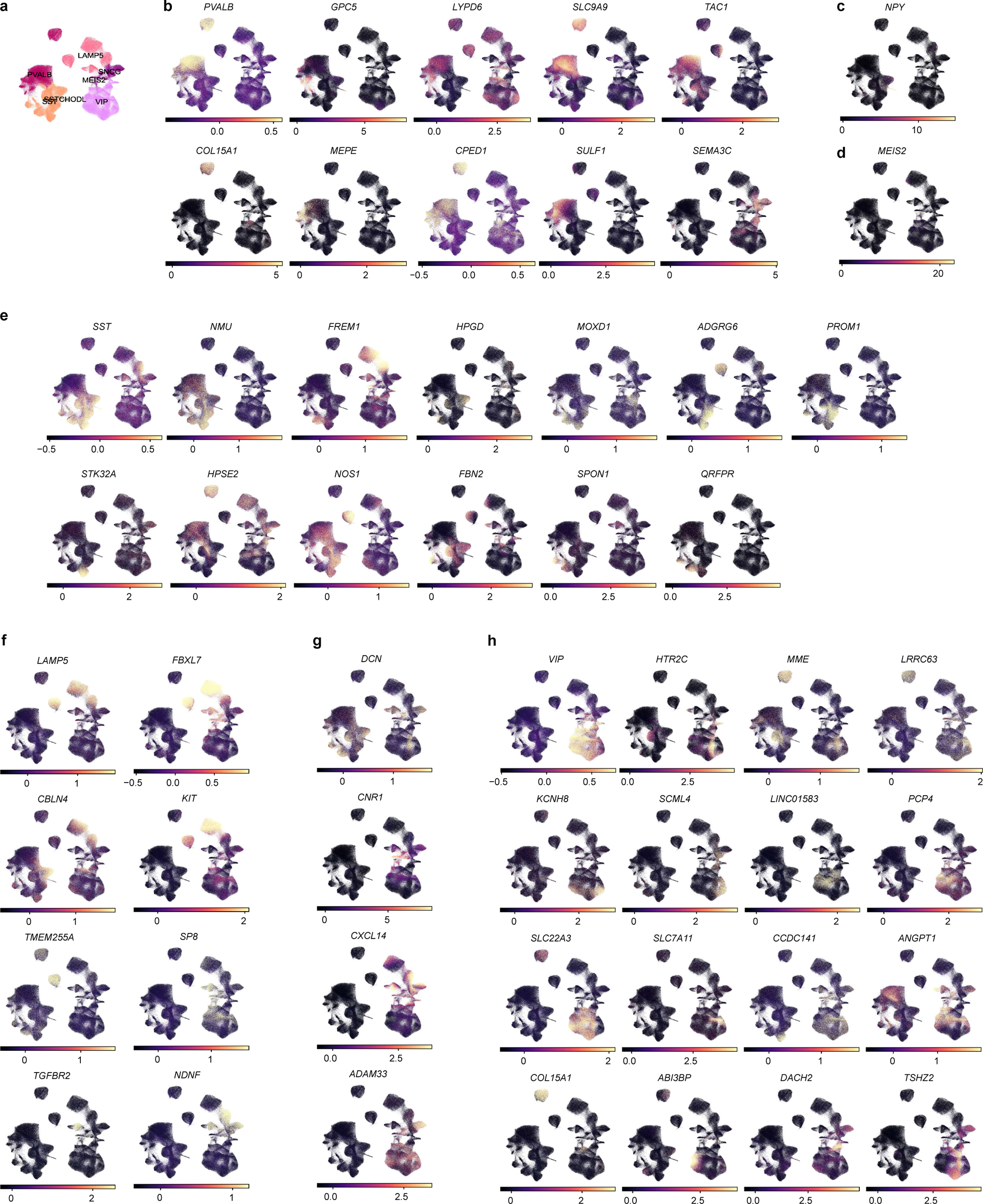
Expression of GABAergic neuron-subtype marker genes. **a,** Two-dimensional projection of the RNA-expression profiles of 238,311 GABAergic neuron nuclei, reproduced from **Fig. 1e**. Nuclei are colored by their assignments to subtypes of GABAergic neurons using classifications from ^75^ and ^76^. The same projection is used in panels B to H below. **b-h,** Expression levels of marker genes for subtypes of **(b)** PVALB, **(c)** SST-CHODL, **(d)** MEIS2, **(e)** SST, **(f)** LAMP5, **(g)** SNCG, and **(h)** VIP GABAergic neurons. Markers are from ^75^ or from transcriptomically similar subtypes in ^76^. Values represent Pearson residuals from variance stabilizing transformation (VST).

**Extended Data Figure 6.**
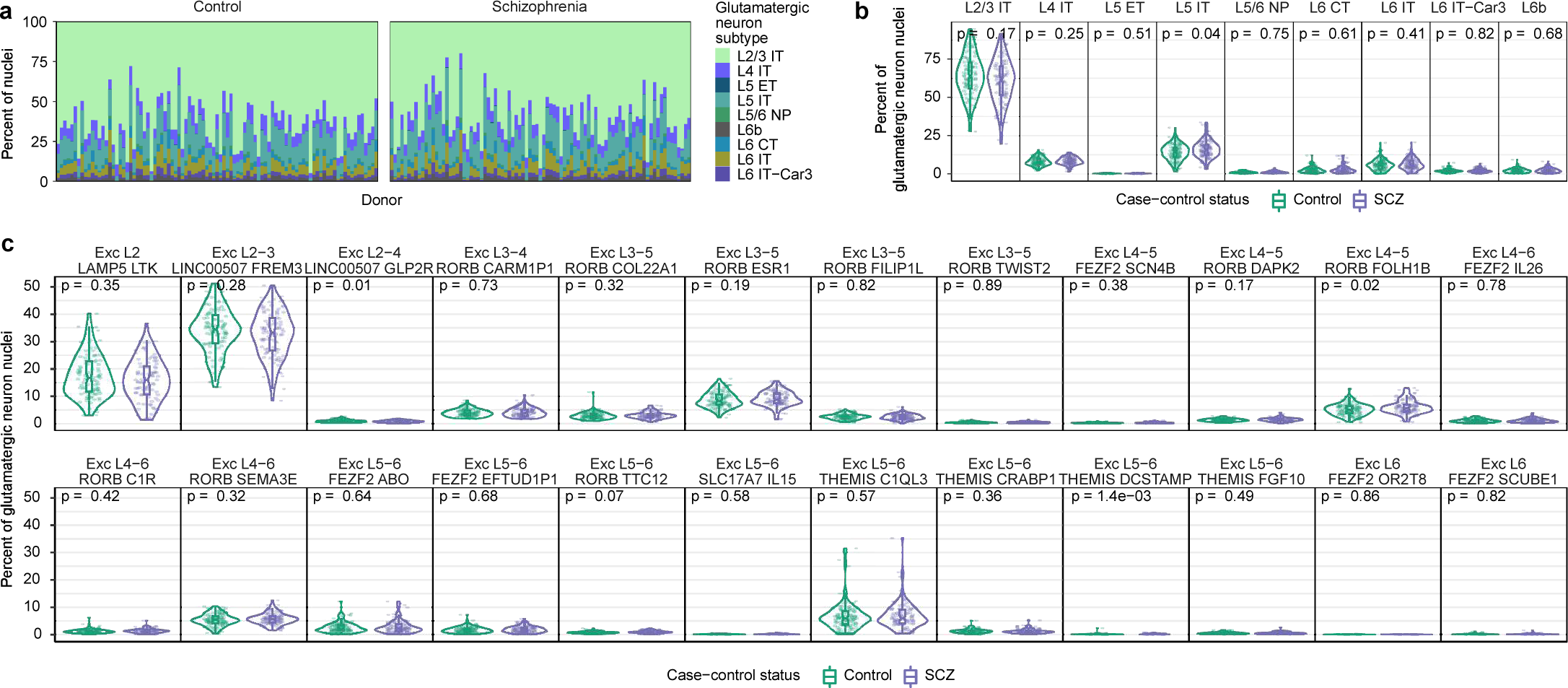
Glutamatergic neuron-subtype composition analysis across donors. **a,** Relative representation of each glutamatergic neuron subtype among nuclei ascertained from each donor. Donors are ordered by their anonymized research IDs at the Harvard Brain Tissue Resource Center. **b-c,** Proportions of **(b)** glutamatergic neuron subtypes and **(c)** subtypes of these subtypes (defined in ^75^) by schizophrenia status (*n* = 93 unaffected and 87 affected). P-values from a two-sided Wilcoxon rank-sum test comparing the affected to the unaffected donors are reported at the top of each panel. Box plots show interquartile ranges; whiskers, 1.5x the interquartile interval; central lines, medians; notches, confidence intervals around medians.

**Extended Data Figure 7.**
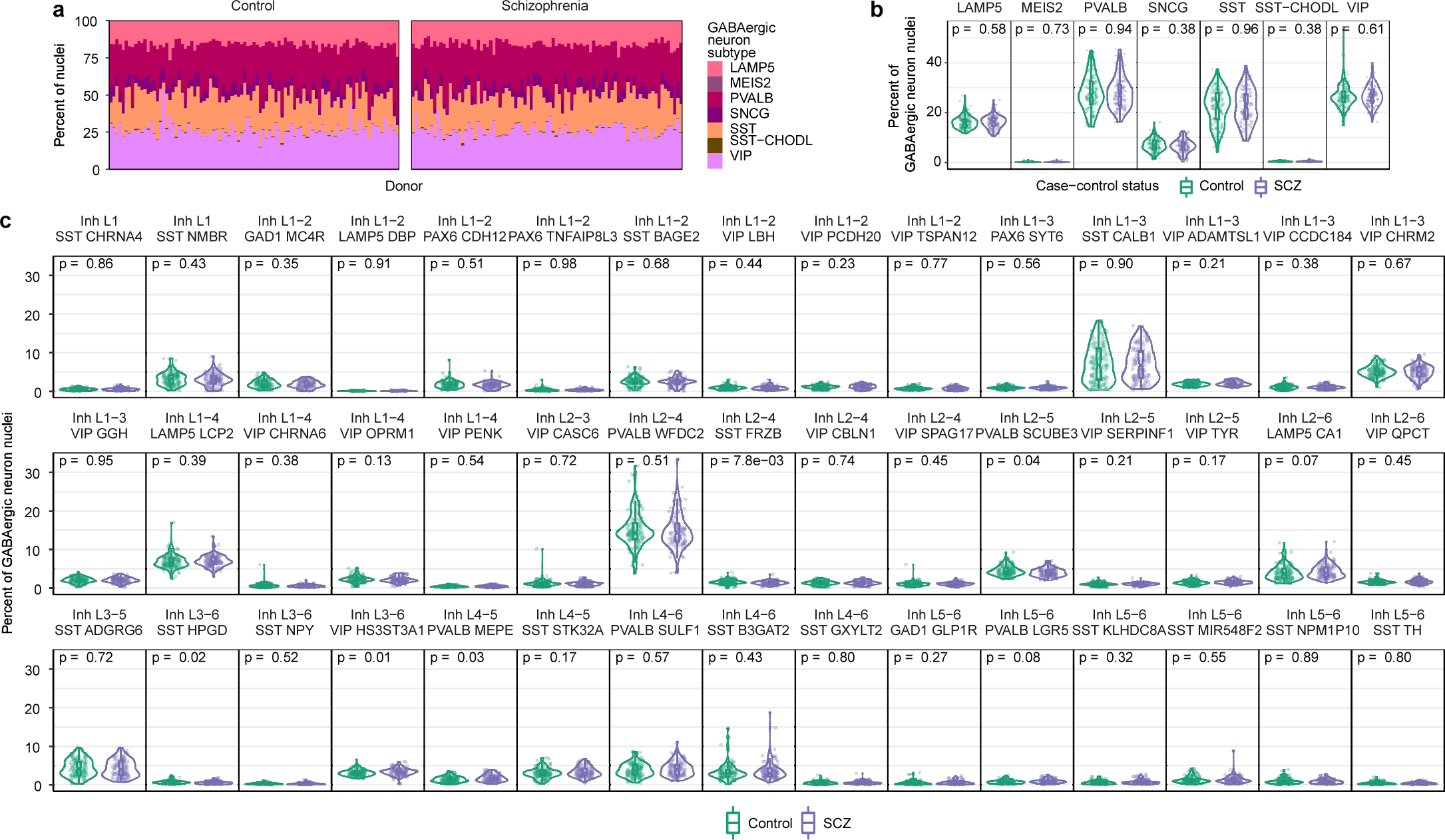
GABAergic neuron-subtype composition analysis across donors. **a,** Relative representation of each GABAergic neuron subtype among nuclei ascertained from each donor. Donors are ordered by their anonymized research IDs at the Harvard Brain Tissue Resource Center. **b-c,** Proportions of **(b)** GABAergic neuron subtypes and **(c)** subtypes of these subtypes (defined in ^75^) by schizophrenia status (*n* = 93 unaffected and 87 affected). P-values from a two-sided Wilcoxon rank-sum test comparing the affected to the unaffected donors are reported at the top of each panel. Box plots show interquartile ranges; whiskers, 1.5x the interquartile interval; central lines, medians; notches, confidence intervals around medians.

**Extended Data Figure 8.**
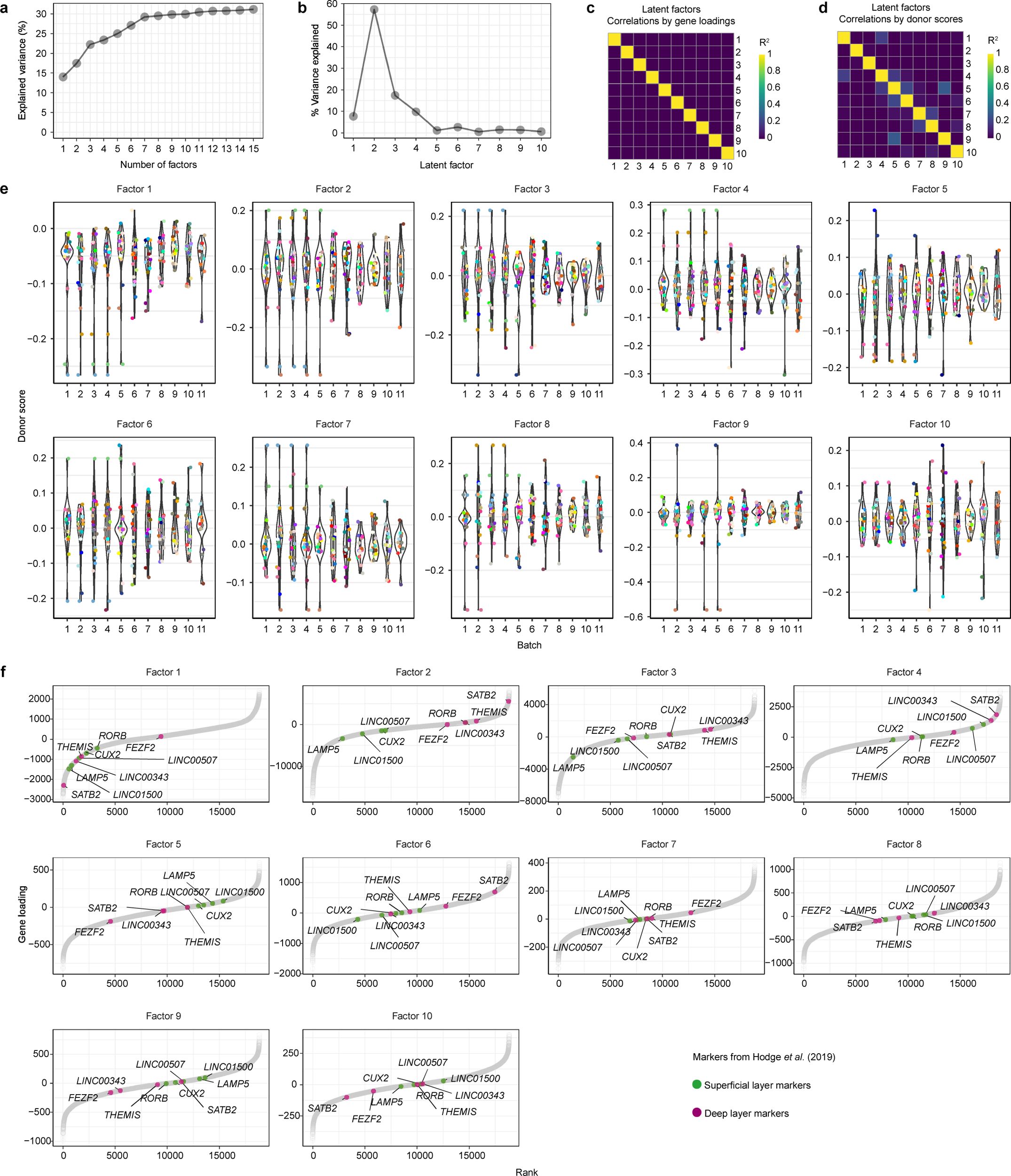
Properties of the latent factors inferred from snRNA-seq data. **a,** Total % variance in expression explained by latent factors with different numbers of requested factors *k*. **b,** Fraction of variance explained by each latent factor in an analysis with 10 requested factors. **c-d,** Independence of latent factors, visualized as Pearson correlation heatmaps of factors’ **(c)** gene loadings (*n* = 125,437 gene/cell-type combinations) and **(d)** donor scores (*n* = 180 donors). **e,** Expression level of each latent factor (panels) in each donor (points), split by batch (*n* = 20 donors per batch). **f,** Relationship of latent factors to markers of superficial and deep cortical layers from ^75^. Markers label dominant classes of glutamatergic neurons (superficial: *LAMP5*, *LINC00507*, *RORB*; deep: *THEMIS*, *FEZF2*) or spatially restricted subtypes (superficial: Exc L2 LAMP5 LTK, marked by *CUX2* and *LINC01500*; deep: Exc L5-6 THEMIS C1QL3, marked by *SATB2* and *LINC00343*). Factor 2 exhibits the most distinct segregation of these superficial and deep layer markers when genes are ranked by their loadings onto each factor. *n* = 18,830 genes expressed in glutamatergic neurons; colored dots are plotted over the dots of genes not among the markers listed above (grey).

**Extended Data Figure 9.**
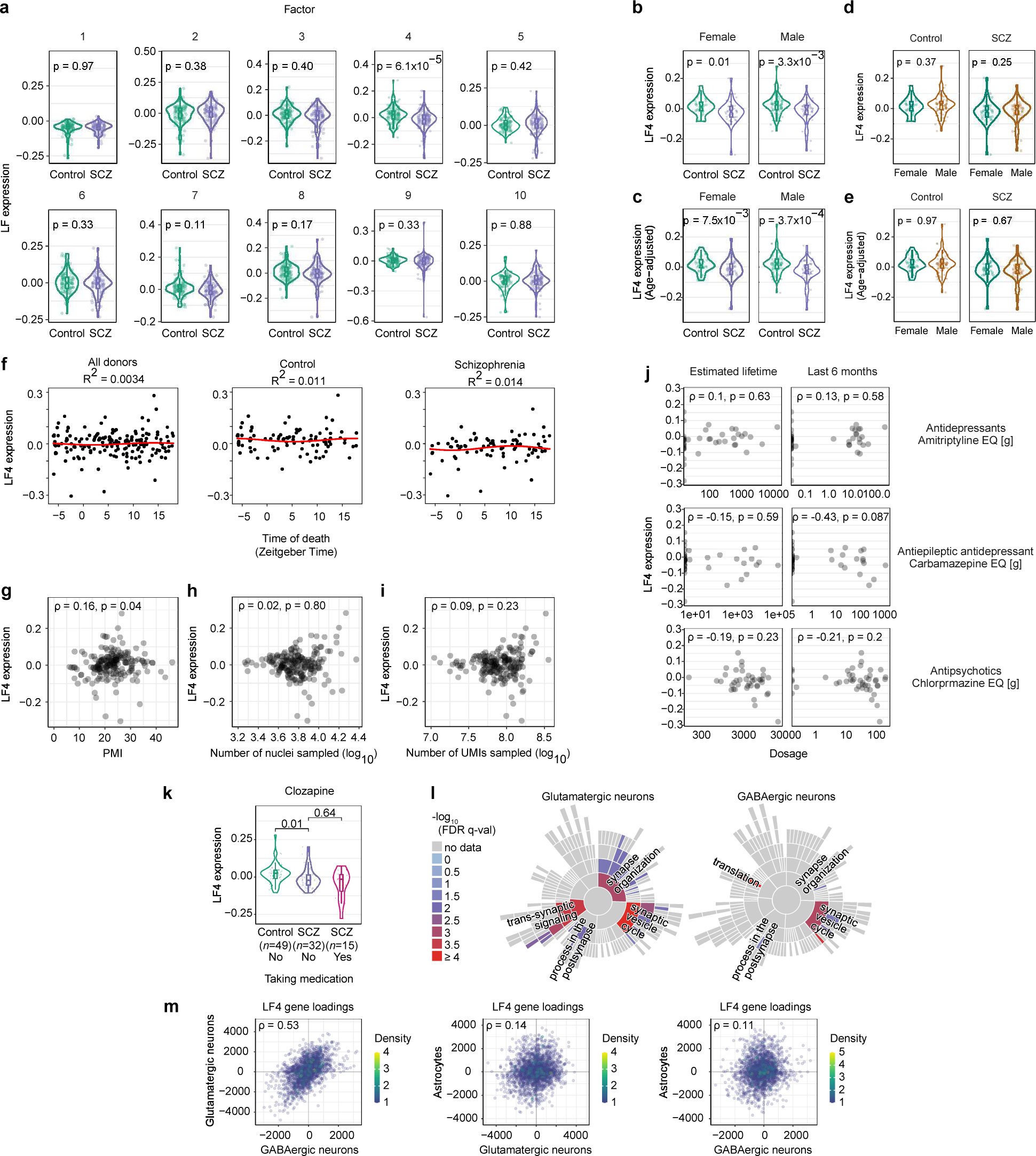
Properties of Latent Factor 4 (LF4). **a,** Expression of each latent factor by case-control status (*n* = 93 controls and 87 cases). P-values are from a two-sided Wilcoxon rank-sum test. Box plots show interquartile ranges; whiskers, 1.5x the interquartile interval; central lines, medians; notches, confidence intervals around medians. **b,** Expression of LF4 by case-control status, split by sex (female: *n* = 31 controls and 39 cases; male: *n* = 62 controls and 48 cases). P-values are from a two-sided Wilcoxon rank-sum test. Box plots show interquartile ranges; whiskers, 1.5x the interquartile interval; central lines, medians; notches, confidence intervals around medians. Note that the more-modest p-value for the females-only analysis relative to the males-only analysis appears to represent the smaller sample (70 females vs. 110 males) rather than a weaker relationship to schizophrenia status; please see also **Extended Data Fig. 18h**. **c,** Similar plots as in **b**, here displaying LF4 expression values adjusted for donor age. **d,** Expression of LF4 by sex, split by case-control status (controls: *n* = 31 females and 62 males; cases: *n* = 39 females and 48 males). P-values are from a two-sided Wilcoxon rank-sum test. Box plots show interquartile ranges; whiskers, 1.5x the interquartile interval; central lines, medians; notches, confidence intervals around medians. **e,** Similar plots as in **d**, here displaying LF4 expression values adjusted for donor age. **f-k,** Relationship of LF4 expression measurements to other available donor and tissue characteristics: **(f)** time of death in zeitgeber time (ZT), with rhythmicity analyses performed as in ^83^; **(g)** post-mortem interval; **(h)** number of nuclei sampled; **(i)** number of UMIs sampled; **(j)** use of psychiatric medications (left column) across each donor’s lifespan or (right column) in the last 6 months prior to death; and **(k)** use of clozapine. Correlation coefficients in **g-j** are Spearman’s ⍴. P-values in **k** are from a two-sided Wilcoxon rank-sum test. Box plots show interquartile ranges; whiskers, 1.5x the interquartile interval; central lines, medians; notches, confidence intervals around medians. **l,** Concentrations of the strongest enriched neuronal gene-expression changes in LF4 among synaptic functions as annotated by SynGO ^91^. Plots show categories of SynGO biological processes. **m,** See also **Fig. 2a**. LF4 involves broadly similar gene-expression effects in glutamatergic and GABAergic neurons, and a distinct set of gene-expression effects in astrocytes. Genes plotted are the protein-coding genes that are expressed (at levels of at least 10 UMIs per 10^5^) in both cell types (Spearman’s ⍴; *n* = 1,538, 1,067, and 1,131 genes respectively).

**Extended Data Figure 10.**
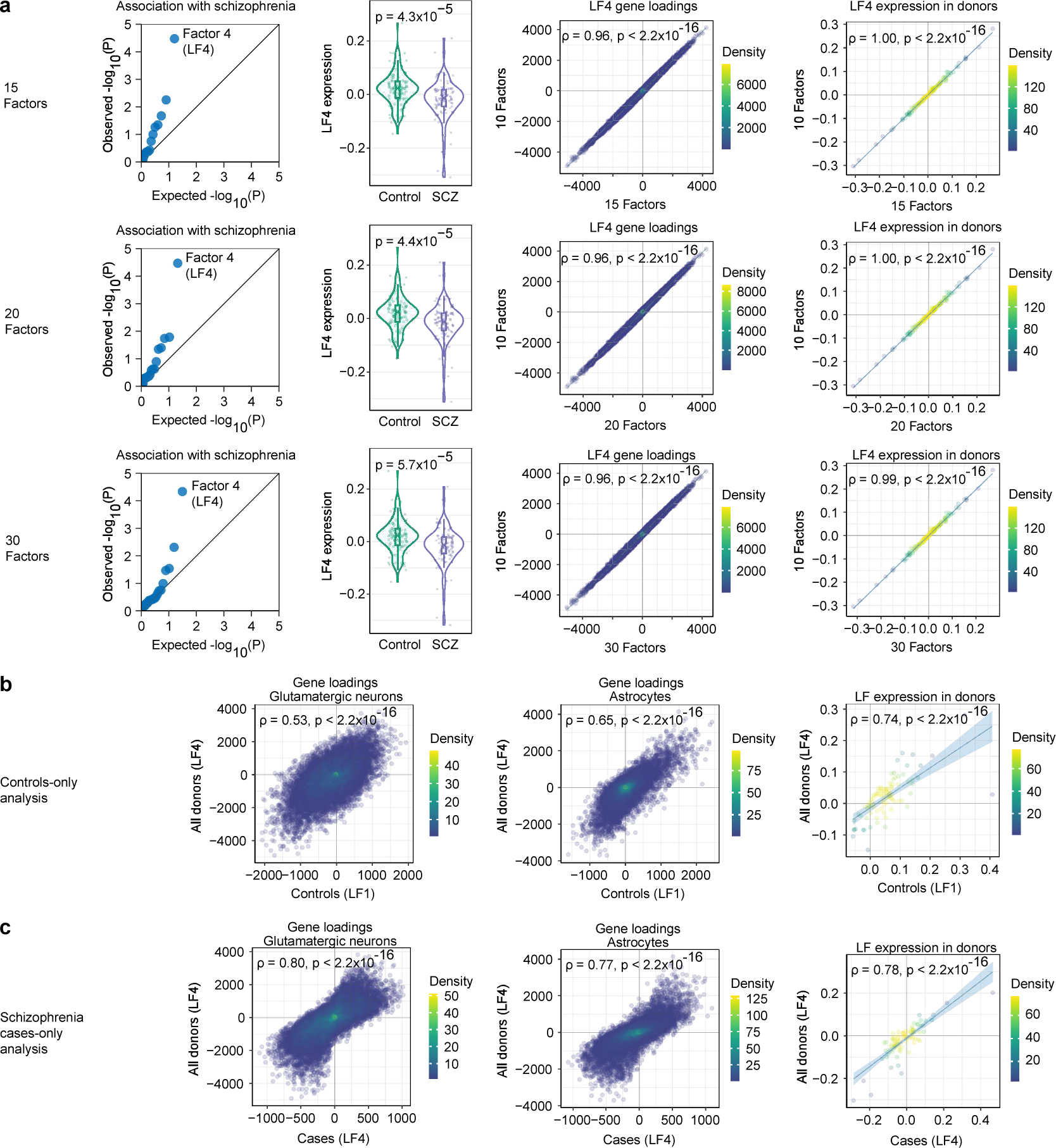
Robustness of Latent Factor 4 (LF4) to analysis parameters. LFs similar to LF4 were identified in **(a)** analyses with different numbers of factors (*n* = 180 donors), **(b)** a controls-only analysis (*n* = 93 donors), and **(c)** a cases-only analysis (*n* = 87 donors). **a,** Column 1: Association of latent-factor expression levels with schizophrenia case-control status, shown as a quantile-quantile plot that compares observed -log_10_ p-values to the distribution of -log_10_ p-values expected under a null hypothesis (*n* = 15, 20, and 30 factors). The observed p-values were calculated for each latent factor by a two-sided Wilcoxon rank-sum test of latent factor expression levels (by donor) between cases and controls. In all analyses, LF4 is the factor that deviates the most from the line of unity and displays the strongest association with schizophrenia case-control status. Column 2: Expression of LF4 by case-control status (*n* = 93 controls and 87 cases). P-values are from a two-sided Wilcoxon rank-sum test. Box plots show interquartile ranges; whiskers, 1.5x the interquartile interval; central lines, medians; notches, confidence intervals around medians. Shaded regions represent 95% confidence intervals. Column 3: Comparison of gene loadings (*n* = 125,437 gene/cell-type combinations) that demonstrates the relationship of LF4 inferred from an analysis requesting 10 factors to LF4 inferred from an analysis requesting 15, 20, or 30 factors (Spearman’s ⍴). Shaded regions around regression lines represent 95% confidence intervals. Column 4: Comparison of donor expression levels (*n* = 180 donors) that demonstrates the relationship of LF4 inferred from an analysis requesting 10 factors to LF4 inferred from an analysis requesting 15, 20, or 30 factors (Spearman’s ⍴). Shaded regions around regression lines represent 95% confidence intervals. **b,** Column 1: Comparison of gene loadings from glutamatergic neurons (*n* = 18,829 genes) that demonstrates the relationship of LF4 inferred from an analysis of all donors to LF1 inferred from an analysis of only control donors (Spearman’s ⍴). Shaded regions around regression lines represent 95% confidence intervals. Column 2: Similar plot as in Column 1, here plotting gene loadings from astrocytes (*n* = 18,346 genes). Column 3: Comparison of donor expression levels (*n* = 180 donors) that demonstrates the relationship of LF4 inferred from an analysis of all donors to LF1 inferred from an analysis of only control donors (Spearman’s ⍴). Shaded regions around regression lines represent 95% confidence intervals. **c,** Similar plots as in **b**, here for the relationship of LF4 inferred from an analysis of all donors to LF4 inferred from an analysis of only donors with schizophrenia.

**Extended Data Figure 11.**
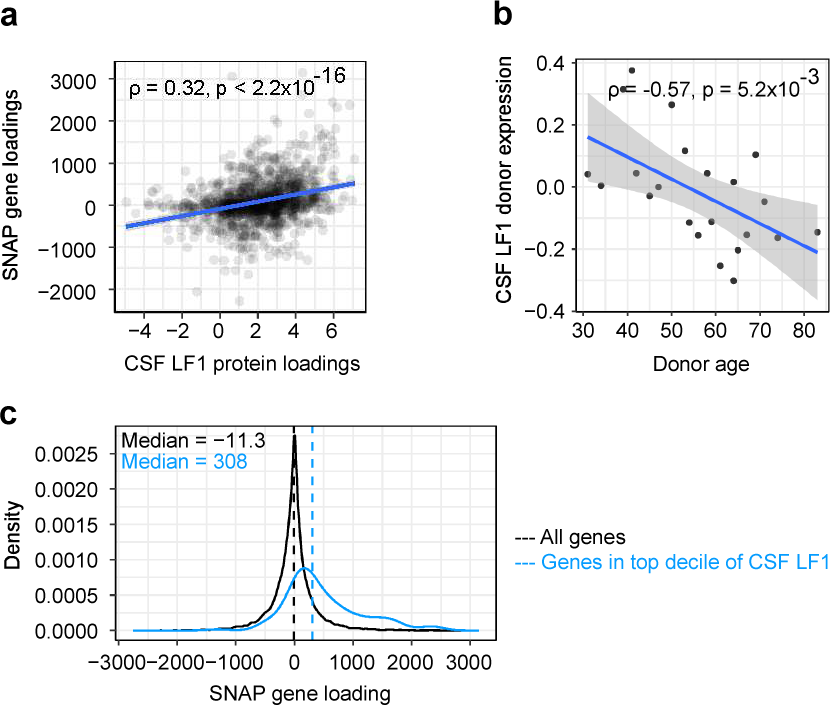
Latent factor analysis of cerebrospinal fluid (CSF) proteomics data from different individuals identifies a factor resembling SNAP. To assess the biological significance of SNAP, we also sought evidence that SNAP manifests in the proteins that can be sampled from cerebrospinal fluid (CSF). We analyzed available data from a mass-spectrometry proteomics analysis of cerebrospinal fluid (CSF) from 22 healthy human donors ^82^, performing a latent factor analysis that is conceptually analogous to our analysis (in Fig. 1f) of cell-type-specific RNA-expression measurements in the brain donors (but of an independent data set, derived from a distinct set of donors). The top latent factor in analysis of the CSF proteomics data (explaining >15% of inter-individual variation in CSF protein measurements) bore a strong resemblance to SNAP. **a,** Relationship of SNAP gene loadings to the top latent factor in an analysis of inter-individual variation in CSF protein levels (CSF LF1) using quantitative protein abundance measurements from ^82^ (Spearman’s ⍴; *n* = 1,341 genes/proteins shared between both analyses). For SNAP, each gene is represented by a single composite loading representing gene loadings from all cell types (weighted by its median expression in each cell type). Shaded region represents 95% confidence interval. **b,** Relationship of CSF LF1 donor scores to age (Spearman’s ⍴; *n* = 22 donors). Shaded region represents 95% confidence interval. **c,** Density plot showing distribution of SNAP gene loadings for (black) all genes and genes encoding proteins that are strongly recruited (top decile) by (blue) CSF LF1. Distributions were found to be different by Wilcox test (p = 2.1×10^−28^, two-sided Wilcoxon rank-sum test).

**Extended Data Figure 12.**
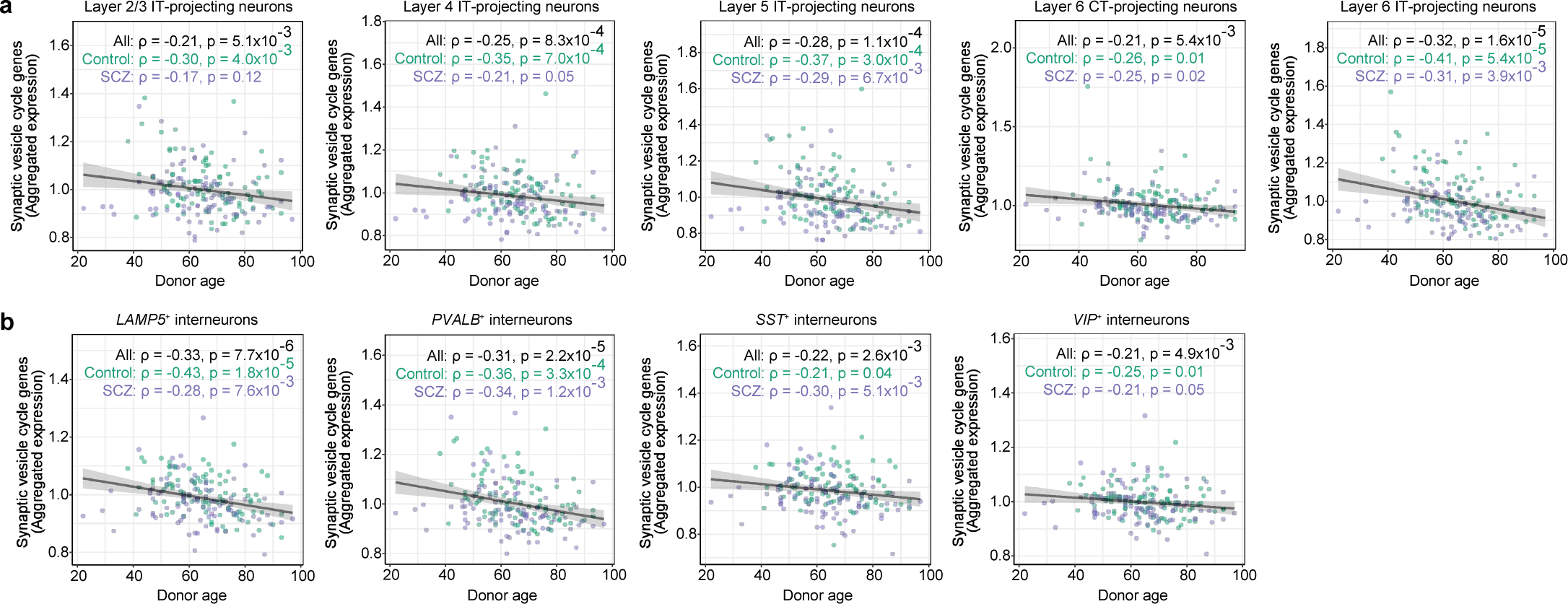
Relationship of synaptic vesicle cycle gene expression in neuronal subtypes to advancing age. **a-b,** See also **Fig. 2c**. Neuronal expression of synaptic vesicle cycle genes in the most abundant subtypes of **(a)** glutamatergic and **(b)** GABAergic neurons (across 180 donors), plotted against donor age (Spearman’s ⍴). Expression values are the fraction of all UMIs in each donor (from the indicated subtype) that are derived from these genes, normalized to the median expression among control donors. Shaded regions represent 95% confidence intervals. The observed decline in schizophrenia and aging was consistent with earlier observations that expression of genes for synaptic components is reduced in schizophrenia ^168^ and with advancing age ^169^.

**Extended Data Figure 13.**
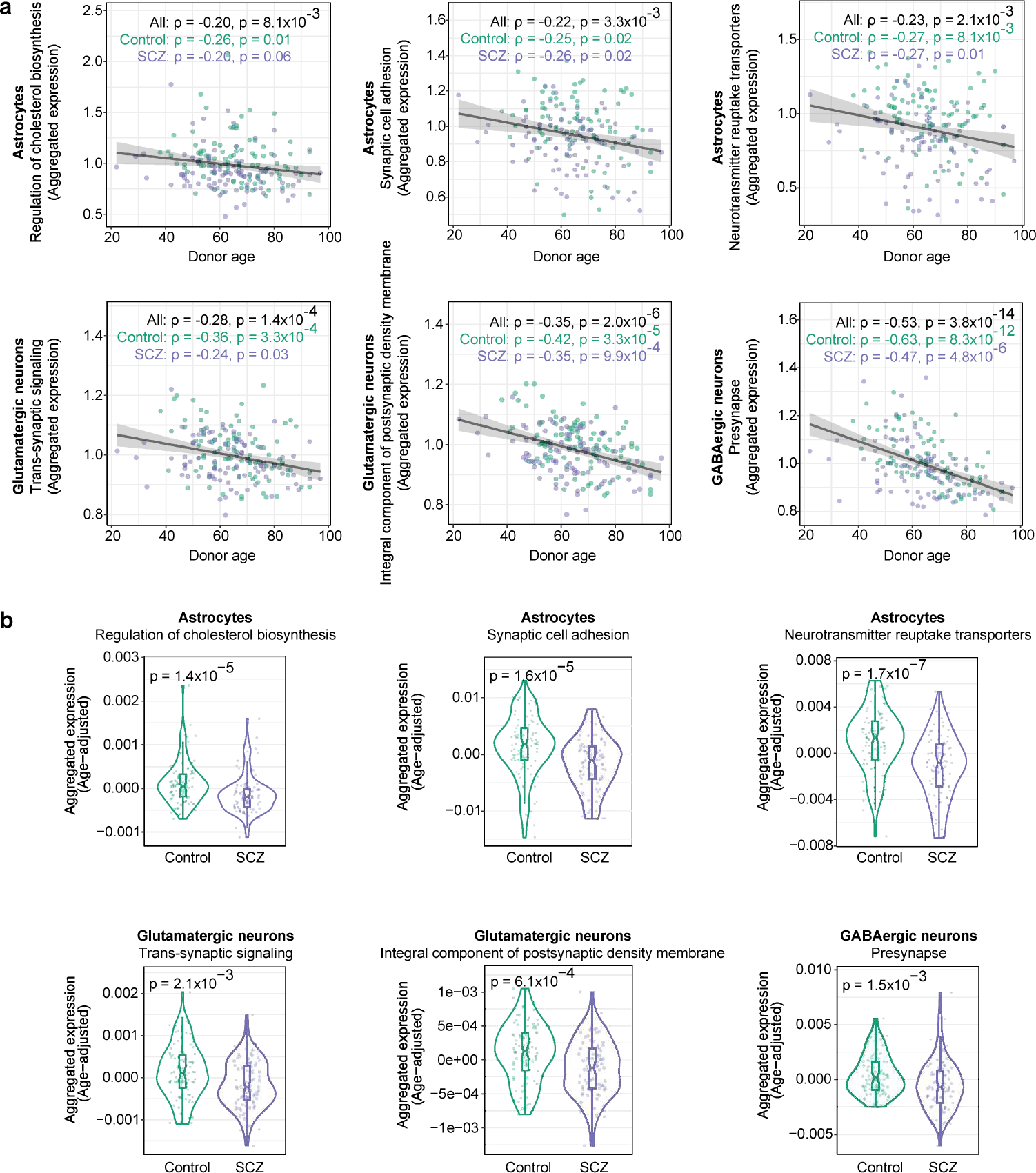
Relationship of gene-set expression in astrocytes and neurons to advancing age and schizophrenia. **a,** Expression of gene sets enriched in the astrocyte and neuronal components of LF4 (across 180 donors), plotted against donor age (Spearman’s ⍴). Expression values are the fraction of all UMIs in each donor (from the indicated cell type) that are derived from these genes, normalized to the median expression among control donors. Shaded regions represent 95% confidence intervals. **b,** Expression (by donor, separated by schizophrenia case-control status; *n* = 180 donors) of gene sets enriched in the astrocyte and neuronal components of LF4. Expression values are the fraction of all UMIs in each donor (from the indicated cell type) that are derived from these genes, adjusted for donor age. P-values from a two-sided Wilcoxon rank-sum test comparing the affected to the unaffected donors are reported at the top of each panel. Box plots show interquartile ranges; whiskers, 1.5x the interquartile interval; central lines, medians; notches, confidence intervals around medians.

**Extended Data Figure 14.**
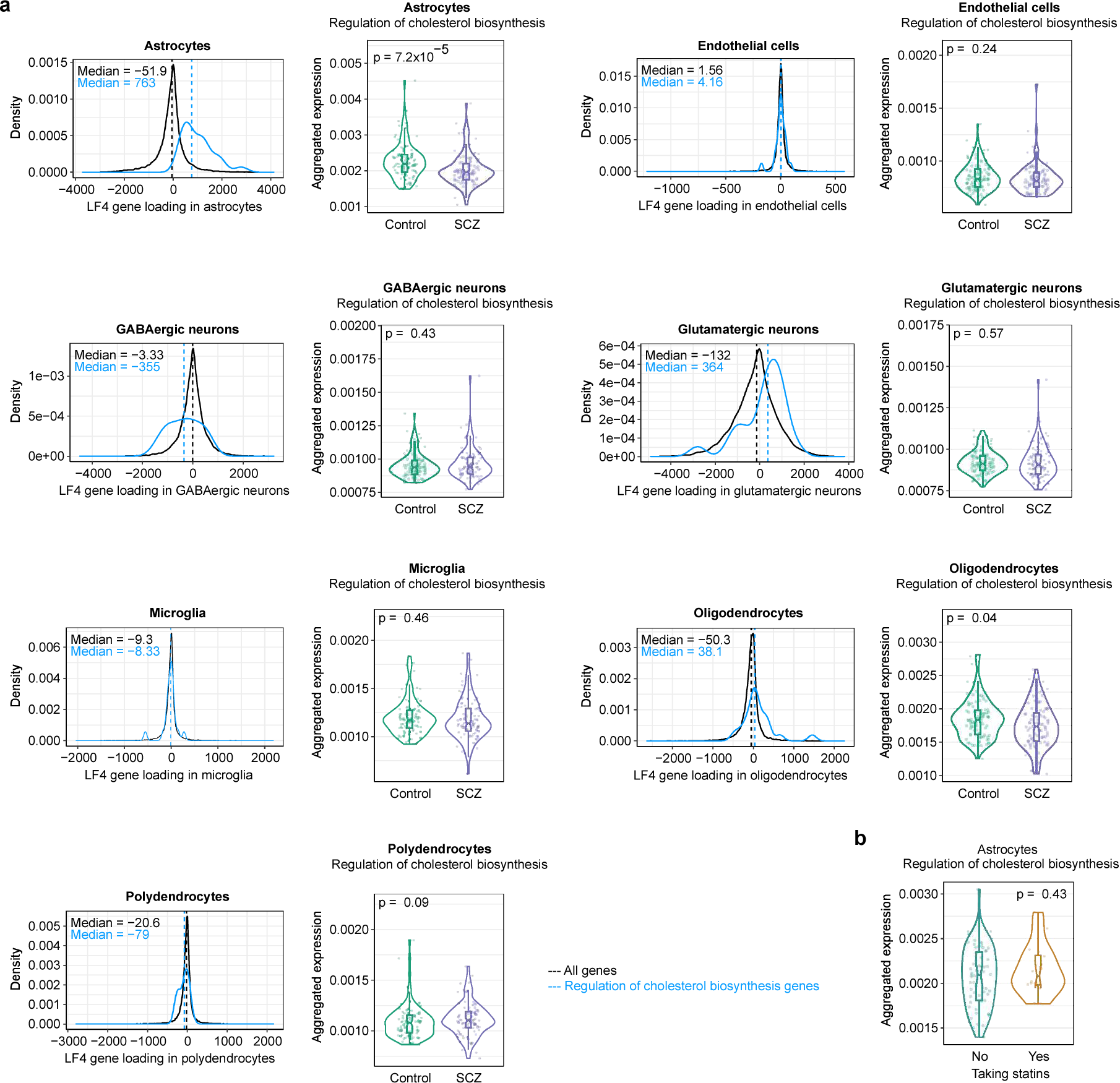
Expression of cholesterol-biosynthesis genes in cortical cell types. **a,** See also **Fig. 2d**. For each cortical cell type: (Left) Distributions of LF4 gene loadings for (black) all expressed genes and (blue) specifically for genes annotated by GO as having roles in cholesterol biosynthesis (core genes contributing to the enrichment of GO:0045540 (“cholesterol biosynthesis genes”) in that cell type’s component of LF4. (Right) Each cell type’s expression of cholesterol biosynthesis genes (by donor, split by schizophrenia case-control status; *n* = 180 donors). Expression values are the fraction of all UMIs in each donor (from the indicated cell type) that are derived from these genes. P-values are from a two-sided Wilcoxon rank-sum test comparing the affected to the unaffected donors. Box plots show interquartile ranges; whiskers, 1.5x the interquartile interval; central lines, medians; notches, confidence intervals around medians. **b,** Expression in astrocytes of cholesterol biosynthesis genes by donor, separated by statin intake among donors with available medication data (*n* = 63 donors not taking statins and 16 donors taking statins). Expression values are the fraction of all UMIs in each donor’s astrocytes that are derived from these genes. P-value is from a two-sided Wilcoxon rank-sum test. Box plots show interquartile ranges; whiskers, 1.5x the interquartile interval; central lines, medians; notches, confidence intervals around medians.

**Extended Data Figure 15.**
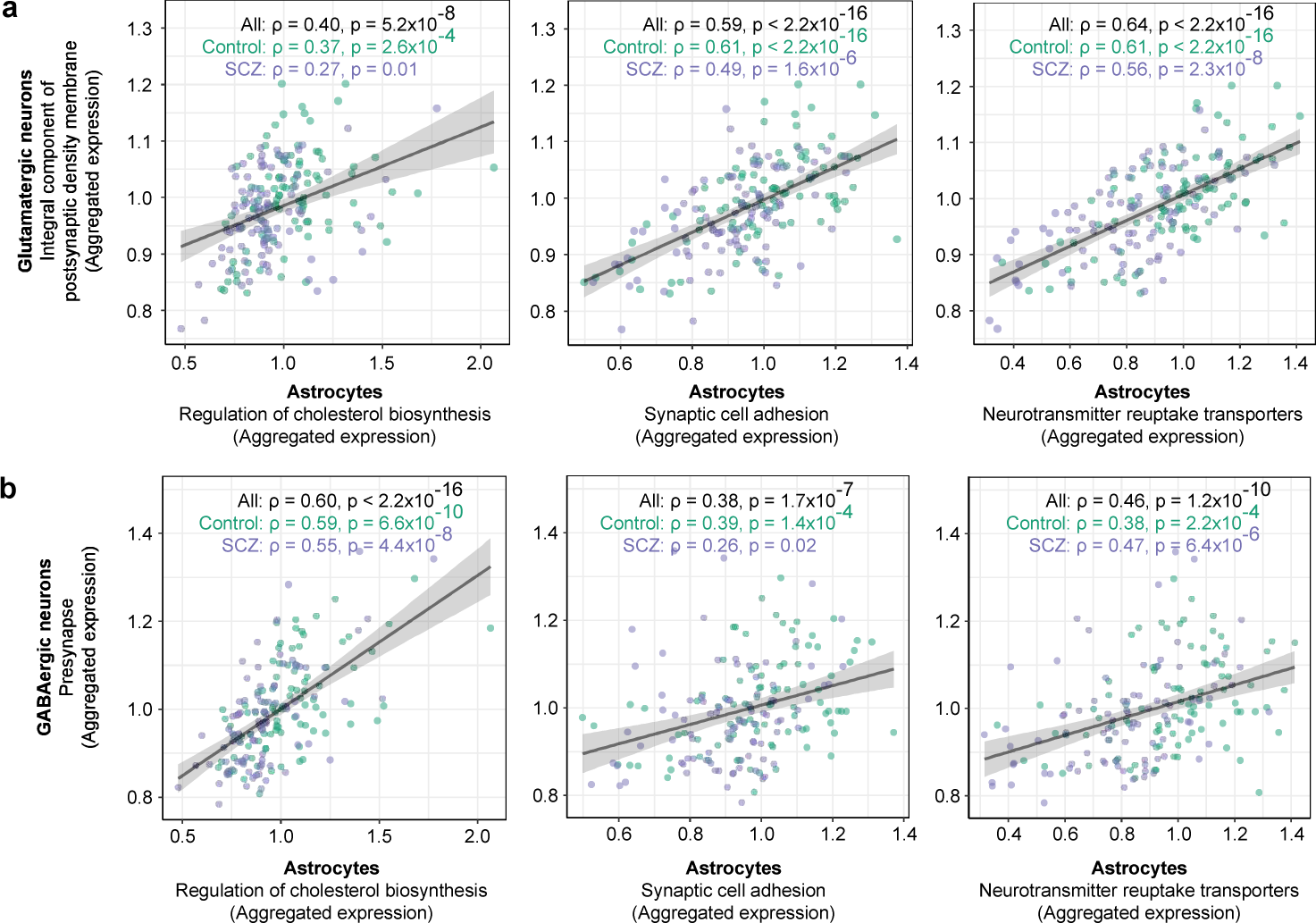
Concerted synaptic investments by neurons and astrocytes. See also Fig. 2e. **a,** Relationship of donors’ glutamatergic-neuron expression of genes that are integral components of the postsynaptic density membrane (core genes contributing to the enrichment of GO:0099061) to astrocyte expression of (top) cholesterol biosynthesis, (middle) synaptic adhesion, and (bottom) neurotransmitter reuptake transporters (Spearman’s ⍴). Expression values are the fraction of all UMIs in each donor (from the indicated cell type) that are derived from these genes, normalized to the median expression among control donors. Shaded regions represent 95% confidence intervals. **b,** Similar plots as in **a**, here for donors’ GABAergic-neuron expression of presynapse genes (core genes contributing to the enrichment of GO:0098793) on the x-axis.

**Extended Data Figure 16.**
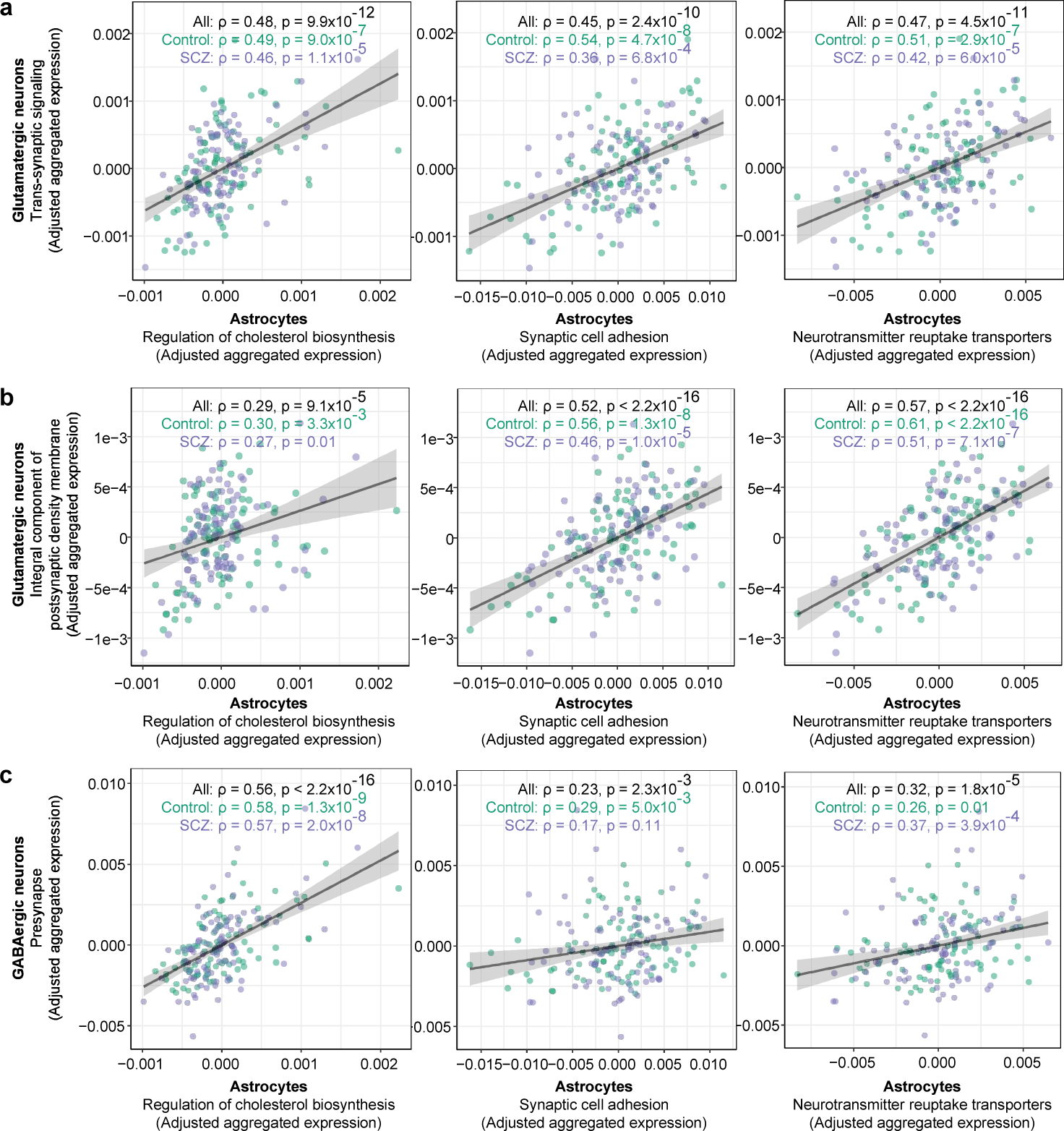
Concerted synaptic investments by neurons and astrocytes, adjusted for age and schizophrenia case-control status. **a-c** See also **Fig. 2e**. Relationship of donors’ neuronal gene expression to astrocyte gene expression (Spearman’s ⍴), adjusted for age and case-control status. Astrocyte gene sets plotted on the x-axes represent (left) cholesterol biosynthesis, (middle) synaptic adhesion, and (right) neurotransmitter reuptake transporters. Neuronal gene sets plotted on the y-axes represent **(a)** trans-synaptic signaling in glutamatergic neurons, **(b)** integral component of postsynaptic density, and **(c)** presynapse genes. Expression values are the fraction of all UMIs in each donor (from the indicated cell type) that are derived from these genes, adjusted for donor age and schizophrenia case-control status. Shaded regions represent 95% confidence intervals.

**Extended Data Figure 17.**
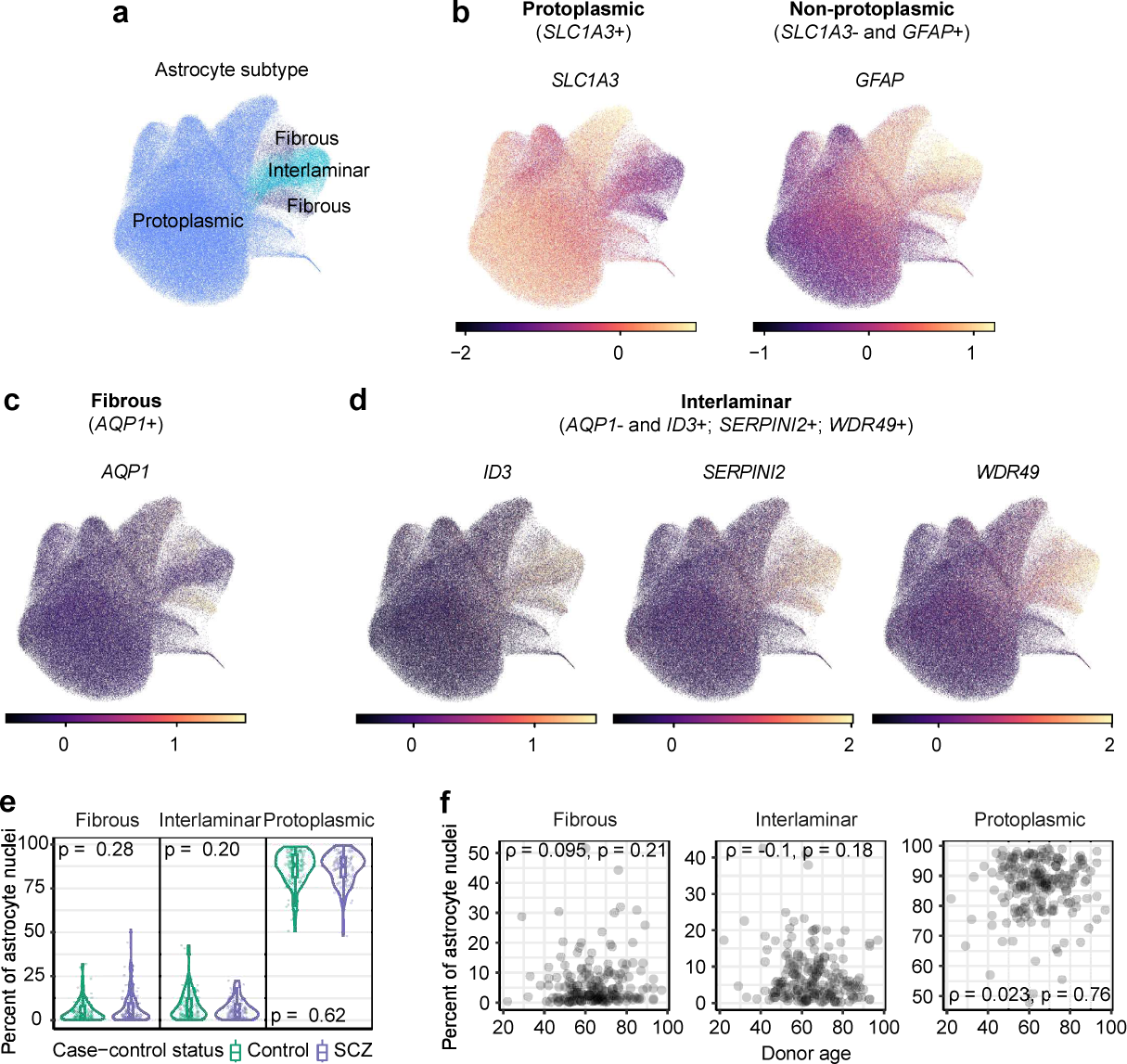
Astrocyte subtype classification and proportions across donors. **a,** Two-dimensional projection of the RNA-expression profiles of 179,764 astrocyte nuclei from 180 donors, reproduced from **Fig. 3a**. Nuclei are colored by their assignments to subtypes of astrocytes using classifications from ^75^ and ^76^. The same projection is used in panels B to D below. **b-d,** Expression levels of marker genes for subtypes of **(b)** protoplasmic astrocytes (*SLC1A3*+) and non-protoplasmic astrocytes (*SLC1A3*- and *GFAP*+) comprising the **(c)** fibrous (*AQP1*+) and **(d)** interlaminar (*AQP1*- and *ID3*+, *SERPINI2*+, and *WDR49*+) subtypes. Markers are from^75^ or from transcriptomically similar subtypes in ^76^. Values represent Pearson residuals from variance stabilizing transformation (VST). **e,** Proportions of astrocyte subtypes in BA46 by schizophrenia status (*n* = 93 unaffected and 87 affected). P-values from a two-sided Wilcoxon rank-sum test comparing the affected to the unaffected donors are reported at the top of each panel. Box plots show interquartile ranges; whiskers, 1.5x the interquartile interval; central lines, medians; notches, confidence intervals around medians. **f,** Relationship of sampled astrocyte subtype proportions to donor age (Spearman’s ⍴).

**Extended Data Figure 18.**
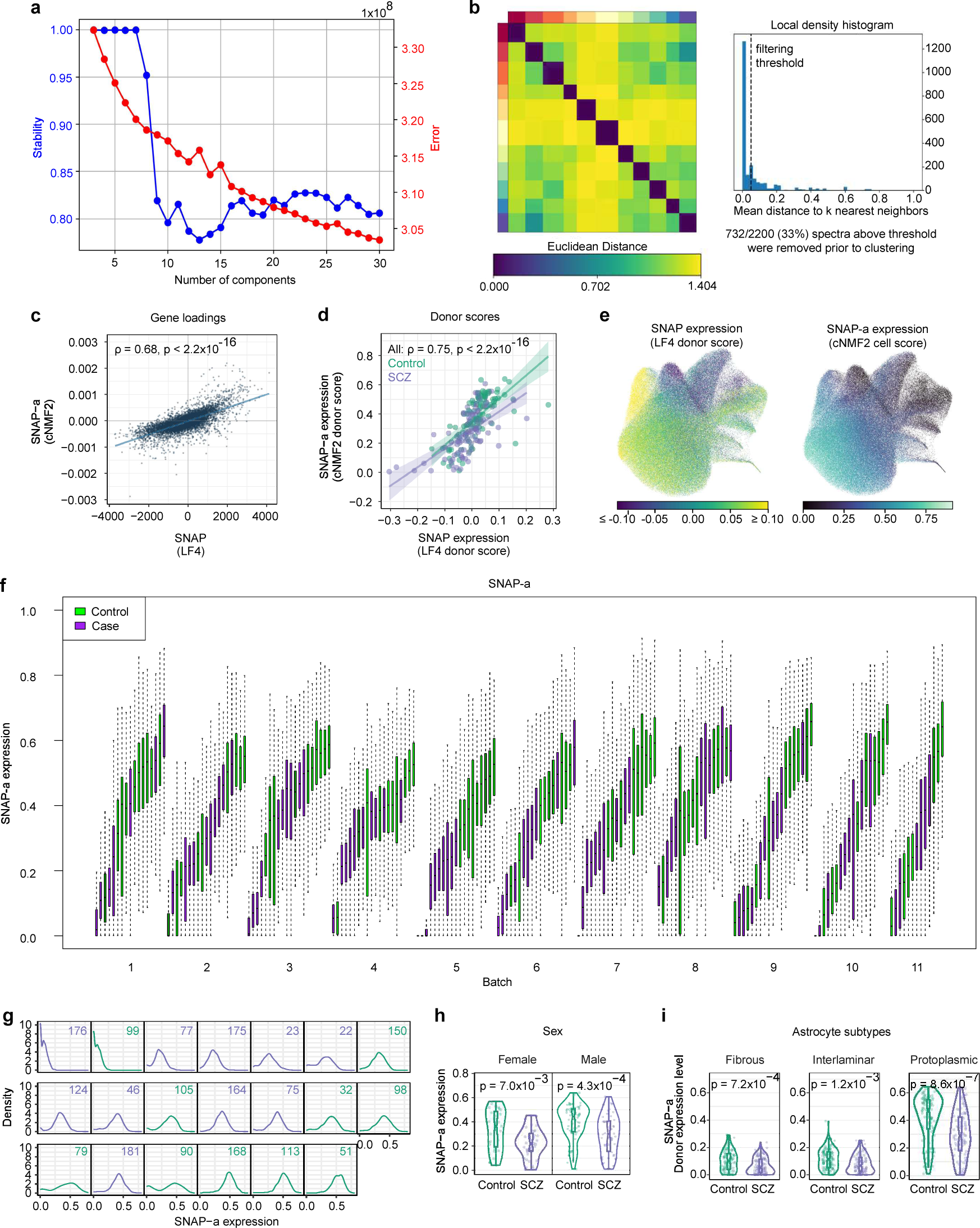
Astrocyte gene-expression programs inferred by cNMF (SNAP-a) and their relationship to SNAP. **a,** Visualization of the trade-off between error and stability of cNMF factors as a function of the number of factors *k*. 11 factors were requested based on these results. **b,** Clustergram of consensus matrix factorization estimates. Each color on the x- and y-axes represents one of 11 cNMF factors. **c-d,** Relationship of SNAP-a to SNAP by **(c)** gene loadings (*n* = 33,611 genes) and **(d)** donors’ expression levels of each factor (*n* = 180 donors) (Spearman’s ⍴). Shaded regions represent 95% confidence intervals. **e,** UMAP of RNA-expression patterns from 179,764 astrocyte nuclei from 180 donors, using the same projection from **Fig. 3a-c**. Nuclei are colored by (left) each donor’s expression of SNAP or (right) each cell’s expression of the astrocyte component of SNAP (cNMF2, also referred to as SNAP-a). SNAP-a is reproduced from **Fig. 3c** for comparison with SNAP. **f,** Distributions of SNAP-a expression levels among astrocytes in each donor, split by experimental batch. Box plots show interquartile ranges; whiskers, 1.5x the interquartile interval; central lines, medians. **g,** Density plots showing distributions of SNAP-a expression levels among astrocytes in each donor for one representative batch (batch 4) out of 11 batches. Labels in top-right corners indicate anonymized research IDs at the Harvard Brain Tissue Resource Center. Colors represent case-control status (green: controls; purple: schizophrenia cases). At the single-astrocyte level, SNAP-a expression exhibited continuous, quantitative variation rather than discrete state shifts by a subpopulation of astrocytes, supporting the idea that astrocyte biological variation extends beyond polarized states ^17,170,171^. **h,** Distributions of SNAP-a expression levels by case-control status, split by sex. P-values from a two-sided Wilcoxon rank-sum test comparing the affected to the unaffected donors are reported at the top of each panel. Box plots show interquartile ranges; whiskers, 1.5x the interquartile interval; central lines, medians; notches, confidence intervals around medians. **i,** Distributions of SNAP-a expression levels by case-control status, split by astrocyte subtype. P-values from a two-sided Wilcoxon rank-sum test comparing the affected to the unaffected donors are reported at the top of each panel. Box plots show interquartile ranges; whiskers, 1.5x the interquartile interval; central lines, medians; notches, confidence intervals around medians.

**Extended Data Figure 19.**
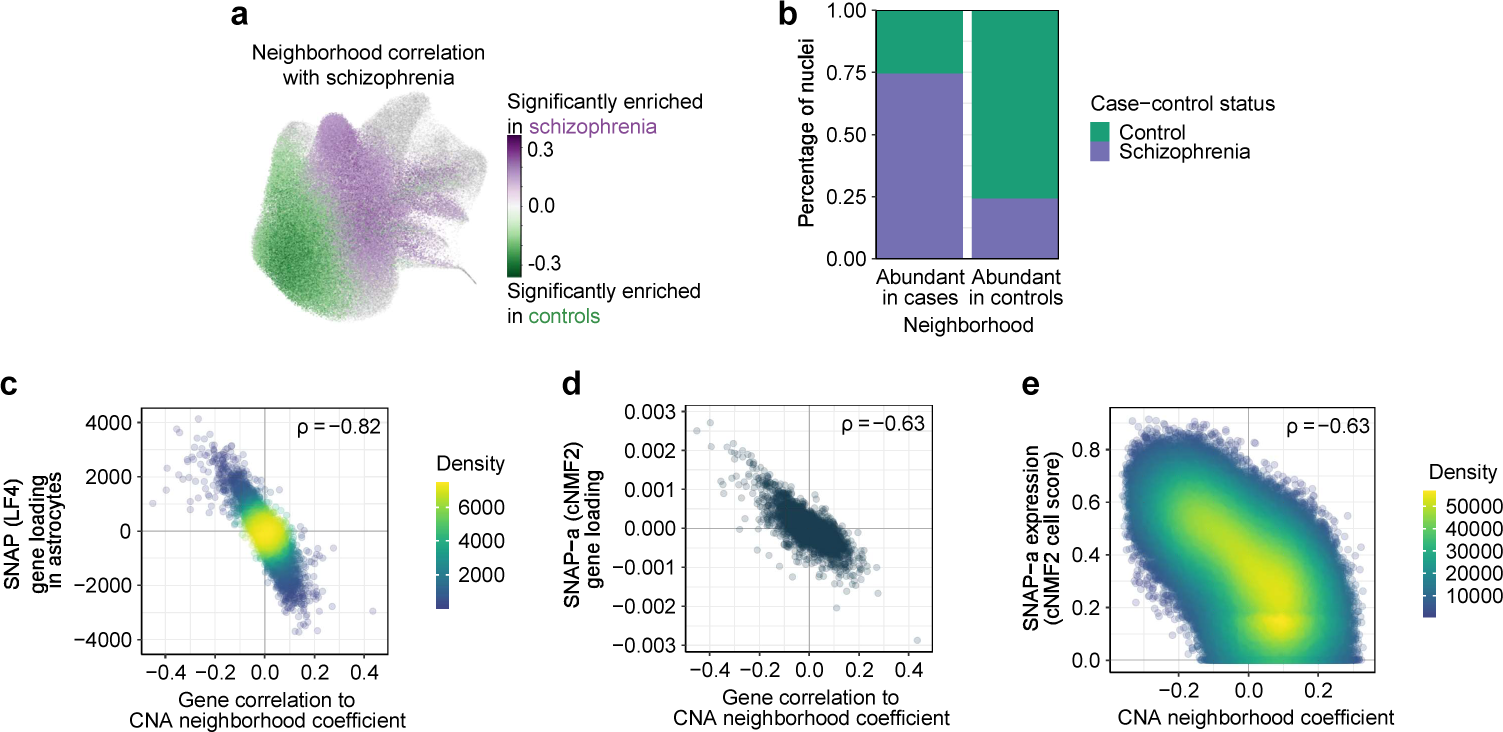
Identification of astrocyte transcriptional neighborhoods associated with schizophrenia case-control status by co-varying neighborhood analysis. To further assess the robustness of the astrocyte gene-expression changes represented by SNAP and SNAP-a, we employed a third computational approach, co-varying neighborhood analysis (CNA) ^96^. **a,** Same projection as in **Fig. 3a-c**, but with points colored according to their transcriptional neighborhood’s correlation to schizophrenia case-control status (*n* = 179,764 astrocyte nuclei from 180 donors). Among cells whose neighborhood coefficients passed an FDR < 0.05 threshold for association, purple indicates high correlation to case status and green indicates high correlation to control status. All other cells with FDR > 0.05 for association are colored in gray. **b,** Proportion of nuclei in each of the indicated astrocyte transcriptional neighborhoods that are assigned to schizophrenia cases and controls (*n* = 34,271 nuclei abundant in cases and 38,327 nuclei abundant in controls). **c-d,** Relationship of genes’ correlation to schizophrenia-associated transcriptional neighborhoods to **(c)** the astrocyte component of SNAP (*n* = 8,997 shared genes) and **(d)** SNAP-a by their gene loadings (*n* = 9,015 shared genes) (Spearman’s ⍴). Genes plotted are the subsets of protein-coding genes (with expression levels of at least 1 UMI per 10^5^) that are shared between the indicated pairs of analyses. **e,** Relationship of cell-level neighborhood coefficients for schizophrenia-associated transcriptional neighborhoods to SNAP-a cell scores (Spearman’s ⍴; *n* = 179,764 astrocytes).

**Extended Data Figure 20.**
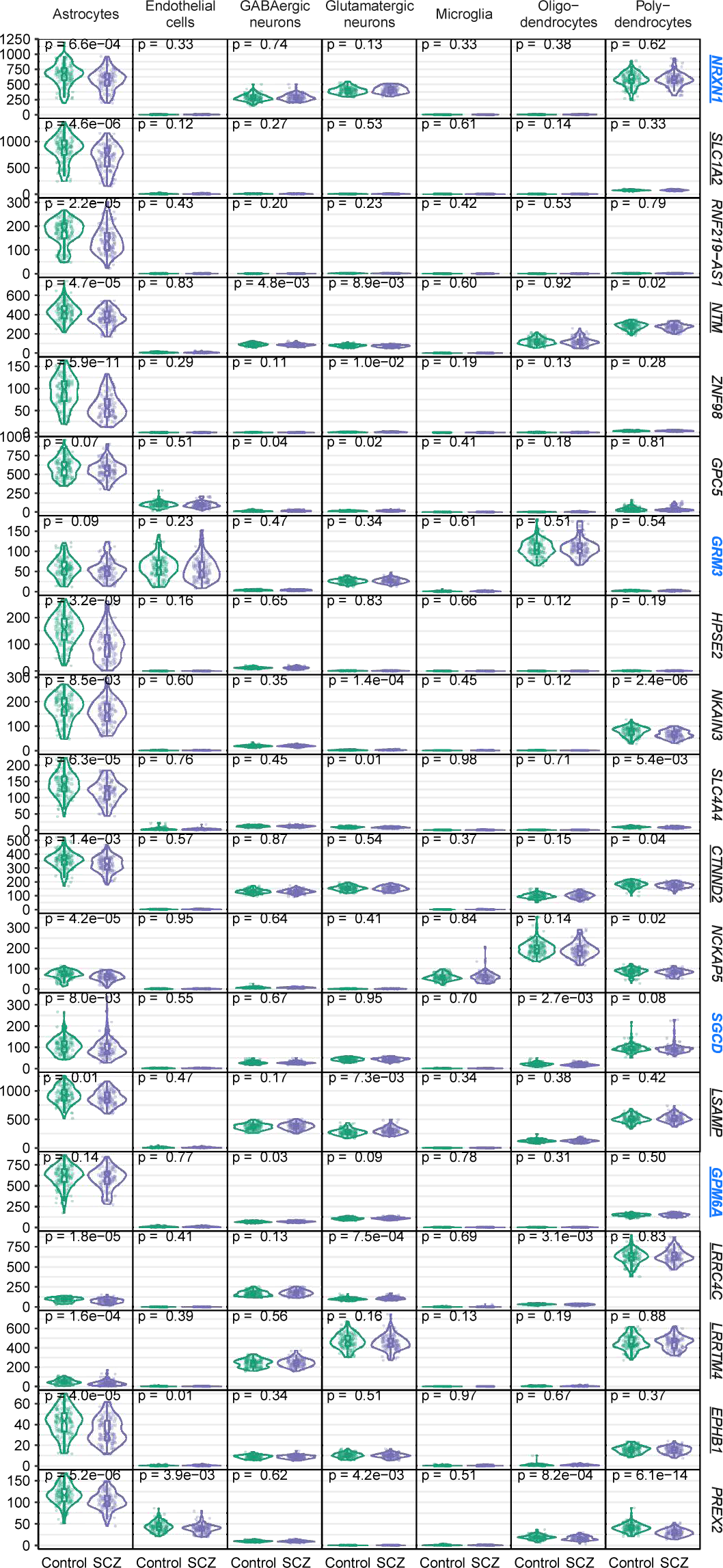
Expression across cell types of genes most strongly recruited by SNAP-a. Expression in each cell type (by donor, separated by schizophrenia status), of the 20 genes that are most strongly recruited by SNAP-a (*n* = 93 unaffected (green) and 87 affected (purple) donors). These included eight genes with roles in adhesion of cells to synapses (*NRXN1, NTM, CTNND2, LSAMP, GPM6A, LRRC4C, LRRTM4,* and *EPHB1*) (as established by earlier work including ^172–181^ and reviewed in ^11,12^). P-values from a two-sided Wilcoxon rank-sum test comparing the affected to the unaffected donors are reported at the top of each panel. Box plots show interquartile ranges; whiskers, 1.5x the interquartile interval; central lines, medians; notches, confidence intervals around medians. Genes that have been strongly implicated in human genetic studies of schizophrenia are highlighted in blue. Genes with known functions in synaptic adhesion (listed above) or neurotransmitter uptake (*SLC1A2*) are underlined.

**Extended Data Figure 21.**
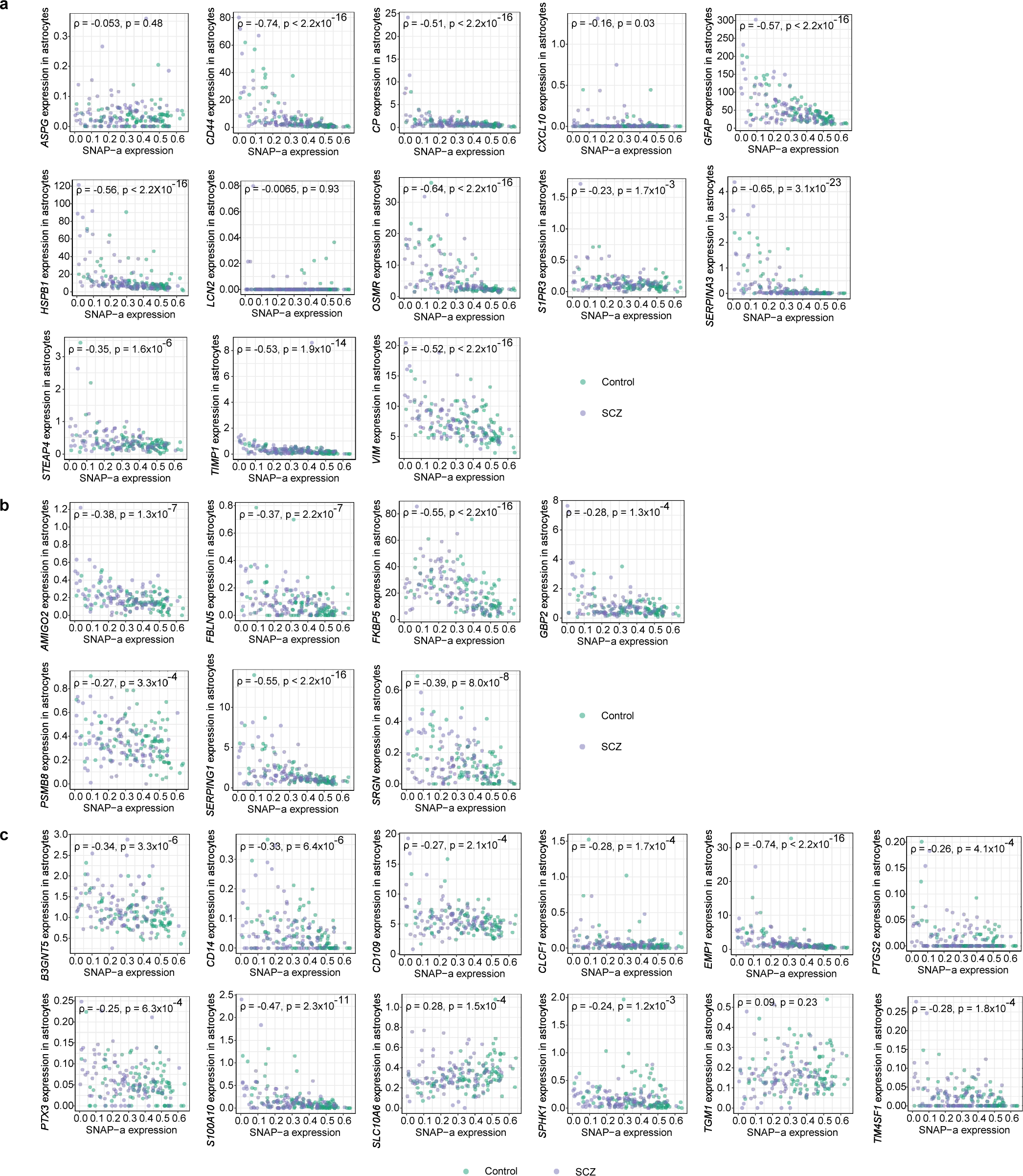
Relationship of reactive astrocyte marker expression to SNAP-a expression. Relationship of donors’ expression levels of reactive astrocyte marker genes to SNAP-a expression (Spearman’s ⍴). Markers are from ^16^ and represent **(a)** pan-reactive (PAN), **(b)** A1, and **(c)** A2 reactive astrocytes.

**Extended Data Figure 22.**
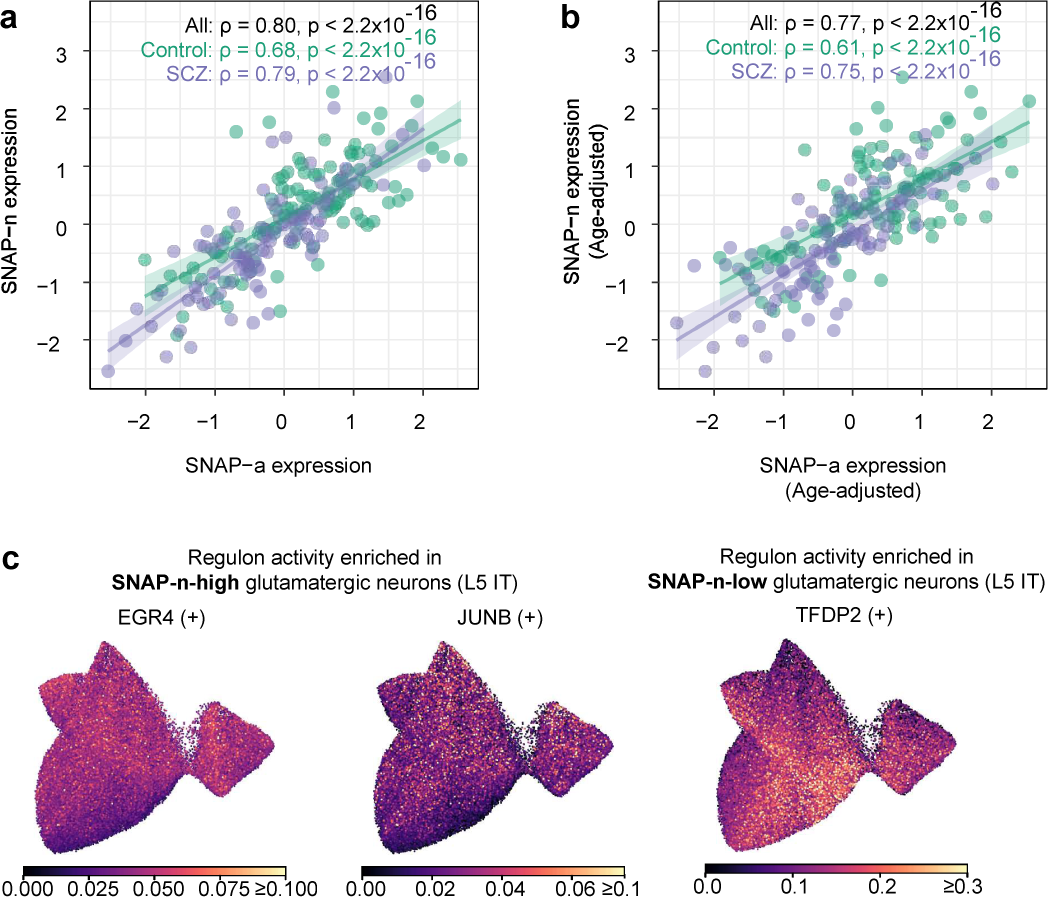
Biological states and transcriptional programs of L5 IT glutamatergic neurons in schizophrenia. **a-b,** Relationship of SNAP-a to SNAP-n (Spearman’s ⍴). Values plotted are **(a)** quantile-normalized and **(b)** donor age-adjusted, quantile-normalized donor scores for each factor. Shaded regions represent 95% confidence intervals. **c,** UMAP of regulon activity scores (as inferred by pySCENIC ^98^) from L5 IT glutamatergic neuron nuclei from 180 donors, using the same projection from **Fig. 3f-h**. Regulons plotted are the most strongly enriched in L5 IT glutamatergic neurons with high versus low SNAP-n expression. (+) indicates that the targets of the indicated regulon were found to be upregulated in expression.

**Extended Data Figure 23.**
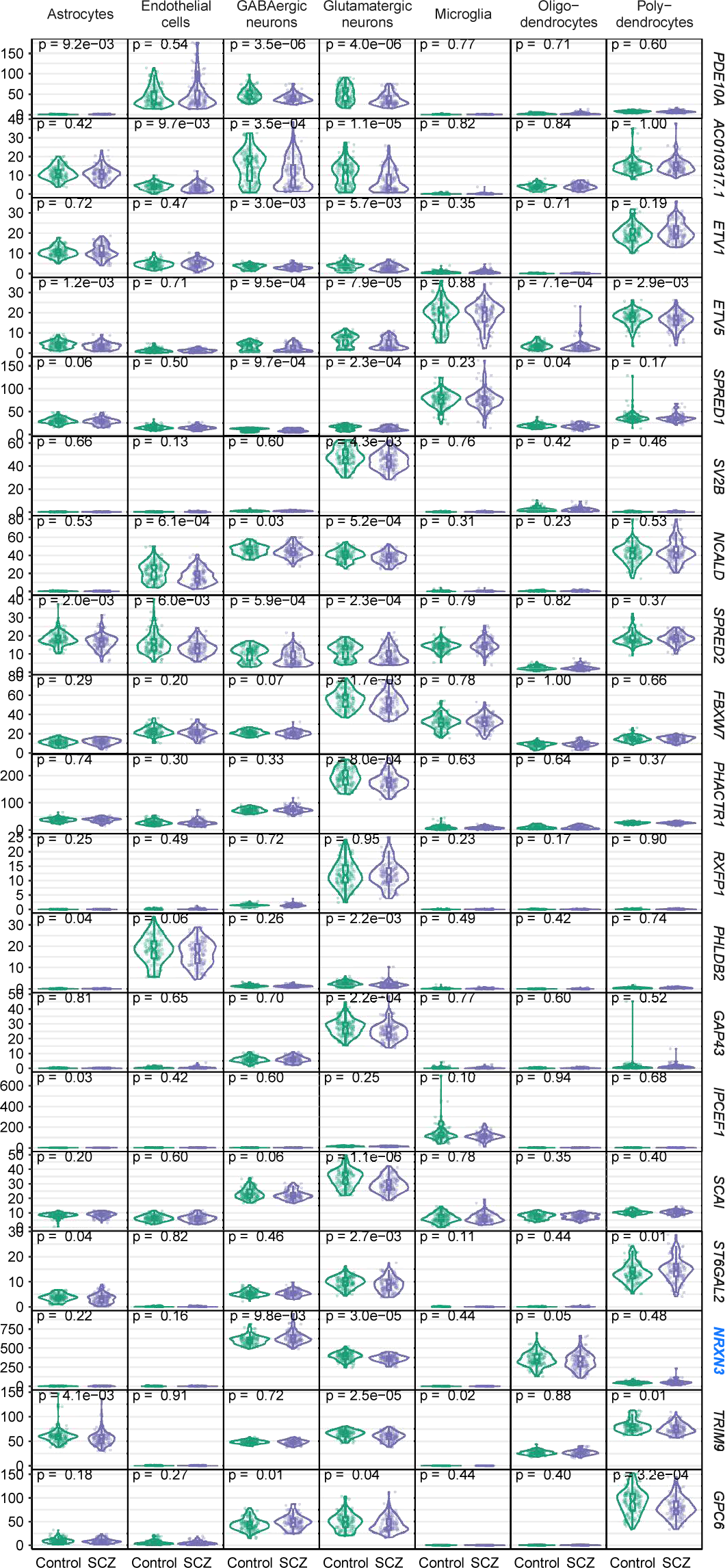
Expression across cell types of genes most strongly recruited by SNAP-n. Expression in each cell type (by donor, separated by schizophrenia case-control status) of the 20 genes that are most strongly recruited by SNAP-n (*n* = 93 controls and 87 cases). P-values from a two-sided Wilcoxon rank-sum test comparing the affected to the unaffected donors are reported at the top of each panel. Box plots show interquartile ranges; whiskers, 1.5x the interquartile interval; central lines, medians; notches, confidence intervals around medians. Genes that have been strongly implicated in human genetic studies of schizophrenia are highlighted in blue.

**Extended Data Figure 24.**
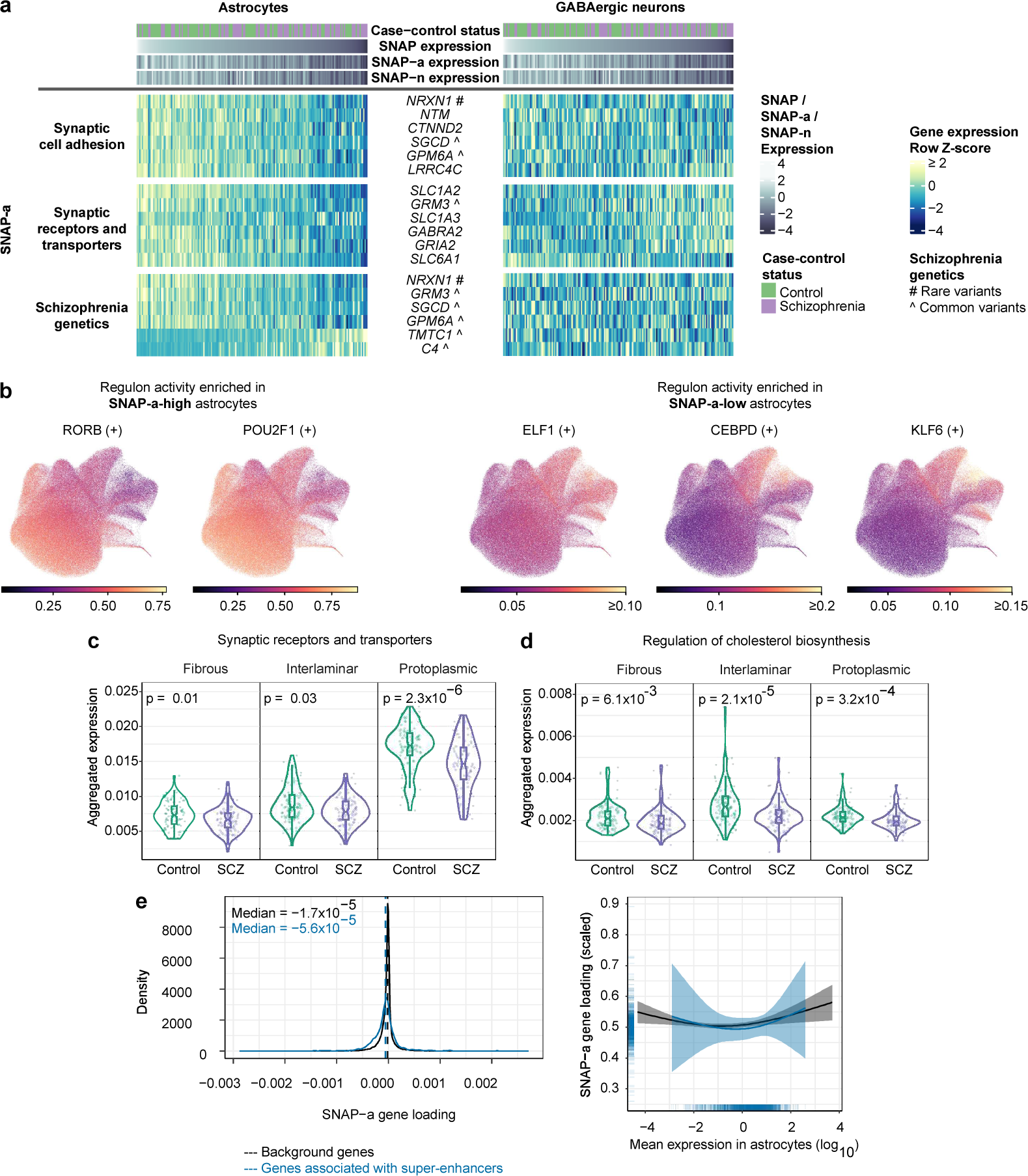
Astrocyte gene-expression programs underlying SNAP-a. **a,** See also **Fig. 3k**. Concerted expression in (left) astrocytes and (right) GABAergic neurons of genes strongly recruited by SNAP-a. These were enriched in genes encoding synaptic-adhesion proteins, intrinsic components of synaptic membranes such as transporters and receptors, as well as genes strongly implicated in human genetic studies of schizophrenia. Genes in the “Schizophrenia genetics” heatmap are from among the prioritized genes from ^23^ (FDR < 0.05) or^22^. Genes annotated by ^ are from among all genes at loci implicated by common variants in ^22^, regardless of prioritization status. **b,** UMAP of regulon activity scores (as inferred by pySCENIC ^98^) from 179,764 astrocyte nuclei from 180 donors, using the same projection from **Fig. 3a-c**. Regulons plotted are the most strongly enriched in astrocytes with high versus low SNAP-a expression. (+) indicates that the targets of the indicated regulon are predicted to be upregulated in expression. **c-d,** Transcriptional investments (by donor, separated by schizophrenia case-control status) in **(c)** genes encoding synaptic receptors and transporters and **(d)** cholesterol biosynthesis genes, in subtypes of astrocytes. Quantities plotted are the fraction of all UMIs in each subtype that are derived from these genes. P-values from a two-sided Wilcoxon rank-sum test comparing the affected to the unaffected donors are reported at the top of each panel. Box plots show interquartile ranges; whiskers, 1.5x the interquartile interval; central lines, medians; notches, confidence intervals around medians. **e,** Relationship of SNAP-a expression to association with super-enhancers. Genes expressed in astrocytes were grouped based on whether their promoters were predicted to contact super-enhancers in astrocytes (using bulk H3K27ac HiChIP and scATAC-seq data from ^99^), and SNAP-a loadings were compared between the two groups. (Left) Distributions of SNAP-a gene loadings for (blue) 1,286 genes whose promoters are predicted to contact super-enhancers in astrocytes and (black) the set of 32,325 remaining expressed background genes. (Right) Binomial smooth results of scaled SNAP-a gene loadings versus log_10_-scaled mean expression values in astrocytes, shown separately for the two groups. Shaded regions represent 95% confidence intervals.

**Extended Data Figure 25.**
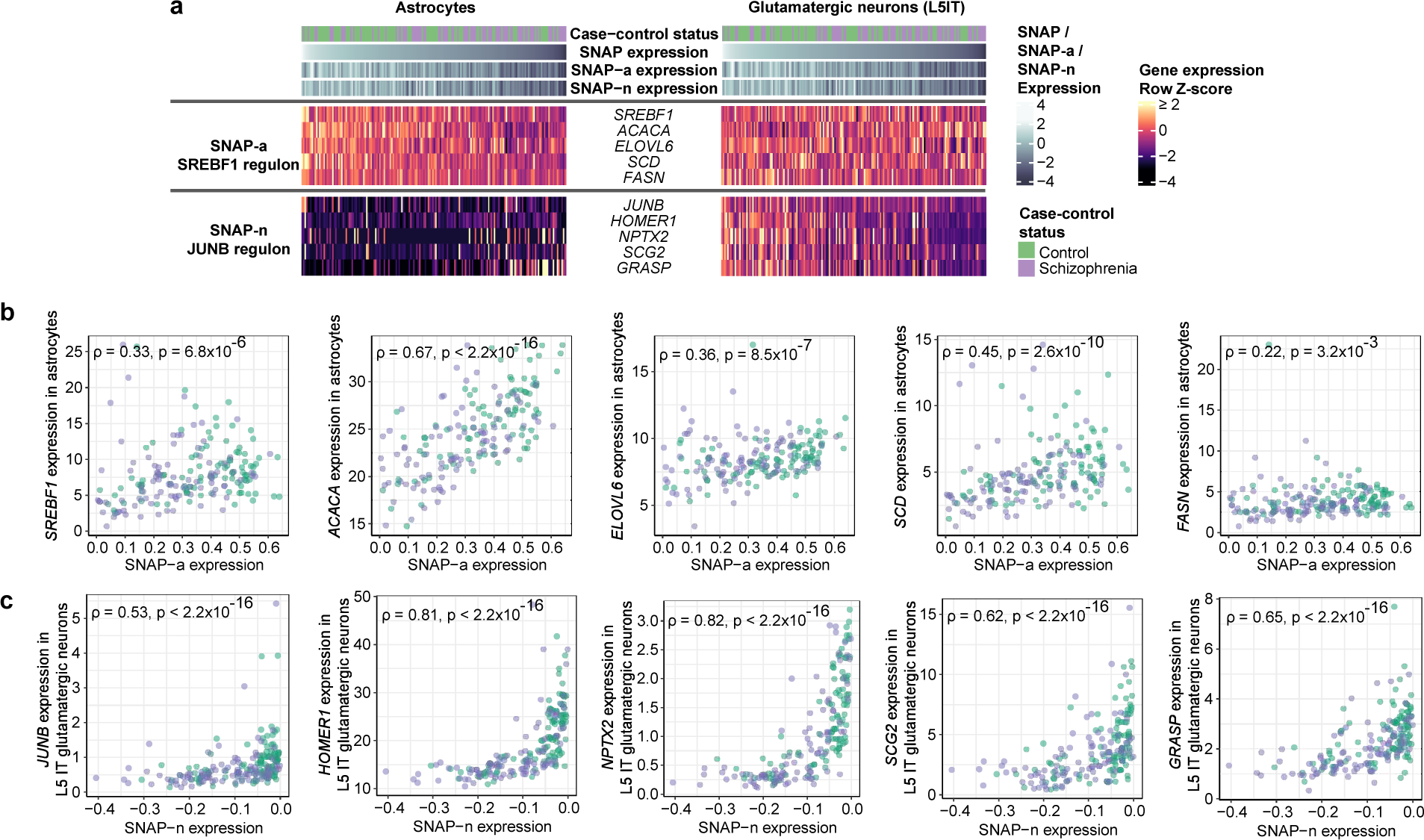
Expression of well-characterized transcriptional programs in SNAP-a and SNAP-n. **a,** Concerted expression in (left) astrocytes and (right) L5 IT glutamatergic neurons of target genes of known transcriptional programs specifically active in SNAP-a or SNAP-n. Genes are listed in decreasing order by their importance for each regulon as scored by pySCENIC. **b,** Relationship of donors’ expression levels of known SREBP1 target genes (involved in fatty acid biosynthesis) ^18,182,183^ to SNAP-a expression (Spearman’s ⍴). Target-gene expression levels in astrocytes are shown. **c,** Relationship of donors’ expression levels of known JUNB target genes (that are late-response genes) ^19,20,184^ to SNAP-n expression (Spearman’s ⍴). Target-gene expression levels in L5 IT glutamatergic neurons are shown.

**Extended Data Figure 26.**
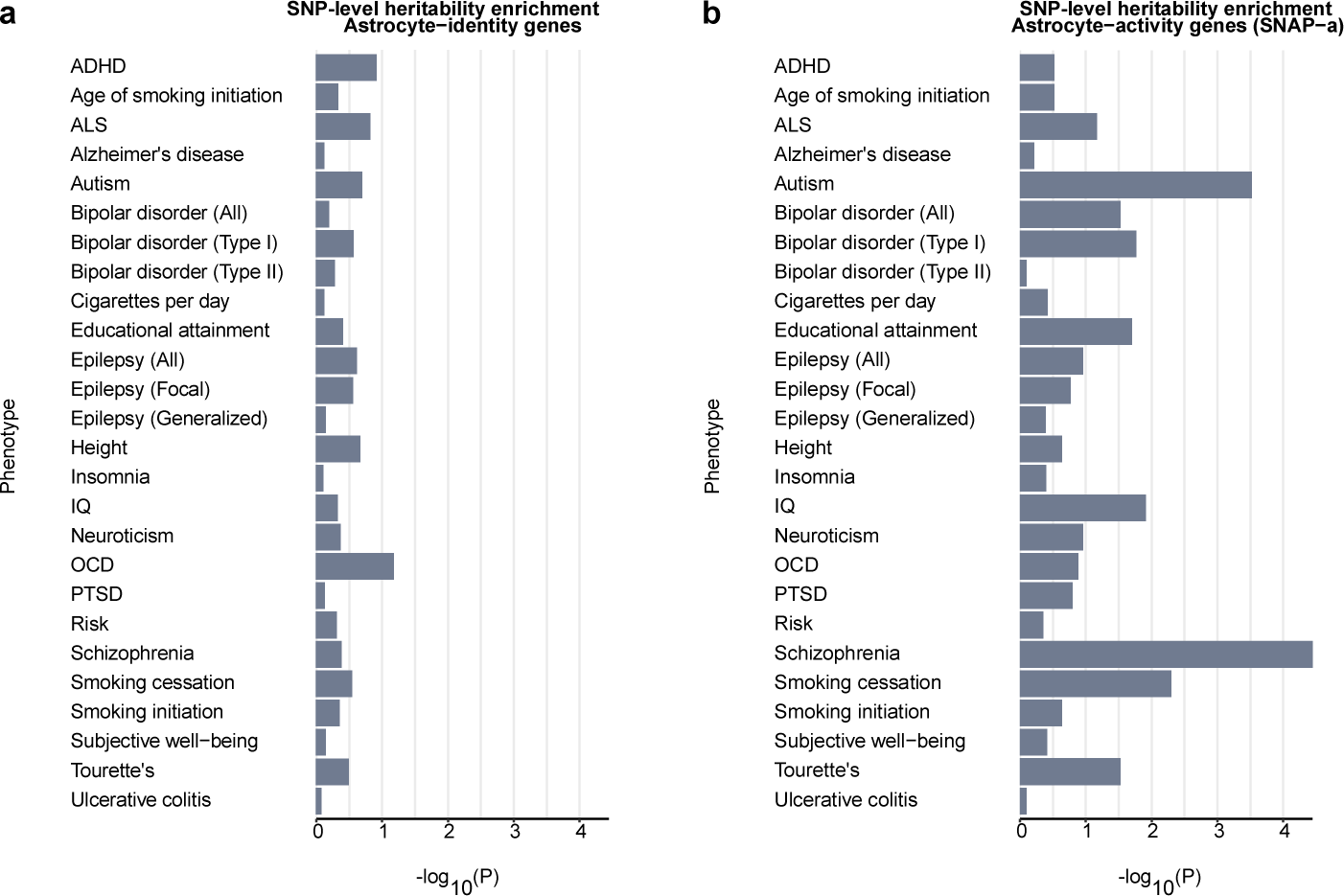
Heritability enrichment for 26 traits among the top 2,000 astrocyte-identity or astrocyte-activity (SNAP-a) genes. Heritability enrichment analysis for the indicated phenotypes in regions surrounding **(a)** the 2,000 genes most preferentially expressed in astrocytes compared to other cell types or **(b)** the 2,000 genes most strongly recruited by SNAP-a in astrocytes. Summary statistics are from the following studies: ADHD ^112^, age of smoking initiation ^115^, ALS ^113^, Alzheimer’s disease ^114^, autism^116^, bipolar disorder (all, type I, and type II) ^117^, cigarettes per day ^115^, educational attainment ^118^, epilepsy (all, focal, generalized) ^119^, height ^120^, insomnia ^122^, IQ ^121^, neuroticism ^123^, OCD ^124^, PTSD ^125^, risk ^126^, schizophrenia ^22^, smoking cessation ^115^, smoking initiation ^115^, subjective well-being ^127^, Tourette’s ^128^, ulcerative colitis ^129^.

**Extended Data Figure 27.**
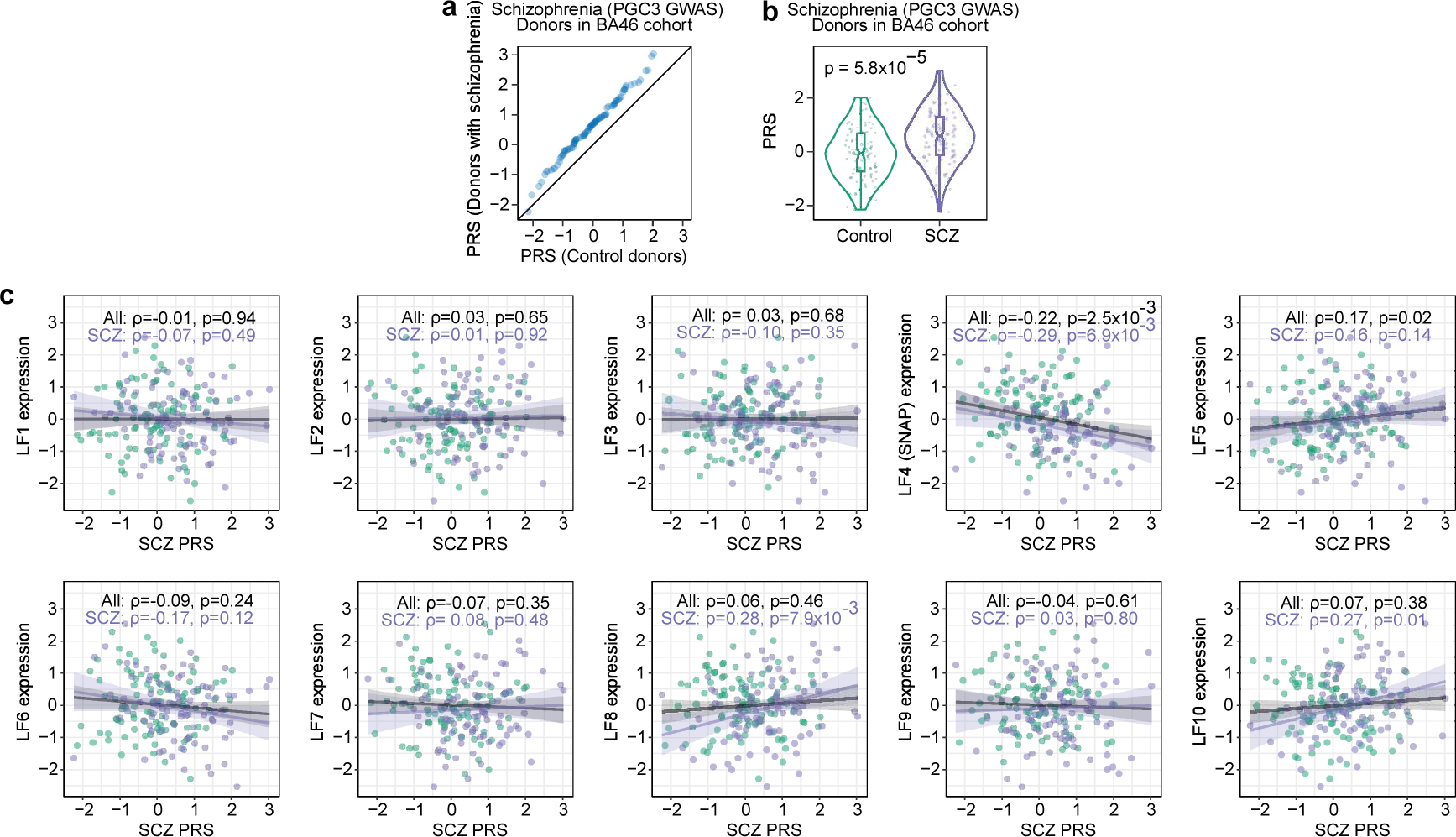
Calculation of polygenic risk scores for schizophrenia. **a,** Association of polygenic risk scores (PRS) for schizophrenia (from PGC3 GWAS, ^22^) with schizophrenia case-control status, displayed as a quantile-quantile plot that compares PRS of control donors to the PRS of donors with schizophrenia (*n* = 191 donors). **b,** Distributions of schizophrenia PRS for 94 schizophrenia cases and 97 controls. P-value is from a two-sided Wilcoxon rank-sum test. Box plots show interquartile ranges; whiskers, 1.5x the interquartile interval; central lines, medians; notches, confidence intervals around medians. **c,** See also **Fig. 4b**. Relationship of inter-individual variation in expression of each of the 10 latent factors inferred by PEER (donor scores, quantile-normalized) to donors’ polygenic risk scores (PRSs) for schizophrenia (Spearman’s ⍴; PGC3 GWAS from ^22^). Shaded regions represent 95% confidence intervals. The observed relationship of schizophrenia PRS to expression of LF4 – which associates with schizophrenia and aging – is consistent with previous observations that a PRS for schizophrenia also associates with decreased measures of cognition in older individuals ^48^ and with psychosis in Alzheimer’s Disease ^185^.

**Extended Data Figure 28.**
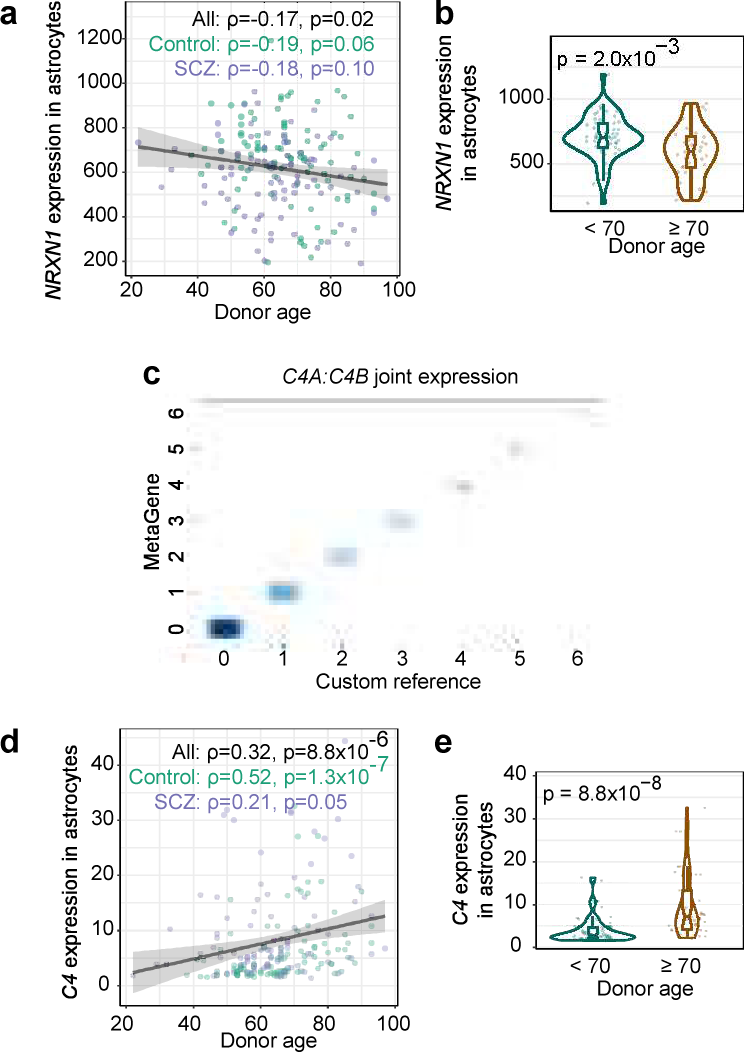
Relationship of astrocytic *NRXN1* and *C4* expression to advancing age. **a,** Relationship of *NRXN1* expression to age in astrocytes (Spearman’s represents 95% confidence interval. ⍴). Shaded region **b,** Expression of *NRXN1* in astrocytes in control donors, split by donor age (*n* = 56 donors younger than 70 years old and 37 donors 70 years old or older). P-value is from a two-sided Wilcoxon rank-sum test. Box plots show interquartile ranges; whiskers, 1.5x the interquartile interval; central lines, medians; notches, confidence intervals around medians. **c,** Validation of a metagene computational approach for identifying RNA transcripts (UMIs) from the *C4* genes. Standard analysis approaches tend to discard sequence reads from *C4A* or *C4B* because these genes are almost identical in sequence, differing only at a few key positions (far from the 3’ end), such that most reads are discarded due to low mapping quality. To measure expression of these genes, UMIs were either aligned to a custom reference genome that contained only one *C4* gene (x-axis) or were processed through a custom pipeline that identified UMIs associated with sets of gene families with high sequence homology, including *C4A/C4B* (y-axis). The two approaches (custom reference approach and joint expression of *C4A/C4B* via the metagene approach) arrived at concordant *C4* UMI counts in 15,664 of 15,669 cells tested. Note that these measurements do not distinguish between *C4A* and *C4B*. **d,** Relationship of *C4* expression to age in astrocytes (Spearman’s represents 95% confidence interval. ⍴). Shaded region **e,** Expression of *C4* in astrocytes in control donors, split by donor age (*n* = 56 donors younger than 70 years old and 37 donors 70 years old or older). P-value is from a two-sided Wilcoxon rank-sum test. Box plots show interquartile ranges; whiskers, 1.5x the interquartile interval; central lines, medians; notches, confidence intervals around medians.

